# Several independent adaptations of archaea to hypersaline environments

**DOI:** 10.1101/2023.07.03.547478

**Authors:** Brittany A. Baker, Ana Gutiérrez-Preciado, Álvaro Rodríguez del Río, Charley G. P. McCarthy, Purificación López-García, Jaime Huerta-Cepas, Edward Susko, Andrew J. Roger, Laura Eme, David Moreira

## Abstract

Several archaeal lineages thrive in high, saturating salt concentrations. These extremely halophilic archaea, including Halobacteria, Nanohaloarchaeota, Methanonatronarchaeia, and Haloplasmatales, must maintain osmotic equilibrium with their environment. For this, they use a ‘salt-in’ strategy, which involves pumping molar concentrations of potassium into the cells, which, in turn, has led to extensive proteome-wide modifications to prevent protein aggregation. However, the evolutionary history underlying these adaptations remains poorly understood. In particular, the number of times that these dramatic proteome-sweeping changes occurred is unclear due to the conflicting phylogenetic positions found for several of these lineages. Here, we present a resolved phylogeny of extremely halophilic archaea obtained using improved taxon sampling and state-of-the-art phylogenetic approaches designed to cope with the strong compositional biases of their proteomes. We describe two new uncultured lineages, Afararchaeaceae and Asboarchaeaceae, which break the long branches at the base of Haloarchaea and Nanohaloarchaeota, respectively. Our extensive phylogenomic analyses show that at least four independent adaptations to extreme halophily occurred during archaeal evolution. Finally, gene-tree/species-tree reconciliation suggests that gene duplication and horizontal gene transfer played an important role in this process, for example, by spreading key genes (such as those encoding potassium transporters) across the various extremely halophilic lineages.

## Main

For many decades, all known extremely halophilic archaea (growing at salt concentrations >30% w/v) were found to belong to the single class Halobacteria (henceforth: Haloarchaea)^1^. They dominate most hypersaline environments, from salterns and soda lakes to fermented foods^2^. However, recent technological advances, notably the ability to reconstruct metagenome-assembled genomes (MAGs), allowed the identification of several additional groups: i) Nanohaloarchaeota, nano-sized symbiotic archaea^3–5^, ii) Methanonatronarchaeia, a new class of extremely halophilic methanogens^6^, and iii) Haloplasmatales, a new order within Thermoplasmatota^7^. While Haloplasmatales have been robustly placed within Thermoplasmatota, the exact phylogenetic position of Nanohaloarchaeota and Methanonatronarchaeia remains debated^4,6,8–12^ (Extended Data Fig. 1). Initial studies found the Nanohaloarchaeota to branch as sister to Haloarchaea within the Euryarchaeota^4^, suggesting a single adaptation to extreme halophily in an ancestor common to both groups. However, subsequent studies instead supported the inclusion of Nanohaloarchaeota within the DPANN super-group^13^ (named after its five original phyla: Diapherotrites, Parvarchaeota, Aenigmarchaeota, Nanoarchaeota, and Nanohaloarchaeota) (Extended Data Fig. 1). This placement, far from Haloarchaea and Haloplasmatales, suggested an independent adaptation to hypersaline environments. Yet, the monophyly of DPANN has been questioned as it could be due to a long-branch attraction (LBA) artifact, a well-described phylogenetic artifact that could be caused by their shared fast evolutionary rates^9,11,14^.

Moderately halophilic methanogens have been known for a long time^1^, but true extremely halophilic ones, the Methanonatronarchaeia, have only been recently characterized^6^. Being placed as a sister group to the Haloarchaea, they were proposed to be an “evolutionary intermediate” between them and Class II methanogens^6^. However, more recent studies favored a much deeper position within the Euryarchaeota, at the base of the superclass Methanotecta^8–10,12^ (Extended Data Fig. 1). Alternatively, the robust placement of hikarchaea (family ‘UBA12382’ in the latest release (r214) of the Genome Taxonomy Database GTDB^15^), a lineage of non-halophilic deep-ocean archaea originally named Marine Group IV^16^, as sister-group to the Haloarchaea supported a relatively recent adaptation to halophily in the latter^10^ (Extended Data Fig. 1).

All extremely halophilic archaea have experienced radical physiological and genomic changes to deal with the high environmental osmotic stress. They pump high concentrations (up to ∼4M) of potassium into their cells^17^ and prevent protein aggregation that would be caused by this high intracellular ionic concentration thanks to their much more acidic proteome than that of non-halophilic archaea. Specifically, haloarchaeal proteomes exhibit a massive enrichment in acidic amino acids (aspartic (D) and glutamic (E) acid) and a depletion in basic and large hydrophobic amino acids (such as isoleucine (I) and lysine (K))^18–21^. How and how many times these adaptations occurred are still unresolved questions because of the uncertain phylogenetic position of the different halophilic archaeal lineages. This difficulty in confidently resolving their phylogeny mostly stems from the combined presence of halophile-specific amino acid biases, which are poorly modeled during standard phylogenetic reconstructions, and the long branches shown by several extremely halophilic archaeal lineages. Altogether, this severely limits our comprehension of the evolutionary trajectories of these organisms to adapt to their extreme habitats.

Here, we identify and describe two new families of extremely halophilic archaea, the Afararchaeaceae and Asboarchaeaceae, which break the long branches at the base of Haloarchaea and Nanohaloarchaeota, respectively. With this enhanced taxon sampling and improved phylogenetic methods, we obtained a fully resolved phylogeny of all halophilic archaea. We propose an updated evolutionary scenario that involves at least four independent adaptations to hypersaline environments and an important role of horizontal gene transfer (HGT) between the various groups of halophilic archaea.

## Results and Discussion

### Characterization of two new groups of extremely halophilic archaea

The Danakil epression (Afar region, Ethiopia) contains several hypersaline lakes largely dominated by extremely halophilic archaea (Belilla et al., 2019, 2021). Among the metagenome-assembled genomes (MAGs) reconstructed from these lakes^22^, we identified 13 belonging to two new lineages of extreme halophiles (Fig. 1a,c and Supplementary Table 1), plus one additional MAG (DAL-9Gt_70_90C3R) placed as the deepest-branching member of the Haloarchaea in a phylogenomic tree of Archaea (Fig. 1a and Supplementary Table 1).

**Fig. 1.**
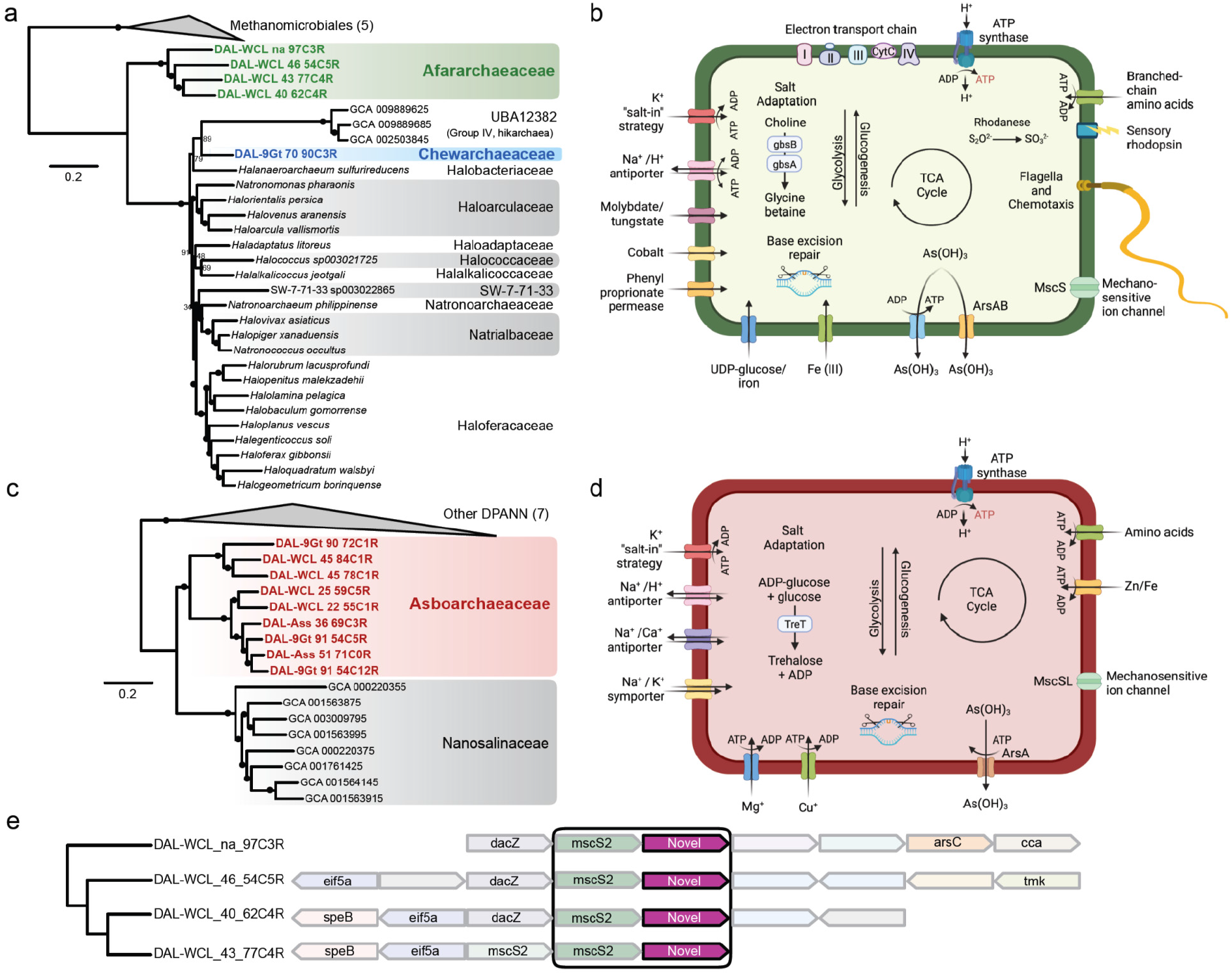
Phylogenetic position and metabolic potential of the new families Afararchaeaceae and Asboarchaeaceae. **(a)** Maximum likelihood phylogenetic tree of 35 euryarchaea, including the four new Afararchaeaceae MAGs (highlighted in green), based on the concatenation of 122 single-copy proteins obtained from the Genome Taxonomy Database (GTDB). The tree was inferred by IQ-TREE with the LG+C60+F+G4 model of sequence evolution. The statistical support for branches, with filled circles representing values equal to or larger than 99% support, corresponds to 1,000 ultra-fast bootstrap replicates. Scale bar indicates the expected average number of substitutions per site. All taxonomic ranks shown are based on the GTDB r207 family-level classification. See Supplementary Fig. 1 for the uncollapsed tree. **(b)** Non-exhaustive metabolic scheme based on the predicted gene content of the most complete afararchaeal MAG (DAL-WCL_na_97C3R). A detailed table of the predicted gene content can be found in Supplementary Data 1. **(c)** Maximum likelihood phylogenetic tree of 24 DPANN archaea, including the nine new Asboarchaeaceae MAGs (highlighted in salmon), based on the concatenation of 99 single-copy proteins obtained from GTDB. The tree was inferred by IQ-TREE with the LG+C60+F+G4 model of sequence evolution. The statistical support for branches corresponds to 1,000 ultra-fast bootstrap replicates. The scale bar indicates the expected average number of substitutions per site. All taxonomic ranks are based on the GTDB r207 family-level classification. See Supplementary Fig. 2 for the uncollapsed tree. **(d)** Non-exhaustive metabolic scheme based on the predicted gene content of the most complete asboarchaeal MAG (DAL-WCL_45_84C1R). A detailed table of the predicted gene content can be found in Supplementary Data 2. **(e)** Gene maps showing a novel gene family (magenta) linked to a conserved mechanosensitive ion channel (mscS2) in the afararchaeal MAGs. Gene abbreviations are as follows: agmatinase (speB), eukaryotic initiation factor 5A (eif5a), di-adenylate cyclase (dacZ), arsenate reductase (arsC), tRNA nucleotidyltransferase (cca), thymidylate kinase (tmk).

The first group – a novel family-level lineage that we have named Afararchaeaceae, for the Afar region in Ethiopia – was represented by four moderately GC-rich (53-60%) MAGs with average nucleotide identity (ANI) values between 72 and 74% among them. Afararchaeaceae were placed with maximal statistical support as a new sister lineage to the group UBA12382 (or ‘hikarchaea’^10^)+Haloarchaea (Fig. 1a). This position breaks the long branch between Methanomicrobiales (methanogenic Euryarchaeota) and hikarchaea+Haloarchaea, suggesting a secondary adaptation of hikarchaea to low salinity from an extremely halophilic ancestor. The most complete afararchaeal MAG (DAL-WCL_na_97C3R), which we propose the name *Afararchaeum irisae* gen. nov., sp. nov. (see species description below), had a genome size of ∼1.9 Mbp (Supplementary Table 1). KEGG annotation^23^ of *A. irisae* indicates that the afararchaeal genomes likely encode aerobic heterotrophic metabolic pathways. These organisms are likely to utilize branched-chain amino acids as a carbon source, similar to many known Haloarchaea^24^ (Fig. 1b, Supplementary Data 1). The Afararchaeaceae appear to be mobile, possessing all genes for the archaeal flagellum (archaellum)^25^ and an operon involved in chemotaxis. Additionally, all four afararchaeal MAGs encode a single type-II sensory (SRII) rhodopsin, a membrane protein able to generate a phototaxis signal^26^. However, we did not identify any bacteriorhodopsin genes in these MAGs, suggesting that these archaea do not use light as an extra energy source as many Haloarchaea do^27^. As expected, the Afararchaeaceae most likely employ a salt-in osmoregulation strategy involving multiple K^+^ transporters (eight Trk-like and two Kef-like), mechanosensitive ion channels (MscS and MscL), and Na^+^/Ca^2+^ exchangers (Supplementary Data 1). They consequently also exhibit a highly acidic proteome (Fig. 2a,b).

**Fig. 2.**
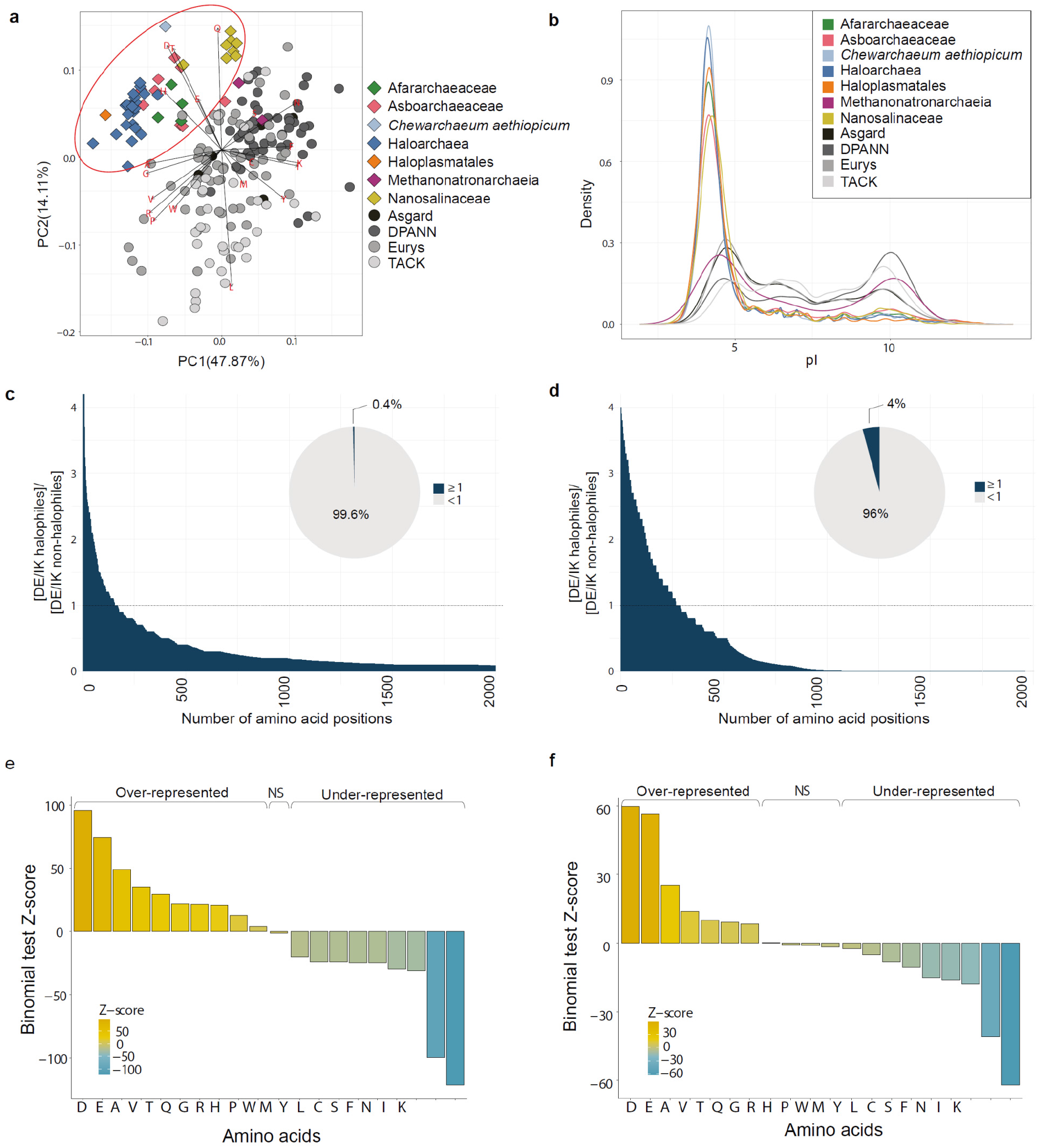
Protein amino acid compositional biases in extremely halophilic archaeal lineages. **(a)** PCA plot of 192 archaeal proteomes based on amino acid frequencies. The red ellipse indicates the clustering of all extreme halophiles (colored diamonds), including the newly identified families Afararchaeaceae (green) and Asboarchaeaceae (salmon color). **(b)** Isoelectric point (pI) distribution of 192 archaeal proteomes. Non-halophilic archaea (grey lines) display a bimodal distribution of pI values, while extreme halophiles (colored lines) exhibit a single spike at pI ∼4, indicating a highly acidic proteome. **(c**,**d)** D+E/I+K site-by-site bias (defined as the ratio [D+E/I+K for halophiles]/[D+E/I+K for non-halophiles]) for the 2,000 most biased sites of the **(c)** NM dataset (39,385 amino acid positions) and **(d)** RP dataset (6,792 amino acid positions). Inset pie charts depict the proportion of amino acids with a ratio greater than or equal to 1 (dark blue) versus less than 1 (grey). **(e**,**f)** Binomial tests for the **(e)** NM and **(f)** RP datasets comparing the proportions of all 20 amino acids between extreme halophiles and non-halophiles. Z-scores were calculated relative to extreme halophiles, with |Z| > 1.96 indicating significant enrichment of a given amino acid in extreme halophile sequences (“Over-represented”), |Z| < -1.96 indicating significant depletion of a given amino acid in extreme halophile sequences (“Under-represented”), and some amino acids showing no significant bias (“NS”).

The second group was represented by nine MAGs with variable GC content (46-64%) with ANI values between 74 and 79% among them. They branch in the DPANN superphylum as a sister group to the family Nanosalinaceae within the Nanohaloarchaeota (Fig. 1c). They are related to MAGs from hypersaline anoxic sediments that were previously classified by Zhao et al. as the new families ‘Nanoanaerosalinaceae’ and ‘Nanohalalkaliarchaeaceae’^5^ (Supplementary Fig. 3).

However, according to the GTDB^15^ classification criteria, these two families have been merged within a single one, which has been informally named ‘JALIDP01’. Our MAGs offer a good coverage of this family, three of them being related to the former ‘Nanoanaerosalinaceae’ and six to the single MAG representing the ‘Nanohalalkaliarchaeaceae’^5^ (Supplementary Fig. 3). Given the taxonomic uncertainty on this group and that, contrary to the assumption of an anaerobic lifestyle^5^, it can also be found in oxic environments as in our case, we propose to formally name the new family Asboarchaeaceae, for ‘*asbo*’ meaning salt in the Afar language, which acknowledges that these organisms have always been found in hypersaline systems.

Due to the streamlined nature of their genomes, certain typically conserved genes are absent in DPANN, leading to an underestimation of genome completeness when evaluating based on the presence of such genes. As a result, DPANN genomes generally have a maximum estimated completeness of around 85%^14^. In this context, we likely have at least one complete (84% according to CheckM^28^) asboarchaeal MAG (DAL-WCL_45_84C1R) (Supplementary Table 1). We propose it to represent the type species for this family under the name *Asboarchaeum danakilensis* gen. nov., sp. nov. (see species description below). Its genome size of ∼1.2 Mbp is consistent with the small genomes found in other DPANN lineages^14^. Similarly to them^29–31^, Asboarchaeaceae lack many major biosynthetic pathways thought to be required for autonomous growth (e.g., lipid, nucleotide, and amino acid biosynthesis) (Fig. 1d and Supplementary Data 2), suggesting they live a symbiotic lifestyle that requires a host for survival. The Asboarchaeaceae lack a canonical electron transport chain, but they do possess all major subunits of a V/A-type ATP synthase (Fig. 1d), as previously observed^29^. We again predict that Asboarchaeaceae utilize a salt-in osmoprotective strategy pertaining to the identification of multiple K^+^ transporters (Supplementary Data 2) and their highly acidic proteome (Fig. 3a,b). Despite their relatively close phylogenetic relationship with the family Nanosalinaceae, they display a distinct amino acid composition (Fig. 3a), further supporting that they constitute a new group within the Nanohaloarchaeota.

**Fig. 3.**
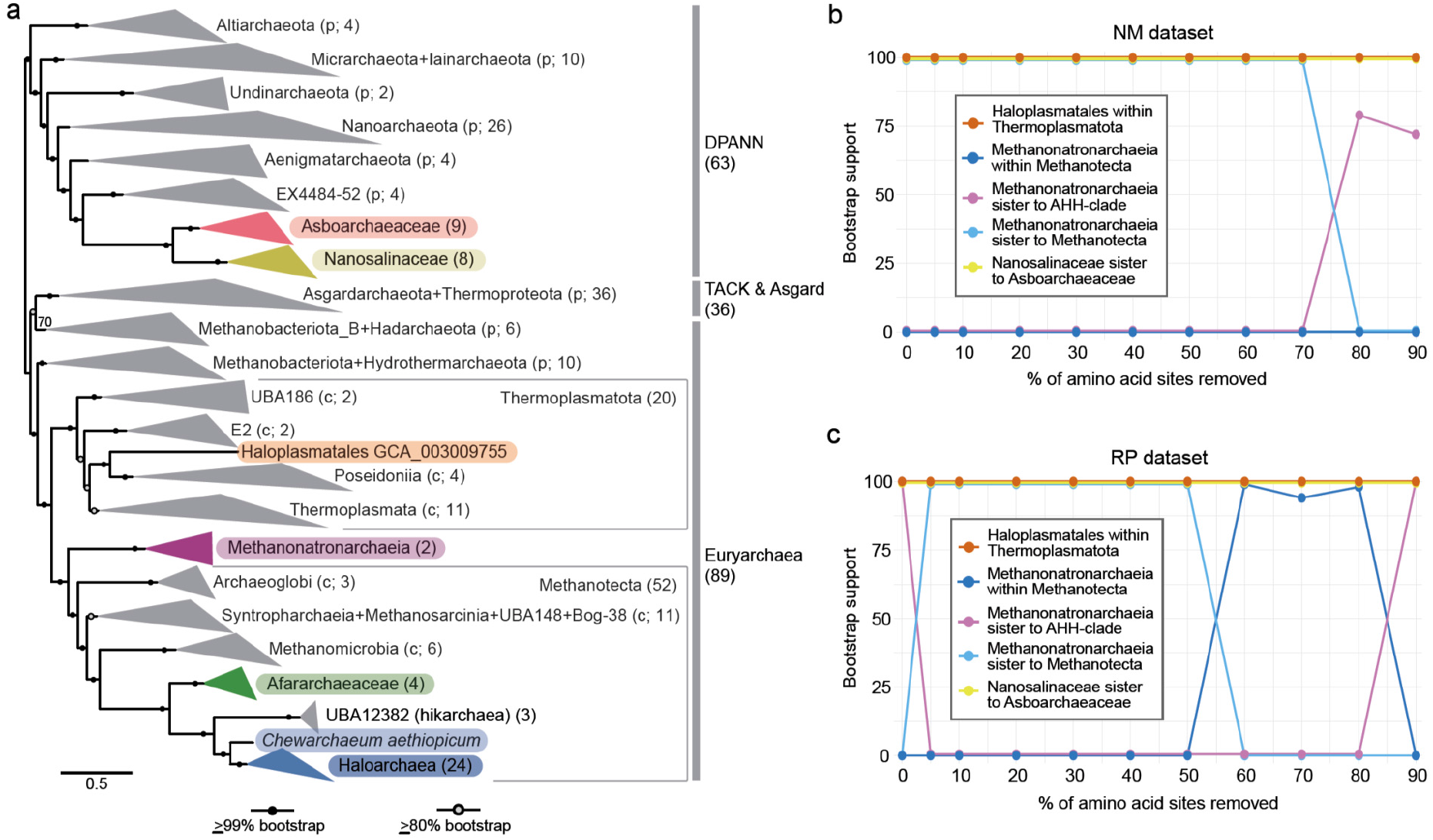
Maximum likelihood phylogeny of archaea, including the new groups Afararchaeaceae and Asboarchaeaceae. **(a)** Phylogenetic tree based on the concatenation of 136 conserved markers (NM dataset) across 192 taxa (39,385 sites) using IQ-TREE under the LG+C60+F+G4 model of evolution. Statistical support indicated on the branches corresponds to 1,000 ultra-fast bootstrap replicates. The scale bar indicates the number of substitutions per site. Colors indicate the currently known groups of extremely halophilic archaea. The size of collapsed clades is indicated in parentheses; see Extended Data Fig. 2 for the uncollapsed tree. **(b**,**c)** Impact of the progressive removal (in steps of 10%) of the most compositionally biased sites from the **(b)** 192-NM (39,385 amino acid positions) and **(c)** 192-RP (6,792 amino acid positions) datasets. Lines show the statistical support values for the position of each of the halophilic clades of interest. These support values were estimated using the ultrafast bootstrap approximation from the ML tree reconstruction (LG+C60+F+G4 model) for each site-removal step.

### Detection of novel gene families in Afararchaeaceae and Asboarchaeaceae

Functional annotation of the genes of divergent species, such as the DPANN archaea, can be difficult. We, therefore, applied a two-step procedure to characterize potential novel genes in the Afararchaeaceae and Asboarchaeaceae. First, we searched for genes present in their genomes but with no detectable homologs in sequence databases of cultured organisms (RefSeq^32^, Pfam^33^, and EggNOG^34^). Both lineages were rich in potentially novel gene families (10 to 30% of their total gene set; Extended Data Fig. 3a). Second, we mapped these against a collection of 169,529 prokaryotic genomes, which also included a large representation of non-cultured species^35^. We confirmed that only 14% of the novel families from asboarchaea and 17.1% from afararchaea have detectable homologs in other uncultured prokaryotic species, indicating that both groups encode many unknown lineage-specific genes (Supplementary Data 3 and 4). Interestingly, the isoelectric point calculated for the proteins encoded by these novel genes was clearly shifted towards acidic pH values (Extended Data Fig. 3b), consistent with adaptation to hypersaline environments^36^. 38% and 24% of the novel proteins of Afararchaeaceae and Asboarchaeaceae have predicted transmembrane domains, respectively, while 9% and 7% have detectable signal peptides. These proteins are most likely targeted to the membrane or extracellular space and interact directly with the hypersaline environment. To gain insight into the function of the novel genes, we investigated their genomic context. We found that 5% in Afararchaeaceae (Supplementary Data 3) and 18% in Asboarchaeaceae (Supplementary Data 4) are tightly coupled with specific genes of known function (i.e., next to the same gene of known function in >90% of the genomes) and thus probably perform functions related to that of their neighbors^35^. One interesting example in Afararchaeaceae is a novel protein found next to a mechanosensitive ion channel (Fig. 1e). In both prokaryotes and eukaryotes, mechanosensitive ion channels provide protection against hypo-osmotic shock^37^, suggesting this novel gene could be involved in the osmotic regulation of afararchaea.

### Identification of a conserved core of archaeal phylogenetic markers

Previous investigations of the phylogenetic placement of extreme halophiles were predominantly based on single proteins^4,9^ or on sets of concatenated ribosomal proteins^6,10^. However, these are small datasets that contain a restricted number of sites and provide limited phylogenetic information^11,38^. In addition, ribosomal proteins, due to their multiple tight interactions (with ribosomal, messenger, and transfer RNAs, and with other ribosomal proteins), can exhibit compositional biases different from the rest of the proteome, which can be amplified when they are concatenated together^11,39^. To overcome these potential biases and to robustly pinpoint the phylogenetic positions of extremely halophilic archaea, we performed in-depth phylogenomic analyses of a dataset of 136 new marker proteins (NM dataset; 39,385 amino acid positions) highly conserved among archaea^11^. Proteins in this dataset display a wide diversity of functions (Supplementary Data 5), which can help minimize phylogenetic artifacts rising from biases linked to co-evolution patterns. We manually curated each individual protein phylogeny to ensure the NM dataset contained no evidence of HGT or hidden paralogy (see Methods). Additionally, we curated a set of 48 ribosomal proteins (RP dataset, 6,792 amino acid positions) to compare their phylogenetic signal with that of the NM dataset.

### Testing the influence of taxon sampling

Some extremely halophilic archaea display long branches in phylogenetic trees (Extended Data Fig. 2) such that their phylogeny might be affected by LBA artifacts^40,41^. To investigate this possibility, we analyzed two additional datasets in addition to the full dataset of 192 taxa representing the four major archaeal supergroups (DPANN, TACK (Thermoproteota according to GTDB), Asgardarchaeota, and Euryarchaeota) (Fig. 3a, Extended Data Fig. 2): (1) a dataset focusing on Euryarchaeota, including the newly discovered Afararchaeaceae (87 taxa: 87-NM and 87-RP for new markers and ribosomal proteins, respectively) (Supplementary Figs. 4 and 5, Supplementary Data 6), and (2) a dataset consisting of the 87 Euryarchaeota plus 17 Nanohaloarchaeota (8 Nanosalinaceae + 9 Asboarchaeaceae) (104 taxa: 104-NM and 104-RP; Supplementary Figs. 6 and 7, Supplementary Data 6). The positions of all the clades of extremely halophilic archaea – except Methanonatronarchaeia – were congruent across maximum likelihood (ML) phylogenies reconstructed from these various taxon samplings and marker sets. By contrast, we observed two different yet highly supported placements of Methanonatronarchaeia (Supplementary Figs. 4-7). All NM-based ML trees (87-NM, 104-NM, and 192-NM) and the 104-RP dataset placed them sister to the Methanotecta (i.e., Haloarchaea, ‘hikarchaea’, Class II methanogens, Methanopagales, ANME-1, Synthrophoarchaeales, and Archaeoglobales) (Extended Data Fig. 2, Supplementary Figs. 4, 6 and 7) whereas two RP-based ML trees (87-RP and 192-RP) placed them sister to the Afararchaeaceae+’hikarchaea’+Haloarchaea (the ‘AHH-clade’; Supplementary Fig. 5 and Extended Data Fig. 4). These results suggested a clear influence of taxon sampling on the position of the Methanonatronarchaeia inferred from the RP dataset.

To further investigate the effect of taxon sampling but also of the sequence evolution model, we ran phylogenetic analyses of the 87 and 104 taxa sets for both NM and RP markers in a Bayesian framework (with four Markov Chain Monte Carlo (MCMC) chains for each analysis) to use the more complex, but time-consuming, CAT+GTR model. Again, conflicting placements of Methanonatronarchaeia were observed. The chains inferred from the NM datasets (Supplementary Figs. 8 and 10) supported either Methanonatronarchaeia sister to the Methanotecta (three 87-NM and one 104-NM chains) or Methanonatronarchaeia sister to the AHH-clade (one 87-NM and three 104-NM chains). This Methanonatronarchaeia-AHH sister relationship was only observed in this particular case among all our NM dataset inferences. Alternatively, two 87-RP and one 104-RP chain placed Methanonatronarchaeia within Methanotecta, while the other RP chains supported two other placements: Methanonatronarchaeia sister to the AHH-clade (two 87-RP and one 104-RP chains) or Methanonatronarchaeia branching with all other extremely halophilic archaea (except Haloplasmatales) within the Euryarchaeota (two 104-RP chains) (Supplementary Figs. 9 and 11). The latter was the only time we observed the Nanosalinaceae+Asboarchaeaceae elsewhere than nested within DPANN or sister to all Euryarchaeota. These results indicate at least three conflicting signals in the RP dataset.

These analyses again showed that the phylogenetic placement of extreme halophiles, especially the Methanonatronarchaeia, is sensitive to taxon sampling, model selection, and phylogenetic framework. The strong compositional biases linked to the ‘salt-in’ osmotic strategy of extremely halophilic archaea can be the reason that makes it difficult to accurately place them in phylogenies using standard substitution models^42^.

### Addressing the effect of compositional biases

Model misspecification induced by compositional bias is a known source of tree reconstruction artifacts. To mitigate this, we aimed to identify significantly differently represented amino acids in halophiles versus non-halophiles in the 192 taxa NM and RP datasets and calculate, for each amino acid, the Z-score from a binomial test of two proportions (see Methods). D+E and I+K were the most under and over-represented in extreme halophiles, respectively (Fig. 2e,f). To limit potential LBA artifacts on the phylogeny of extreme halophiles, previous studies have focused on either recoding data from 20 to 4 character states10,43 or removing the fastest-evolving sites from the sequence alignments8,10,43. However, the latter required removing up to 50% of the alignment sites before a change in the tree topologies was observed, particularly for the Methanonatronarchaeia8,10. This significantly reduces the amount of phylogenetic signal left, which is problematic for small datasets such as the RP-based ones12. Therefore, we tested two alternative approaches to alleviate the halophile-specific compositional biases while maintaining a high amount of phylogenetic information.

First, we implemented the GFmix modeling framework^42^ to the specific compositional biases of halophilic archaea. GFmix is a site-heterogeneous mixture model that adjusts amino acid frequencies in each class of the mixture model in a branch-specific manner to accommodate overall shifts in amino acid composition over the branch. To accomplish this, amino acids are categorized into three groups: those that increase in frequency on the branch (i.e., become over-represented in descendant taxa), decrease (i.e., become under-represented), and remain unchanged (i.e., ‘other’).

We used the mixture model LG+C60+F+G4 with GFmix, setting [D,E]/[I,K] as the compositional ratio varying over branches (henceforth referred to as GFmix-DE/IK model) and calculated the likelihood of four different tree topologies under this complex model: i) Nanosalinaceae+Asboarchaeaceae within DPANN and Methanonatronarchaeia sister to the AHH-clade; ii) Nanosalinaceae+Asboarchaeaceae within DPANN and Methanonatronarchaeia deep within Euryarchaea; iii) monophyly of the AHH-clade, Methanonatronarchaeia, and Nanosalinaceae+Asboarchaeaceae, with Methanonatronarchaeia as the deepest branch, and iv) monophyly of the AHH-clade, Methanonatronarchaeia, and Nanosalinaceae+Asboarchaeaceae, with Nanosalinaceae+Asboarchaeaceae as the deepest branch (Extended Data Fig. 6). In all cases, the LG+C60+F+G4+GFmix-DE/IK model yielded better likelihood values than the classical LG+C60+F+G4 model alone. The highest-scoring topologies for both datasets placed Nanosalinaceae+Asboarchaeaceae within DPANN. However, despite the improvement in fit to the data by LG+C60+F+G4+GFmix-DE/IK model, this approach still produced incongruent results between the RP and NM datasets for the position of Methanonatronarchaeia, with the 192-RP dataset supporting them as sister to the AHH-clade and the 192-NM as sister to Methanotecta (Extended Data Fig. 6). We also tested a GFmix variant with larger groups of over- and under-represented amino acids based on all of the statistically significant (|Z|>1.96) over-represented and under-represented amino acids from the binomial test of the NM dataset (see Fig. 2e). Although this yielded an even better fit (Extended Data Fig. 6), the relative preferences of topologies for each dataset did not change.

We, therefore, applied a second strategy based on the progressive removal of the most compositionally biased alignment sites. We calculated the site-by-site D+E/I+K ratio for the halophilic lineages and divided it by the same ratio for the non-halophilic lineages (Extended Data Fig. 5). We then ranked the alignment sites from the highest to the lowest ratio (Fig. 2c,d) and then removed sites in 10% increments for the 192 taxa NM and RP datasets (Fig. 3b,c). For the 192-NM dataset, the position of Methanonatronarchaeia remained consistent with the topology inferred from the full dataset up until 80% of sites were removed; they then branched with low support (bootstrap <80%) as sister group of the AHH-clade (Fig. 3b). By contrast, for the 192-RP dataset, Methanonatronarchaeia shifted to a sister position to Methanotecta after removing only 5% of the most biased alignment sites (Fig. 3c). This suggested that, while the NM dataset does contain sites with biased D+E/I+K ratio between halophiles and non-halophiles (Fig. 2c), the impact of these sites (only 0.4% of 39,385 amino acid positions with a ratio ≥1) has less of an influence when compared to the ten times more numerous highly biased sites in the RP dataset (4% of 6,792 amino acid positions with a ratio ≥1; Fig. 2d).

We examined the ribosomal proteins containing the most biased sites and found that nearly all of these sites were located in proteins exposed on the external surface of the ribosomal complex (e.g., L1, L12e, S6, and S15; Supplementary Fig. 12). These proteins, therefore, are in close interaction with the K^+^-rich cytoplasm. To confirm further the impact of the D+E/I+K bias on the RP-based phylogeny, we inferred a ML tree using a concatenation of the 18 most biased ribosomal proteins. We observed all extremely halophilic groups clustering with 100% ultrafast bootstrap support (Supplementary Fig. 13). We also inferred Bayesian phylogenies (CAT+GTR model; four MCMC chains) based on the 104-NM and 104-RP datasets with 20% of the most biased alignment sites removed. Contrary to the trees reconstructed with the untreated datasets (see above), all chains for both datasets yielded a topology with full support for the deeper-branching position of Methanonatronarchaeia sister to Methanotecta (Supplementary Figs. 14 and 15).

A recent study using the ATP synthase subunits A and B for phylogenetic inference concluded that Nanohaloarchaeota placed sister to Haloarchaea within the Euryarchaeota^9^. The authors suggested that these proteins are less susceptible to phylogenetic reconstruction artifacts, such as LBA, due to two reasons: i) their slow evolutionary rate and ii) their belonging to a single complex, which is expected to have a more consistent phylogenetic signal compared to larger protein sets that may reflect different evolutionary histories. Alternatively, this Nanohaloarchaeota+Haloarchaea relationship has been explained as a case of HGT from Haloarchaea to their nanohaloarchaeal symbionts^44^. However, when we removed 15% of the sites with the highest D+E/I+K ratio from the ATP synthase dataset (150 sites), Nanohaloarchaea moved to a deeper-branching position, no longer sister to Haloarchaea (Extended Data Fig. 7). Again, this suggested that only a few but highly biased alignment sites artificially drove the extreme halophilic archaeal lineages to branch together.

In conclusion, our phylogenetic analyses, especially those mitigating the strong convergent compositional bias shared by the different halophilic lineages, robustly support that adaptation to extreme halophily occurred at least four times independently in archaea: in the AHH-clade, in Methanonatronarchaeia, in Haloplasmatales, and in Nanosalinaceae+Asboarchaeaceae, respectively.

### Tree-aware reconstruction of gene content evolution in archaeal extreme halophiles

To investigate the gene content evolution underlying these adaptations, we applied the amalgamated likelihood estimation (ALE) method^45^ to 17,288 orthologous proteins in the 192 taxa genomic dataset. Using this species tree-gene tree reconciliation approach, we estimated the number of gene duplications, transfers, originations, losses, and copy numbers at all ancestral nodes of the 192-NM phylogenomic tree (Fig. 3a), which we used as the species tree. In contrast with a previous analysis focused only on Methanotecta^10^, we analyzed representatives of all major archaeal lineages, including, most importantly to our study, Methanonatronarchaeia, which were excluded from previous analyses because of their unresolved phylogenetic position at that time. We observed that the main processes governing gene content in archaea, including the halophilic groups, are gene transfer and gene loss (Fig. 4 and Extended Data Fig. 8). In the case of Haloarchaea, which figure among the archaea with the largest genome sizes^46^, gene originations, and duplications have also been significant during their early evolution. This is the case for several inorganic ion transporters important for the maintenance of osmotic equilibrium in these organisms, including Trk- and Kef-type K^+^ transporters (Supplementary Figs. 16-19), Mg^2+^ transporters (Supplementary Fig. 20), SSF Na^+^/solute symporters (Supplementary Fig. 21), NhaP-type K^+^/H^+^ antiporters (Extended Data Fig. 10a), Ca^+^/Na^+^ and Na^+^/H^+^ antiporters (Supplementary Figs. 22 and 23, respectively), and also of the molecular chaperone GrpE, which participates in the response to hyperosmotic stress by preventing the aggregation of stress-denatured proteins^47^ (Supplementary Fig. 24). Amino acid transporters also show duplications in this group (Extended Data Fig. 9), which contains many species that thrive on amino acids^24^. Haloplasmatales also exhibit a relatively large number of duplications, not only in genes related to metabolism but also to informational processes such as transcription and DNA replication and repair (Extended Data Fig. 9). In Nanosalinaceae and Asboarchaeaceae, gene transfer was a dominant process, although less so than in the other halophilic groups, most likely because of the strong evolutionary constraints that operate in these nanosized archaea to keep small genome sizes^48^. The branch leading to the ‘hikarchaea’ shows a very different pattern, clearly dominated by gene loss. It supports the hypothesis that these halotolerant archaea evolved secondarily from an extremely halophilic ancestor (the Hik-Haloarchaea ancestor, with 1,416 inferred protein-coding genes, Fig. 4) during their adaptation to marine oligotrophic environments, where many prokaryotic species typically show streamlined genomes^49,50^. Nevertheless, this adaptation was also accompanied by duplications of some specific genes involved in energy production and conversion and carbohydrate and amino acid transport and metabolism, most likely also related to the adaptation to the nutrient-poor deep sea habitat (Extended Data Fig. 9). A notable example is the presence of multiple copies of the aerobic-type carbon monoxide dehydrogenase (Supplementary Fig. 25), an enzyme previously found in other microorganisms adapted to this environment^51^.

**Fig. 4.**
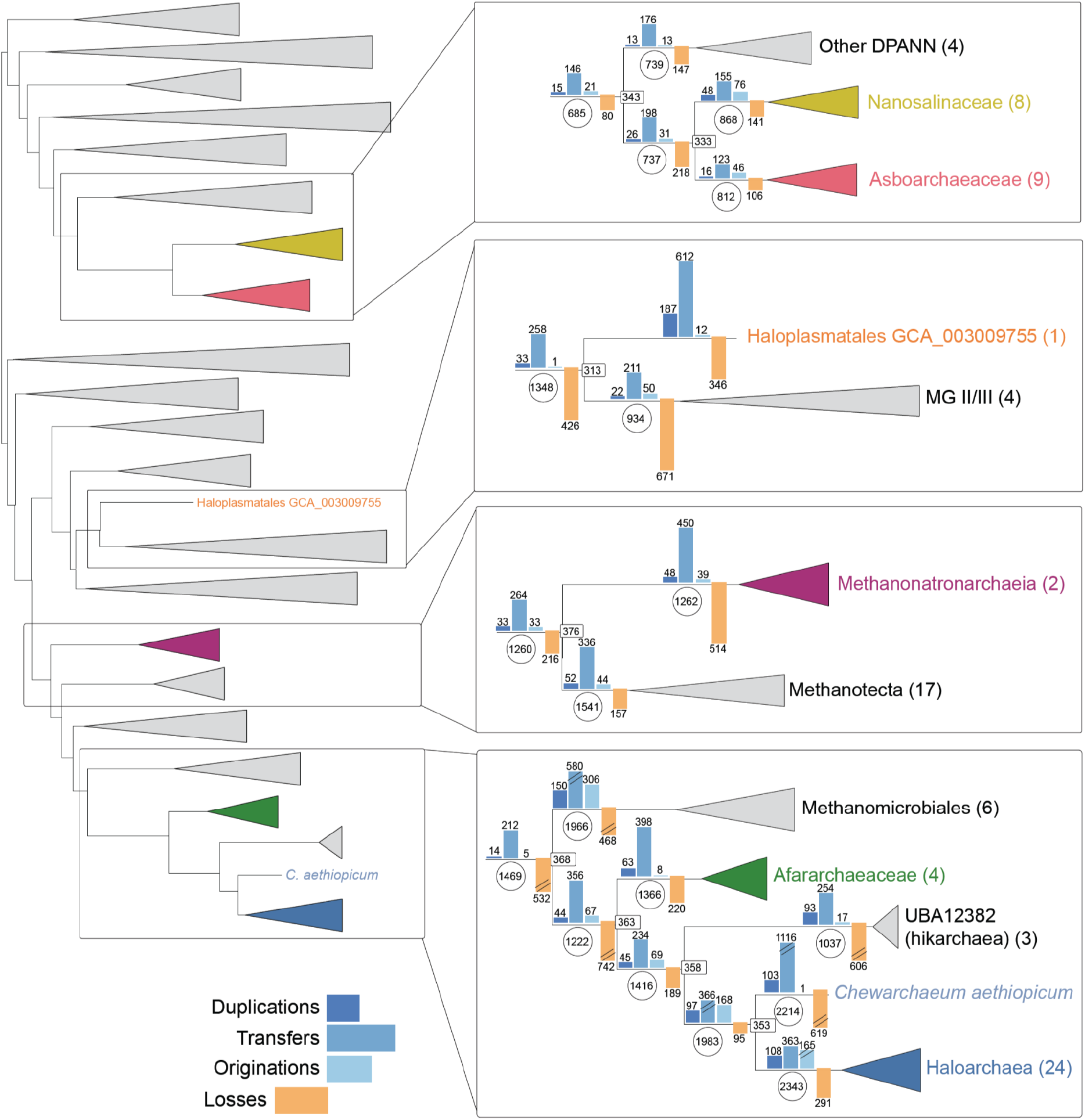
Schematic representation of the tree reconciliation analysis based on the NM species tree. The full archaeal tree is shown on the left; boxes on the right highlight the details for the four main groups of halophilic archaea: Nanosalinaceae+Asboarchaeaceae, Haloplasmatales, Methanonatronarchaeia; and Afararchaeaceae+Haloarchaea. The bar plots on the branches represent the number of gene duplications, transfers, originations, and losses, and the circles indicate the number of predicted ancestral genes. The number of taxa in each collapsed clade is indicated by the number in parentheses next to the clade name. The complete version of this tree with the events for all archaeal nodes and leaves can be found in Extended Data Fig. 8.

Although the extent and timing remain debated, massive HGT from bacteria appears to have played a significant role in the evolution of Haloarchaea^52–55^. Several of these transfers predated the separation of Afararchaeaceae and Haloarchaea and were most likely important in the adaptation of an ancestor of both groups to extreme halophily. One example is the choline dehydrogenase BetA (Extended Data Fig. 10b), which is involved in the biosynthesis of the osmoprotectant glycine-betaine^56^. Interestingly, this gene is not found in hickarchaea, reinforcing the idea that gene loss accompanied the secondary adaptation of this group to low-salt environments from an extremely halophilic ancestor. Another example is a transporter of the BCCT family involved in the uptake of osmoprotectants like glycine and betaine^56^, which Methanonatronarchaeia acquired from bacteria (Supplementary Fig. 26). Our tree reconciliation analysis supports that HGT between Haloarchaea and the other groups of halophilic archaea has also been influential in driving their convergent adaptations to extreme halophily. For example, this has been the case for the chaperone GrpE and several multi-copy haloarchaeal transporters cited before, including the K^+^ (Trk- and Kef-type), and Mg^2+^ transporters and the K^+^/H^+^, Ca^2+^/Na^+,^ and Na^+^/H^+^ antiporters. In addition to them, other transporters of inorganic molecules have also been transferred among the halophilic archaeal groups, such as SNF-family Na^+^-dependent transporters (Supplementary Fig. 27), ZupT- and FieF-type metal transporters (Supplementary Figs. 28 and 29, respectively), sulfur transporters (Supplementary Fig. 30), Na^+^/H^+^ antiporters (Supplementary Figs. 31 and 32), and Na^+^/phosphate symporters (Supplementary Fig. 33). HGT of transporters of organic molecules can also be observed, such as a transporter of di- and tricarboxylate Krebs cycle intermediates shared by Haloarchaea and Nanosalinaceae+Asboarcheaceae (Supplementary Fig. 34). In agreement with previous reports of inter-domain HGT followed by intra-domain HGT^57^, we detected several genes of bacterial origin encoding other transporters that have been subsequently transferred between different halophilic archaeal groups. They include an AmiS/UreI urea transporter, transferred between Haloarchaea and Nanohaloarchaea (Supplementary Fig. 35), and a TauE/SafE sulfite exporter, transferred between Haloarchaea and Methanonatronarchaeia (Supplementary Fig. 36).

## Conclusions

Living under salt-saturating conditions is challenging and requires coping with strong osmotic stress and maintaining the hydration state of cellular macromolecules against all odds^17^. For a long time, only one group of archaea, which generally excel in their adaptations to life-limiting conditions, was known to thrive in hypersaline systems, the Halobacteria or Haloarchea^1^. Haloarchaea are thought to have evolved from mildly halophilic methanogens acquiring many genes by HGT^54^ and developing a ‘salt-in’ adaptive strategy that implies pumping huge amounts of K^+^ into the cytoplasm and keeping molecular surfaces negatively charged^58^. In the case of proteins, this is achieved by including acidic amino acids, such that massive changes in the proteome of haloarchaea took place during evolution to adapt to extreme halophily. However, although they dominate hypersaline systems, haloarchaea are not alone, and other groups of halophilic archaea, including Nanohaloarchaeota, Methanonatronoarchaeia and Haloplasmatales, and even some bacteria (Salinibacteraceae), can cope with these extreme conditions^1,2^. Did the adaptation to extreme halophily in archaea evolve only once or several times (and if so, how many)? To answer, a resolved phylogenetic tree including these archaea is needed, but this task has been largely hampered by the excessively biased nature of their acidic proteome with, in particular, Nanohaloarchaeota and Methanonatronarcheia displaying incongruent positions across previous studies^4,6,8,10–13,43^. Here, we have achieved this task by improving the taxon sampling and using different approaches to cope with compositional biases, including the application of a specific model of sequence evolution and the removal of the most biased positions from phylogenomic analyses. This was possible by incorporating two newly discovered archaeal lineages from geothermally influenced hypersaline settings^22^, Asboarchaeaceae, and Afararchaeaceae, that strategically and robustly branch sister to, respectively, Nanosalinaceae (Nanohaloarchaeota), within the DPANN, and the Haloarchaea plus their sister group, the marine Group IV^16^ or ‘hikarchaea’^10^ (GTDB family UBA12382^15^). Accordingly, ‘hikarchaea’ do not represent an intermediate between methanogenic archaea and haloarchaea as previously thought^10^ but secondarily adapted to low salinity from an extremely halophilic ancestor. In addition, our phylogenomic analyses robustly place Methanonatronarchaeia in a deep-branching position sister to the Methanotecta. This suggests that contrary to the initial proposal^6^, Methanonatronarchaeia are not evolutionary intermediates between Class II methanogens and haloarchaea. Based on our resolved archaeal tree, we conclude four-independent adaptations to extreme halophily in archaea: in Haloarchaea+Afararchaeia, in Methanonatronarchaeia, in Haloplasmatales, and in Nanosalinaceae+Asboarchaeaceae. A salt-in strategy was adopted in the four cases, with extensive concomitant acidification of the proteomes.

While convergent evolution independently led to the massive adaptation of the proteome to high intracellular K^+^ levels, HGT seems to have also played an important role in this process by spreading key genes (such as ion transporters) among the various halophilic lineages. This opens the question of whether the key initial adaptation(s) to extreme halophily evolved only once and spread by HGT and which lineage of extreme halophiles evolved first. Identifying and studying the distribution and phylogeny of such adaptive genes in known and potentially novel halophilic archaea and determining the directionality of HGT involving those genes should help unravel the evolutionary history of this fascinating adaptation to salty extremes.

### *Candidatus* “*Afararchaeum irisae*” (gen. nov., sp. nov.)

“Afar” refers to the Afar region (northeastern Ethiopia) where this organism has been found. The species is named after the Iris Foundation (France), which supports the study and preservation of endangered ecosystems such as those in the Afar region. This halophilic archaeon lives in oxic hypersaline waters. It encodes genes for aerobic respiration and likely uses amino acids for organoheterotrophic growth. Its genome is around 1.9 Mbp with a GC content of 55%. It currently remains uncultured and known from environmental sequencing only, with one MAG presented here. DAL-WCL_na_97C3R is the designated type MAG.

### Description of Afararchaeaceae (fam. nov.)

Description is the same as for the genus *Afararchaeum*. Suff. -aceae, ending to denote a family. Type genus: *Afararchaeum* gen. nov.

### *Candidatus* “Asboarchaeum danakilensis” (gen. nov., sp. nov.)

“Asbo” means “salt” in the Afar language spoken in the northeastern Ethiopia region where this organism has been found. This halophilic archaeon lives in oxic hypersaline waters of the Danakil Depression. It has a streamlined genome (around 1.2 Mb) with a relatively high GC content (61%). It lacks most biosynthetic pathways (for amino acids, nucleosides, nucleotides, and phospholipids), so most likely it grows as a symbiont of an unknown host. It currently remains uncultured and known from environmental sequencing only, with one MAG presented here. DAL-WCL_45_84C1R is the designated type MAG.

### Description of Asboarchaeaceae (fam. nov.)

Description is the same as for the genus *Asboarchaeum*. Suff. -aceae, ending to denote a family. Type genus: *Asboarchaeum* gen. nov.

### *Candidatus* “*Chewarchaeum aethiopicum*” (gen. nov., sp. nov)

“Chew” means “salt” in the Amharic language spoken as the official language of Ethiopia, where this organism has been found. This halophilic archaeon lives in oxic hypersaline waters of the Danakil Depression. It encodes genes for aerobic respiration and likely uses amino acids for organoheterotrophic growth. Its genome is around 2.9 Mb with a relatively high GC content (61%). It currently remains uncultured and known from environmental sequencing only, with one MAG presented here. DAL-9Gt_70_90C3R is the designated type MAG.

### Description of *Chewarchaeaceae* (fam. nov.)

Description is the same as for the genus *Chewarchaeum*. Suff. -aceae, ending to denote a family. Type genus: *Chewarchaeum* gen. nov.

## Methods

### Selection of metagenome-assembled genomes

We searched for MAGs related to known groups of extremely halophilic archaea in the Danakil Depression dataset obtained by Gutiérrez-Preciado et al.^22^. For this, we included 61 Danakil MAGs in a preliminary phylogenetic tree containing 427 representatives of archaeal diversity and constructed a phylogenetic tree using 56 concatenated ribosomal proteins with IQ-TREE v1.6.10^59^. The tree was built using the LG+C20+F+G model of sequence evolution, and support at branches was estimated from 1000 ultrafast bootstrap replicates. From this analysis, we selected 14 high-quality MAGs (>50% completeness, ≤5% redundancy) representing potential new groups of extremely halophilic archaea based on their position compared to other halophilic archaea. These 14 MAGs were taxonomically classified using GTDB-Tk^60^ (version 2.3.0, r207; April 1st, 2022) and assigned to novel families within three GTDB orders: four MAGs were assigned to a novel family belonging to the order ‘JAHENH01’, which we have named Afararchaeaceae; nine MAGs were assigned to another novel family belonging to the order Nanosalinales, which we propose to name Asborarchaeaceae; and one MAG belonged to a third novel family in the order Halobacteriales, which we have named Chewarchaeaceae (see taxonomic description above for more details).

### Metagenome-assembled genome annotation

Coding DNA sequences (CDSs) were predicted with Prodigal v2.6.3^61^ and subjected to Pfam^33^ and COG^62^ functional annotations inside the Anvi’o v5 pipeline^63^. Genes were also annotated with KofamKOALA^64^ and eggNOG-mapper v2.1.5^34^. Additional manual curation was done for the two most complete Afararchaeaceae and Asboarchaeaceae MAGs (DAL-WCL_na_97C3R and DAL-WCL_45_84C1R, respectively). Further information on gene annotations and functional predictions can be found in Supplementary Data 1 and 2.

### Detecting novel protein families in Afararchaeaceae and Asboarchaeaceae

We computed family clusters of the proteins predicted for the MAGs of the new archaeal families Afararchaeaceae and Asboarchaeaceae using MMseqs2^65^ with relaxed thresholds: minimum percentage of amino acids identity of 30%, e-value <1e-3, and a minimum sequence coverage of 50% (--min-seq-id 0.3 -c 0.5 --cov-mode 2 --cluster-mode 0). To detect families with no homologs in reference databases, we mapped i) the protein sequences encoded in the MAGs against EggNOG using eggNOG-mapper v2^34^ (hits with an e-value <1e-3 were considered as significant) ii) the protein sequences encoded in the MAGs against PfamA domains using HMMER^66^ (hits with an e-value <1e-5 were considered as significant), iii) the protein sequences encoded in the MAGs against PfamB domains using HMMER^66^ (hits with an e-value < 1e-5 were considered as significant) and iv) the CDS sequences of the MAGs against RefSeq using diamond blastx^67^ (‘sensitive’ flag, hits with an e-value <1e-3 and query coverage >50% were considered as significant). We only considered novel families those with no detectable homologs in these databases. To address the taxonomic breadth of the novel families, we mapped the longest sequence of each family against the proteins encoded in a collection of 169,484 genomes, including non-cultured species, coming from diverse sequencing efforts and spanning the prokaryotic tree of life using diamond blastp^67^ (‘sensitive’ flag, hits with an e-value <1e-3 and query coverage >50% were considered as significant). We then expanded each protein family with the hits from this database. If, after expanding, a family incorporated genes with homologs in EggNOG, that family was then discarded from the novel family set. We predicted signal peptides and transmembrane domains on the gene families using SignalP^68^ and TMHMM^69^. Protein families were considered as transmembrane or exported if >80% of their members had a predicted transmembrane domain or a signal peptide, respectively.

### Phylogenetic analyses

We collected the proteomes of 192 taxa spanning all major archaeal super-groups (including the new Afararchaeaceae and Asboarchaeaceae). We reconstructed two phylogenomic datasets consisting of 48 ribosomal proteins (RP) and 136 new markers (NM). The 136 NM dataset was based on curating a set of 200 markers previously shown to be highly conserved across the archaeal domain^11^. To ensure standardized protein-coding gene predictions, all 192 genomes were first run through Prodigal^61^. Next, sequences similar to the RP and NM proteins were identified using BLAST^71^ with relatively relaxed criteria (>20% sequence identity over 30% query length) to retrieve even divergent homologs, such as those found in fast-evolving lineages like the DPANN archaea. For each of the 192 taxa, up to five BLAST hits were kept to ensure that we could detect cases of contamination, HGT, or paralogy and identify the correct orthologue for each taxon. This required multiple rounds of manual curation based on examining single protein trees (reconstructed with FastTree2^72^). Once manually verified, each orthologous group was aligned with MAFFT L-INS-i v7.450^73^ and trimmed with BMGE v1.12^74^ (-m BLOSUM30 -b 3 -g 0.2 -h 0.5). We performed a final round of verification of the single gene trees reconstructed using the more sophisticated LG+C60+ F+G4 model in IQ-TREE before concatenating the individually trimmed alignments into two super matrices (RP and NM). The 192-RP and 192-NM alignments were then subsampled to generate two additional alignments consisting of 87 taxa containing only Euryarchaea (87-RP and 87-NM) and 104 taxa, including the 87 Euryarchaea plus 8 Nanosalinaceae and 9 Asboarchaeaceae (104-RP and 104-NM). These six alignments were then used for maximum likelihood (ML) phylogenetic reconstruction under the LG+C60+F+G4 sequence evolution model (with 1000 ultra-fast bootstrap replicates) using IQTREE v2.0.3^75^. For four of the six alignments (87-RP, 104-RP, 87-NM, and 104-NM), Bayesian phylogenetic reconstructions were also run using the CAT+GTR model as implemented in PhyloBayes v1.8^76^. Four MCMC chains were run in parallel for each alignment until a sufficient effective sample size was reached (effsize >300) while using a burnin of 3000 cycles and sampling every 50 generations after the burn-in.

### Amino acid composition analysis

We used an in-house Python script (https://github.com/bbaker567/phylogenetics) to estimate the frequency of each amino acid in our selection of 192 archaeal taxa for the whole predicted proteomes, as well as for the RP and NM datasets. These frequencies were analyzed using principal component analysis with ggplot2 ^77^.

In addition, for each amino acid, the compositional bias between halophiles and non-halophiles was measured for the RP and NM datasets with the Z-score from a binomial test of two proportions:

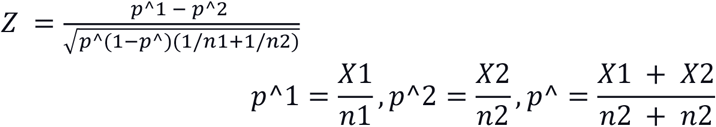

where X_1_ and X_2_ are the total numbers of that amino acid, and n_1_ and n_2_ are the total numbers of all 20 amino acids across halophiles and non-halophiles, respectively. Calculating Z-scores in this way assumes that the proportions of an amino acid across halophiles and non-halophiles are approximately normal, with the null hypothesis that p_1_ = p_2_. |Z| >1.96 indicates rejection of the null hypothesis at a significance level of p <0.05. Amino acids with |Z| >1.96 were considered significantly enriched in halophiles relative to non-halophiles, whereas amino acids with |Z| <-1.96 were considered significantly depleted in halophiles relative to non-halophiles. Amino acids were divided into ‘Over-represented’ (|Z| >1.96), ‘Under-represented’ (|Z| <-1.96), and ‘Not significant’ (|Z| not statistically significant).

We also implemented the new GFmix-DE/IK model by transforming the b parameter of the GFmix model^42^ (originally designed to represent the ratio of GARP/FYMINK amino acids across all descendant taxa at each branch in a tree) to accommodate amino acid groupings other than GARP/FYMINK, in our case those identified to be biased in extreme halophiles. We then calculated the likelihood of different tree topologies under these variants of the GFmix model with LG+C60+F+G4^42^. Branch length and alpha shape parameters for each tree tested were estimated using IQTREE v2.0.3^75^ and then fed into GFmix, specifying the custom enriched and depleted amino acid bins for halophiles versus non-halophiles.

### Progressive removal of compositionally biased sites

To remove the most compositionally biased sites from the sequence datasets, we split the sequence alignments in two based on whether the taxa were classified as extreme halophiles or non-halophiles. We then calculated the ratio of D+E divided by I+K for each alignment site for both the halophiles and non-halophiles sub-alignments. We then divided the D+E/I+K ratio for each halophile sub-alignment site by the corresponding ratio in the non-halophile sub-alignment. When the denominator of one of the ratios was equal to zero, we substituted ‘0’ for ‘0.1’ in order to still consider the alignment position. Alignment sites were then ranked from the highest to the lowest ratio, using the highest ratio as a proxy for the most biased alignment site. Next, we progressively removed alignment sites in increments of 1%, 5%, 10%, 20%, 30%, and up to 90%. This resulted in 11 alignments for both the RP and NM datasets. These 11 alignments were then used for ML phylogenetic reconstruction under the LG+C60+F+G4 model (with 1000 ultra-fast bootstraps).

### Orthologous groups and single-gene trees

Orthologous groups (OGs) were identified for all the proteins of the species included in the 192 taxa dataset using OrthoFinder v2.5.1^78^ with Diamond BLAST (--ultra-sensitive, --query-cover 50%, and --id 30%) and an inflation parameter of 1.1^78^. This resulted in 17,827 OGs, which were aligned using MAFFT --auto v7.450^73^ with default settings and trimmed using trimAl^79^ (-automated1 - resoverlap 0.75 -seqoverlap 75). To avoid poorly resolved single gene trees due to little phylogenetic information, we removed OGs that presented a trimmed alignment length of less than 60 amino acids. This resulted in 17,288 OGs, which were used to reconstruct individual trees with IQTREE v2.0.3^75^. For computational time reasons, the trees of the 200 OGs containing the largest number of sequences were inferred under the LG+C20+F+G4 model of evolution, while the remaining phylogenies were run under LG+C60+F+G4. Statistical support at branches were estimated using 1,000 ultrafast bootstrap replicates. Finally, for OGs containing only two or three sequences, “bootstrap” samples were artificially generated for subsequent analysis in ALE^45^, corresponding to the single possible unrooted tree topology.

### Gene tree-aware ancestral reconstruction

The 17,288 single-gene trees were reconciled with the species tree inferred from the 192-NM dataset using the ALEml_undated algorithm of the ALE suite v0.4^45^. ALE infers, for each gene family, duplications, losses, transfers, and originations events along a species tree^45^. These events were counted only if the relative reconciliation frequencies output by ALE were at least 0.3, following the recommendations of previous analyses^10,39,80^. These relative frequency values support an evolutionary event occurring at a given node by incorporating the uncertainty of the reconstructed individual gene tree, as represented by the bootstrap replicates. ALE also predicts the ancestral copy number for each node in the species tree. Phylogenetic trees were visualized using Figtree v.1.4.4 (http://tree.bio.ed.ac.uk/software/figtree), iTOL^81^, and the ETE3 Toolkit v.3.1.2^82^.

## Data availability

The MAGs reported in this study have been deposited in GenBank under BioProject number PRJNA901412. All raw data underlying phylogenomic analyses (raw and processed alignments and corresponding phylogenetic trees) and all predicted proteomes have been deposited into Figshare (https://figshare.com/account/home#/projects/154868).

## Code availability

Custom code used for data analysis is available at GitHub: (https://github.com/bbaker567/phylogenetics).

## Acknowledgments

D.M. and L.E were supported by grants from the European Research Council (ERC Advanced grant 787904 and ERC Starting grant 803151, respectively). This work was also supported by the Moore-Simons Project Call on the Origin of the Eukaryotic Cell, Simons Foundation 812811 (A.J.R, E.S., and L.E.), and Moore Foundation GBMF9739 (P.L.G.). We thank P. Deschamps for help in managing our bioinformatic cluster. We are grateful to the Iris Foundation for the continuous support of our work on the microbial diversity of the Danakil Depression.

## Author contributions

D.M., P.L.G., and L.E designed the study. A.G.P. and B.B. annotated the new archaeal MAGs. A.R.R., B.B., and J.H.C. studied the new protein families. C.G.P.MC., A.J.R., and E.S. conceived of the binomial methods to identify significant shifts in amino acid composition, and E.S. implemented the new features of the GFmix model in the GFmix software. B.B., L.E., D.M., C.G.P.M., A.J.R., and E.S. carried out phylogenetic analyses. B.B., L.E., P.L.G., and D.M. wrote the paper with contributions from all authors.

