## Supplementary information for "Several independent adaptations of archaea to hypersaline environments"

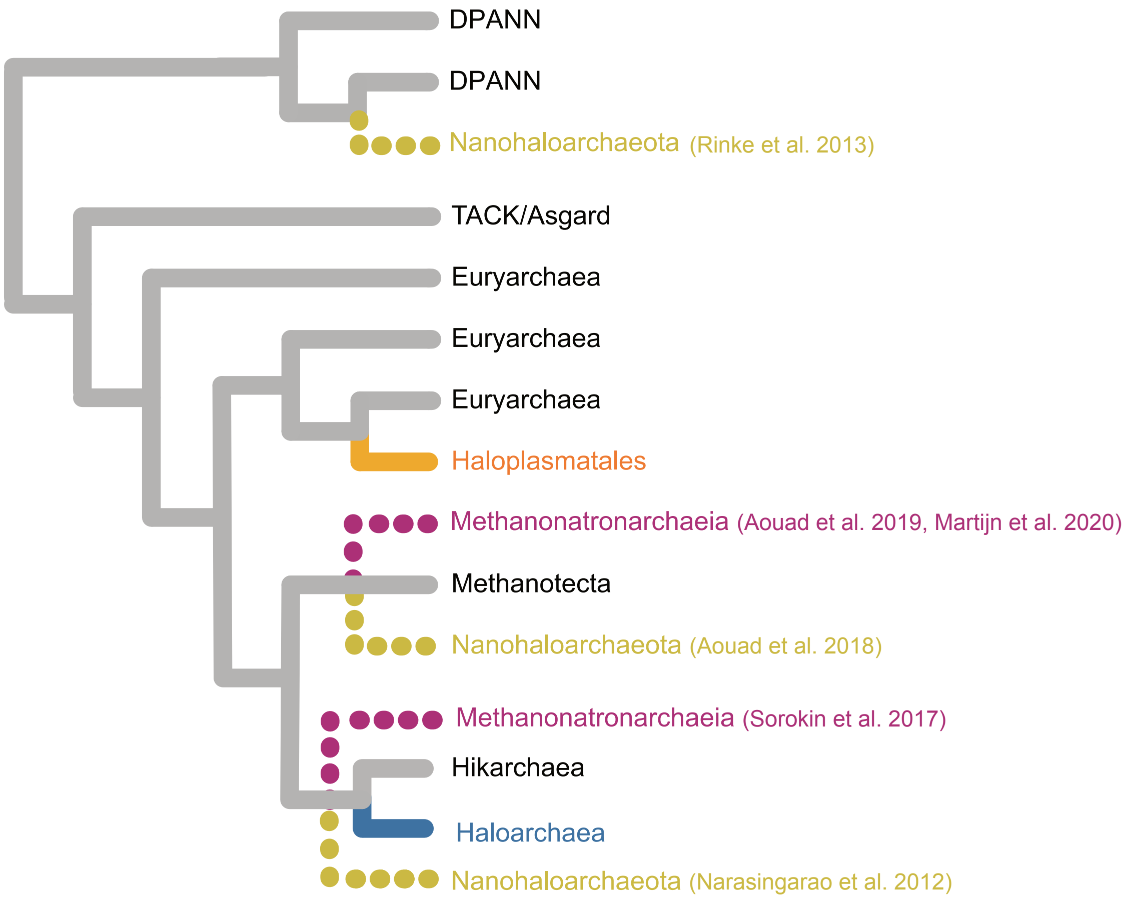


**Extended Data Fig. 1 | Schematic tree showing the phylogenetic position of extremely halophilic archaeal groups (colored branches) proposed in previous articles.** Branches that have been found at different places in the tree of archaea are indicated with dashed lines (Narasingarao et al. 2012, Rinke et al. 2013, Sorokin et al. 2017, Aouad et al. 2018, Aouad et al. 2019, Martijn et al. 2020).

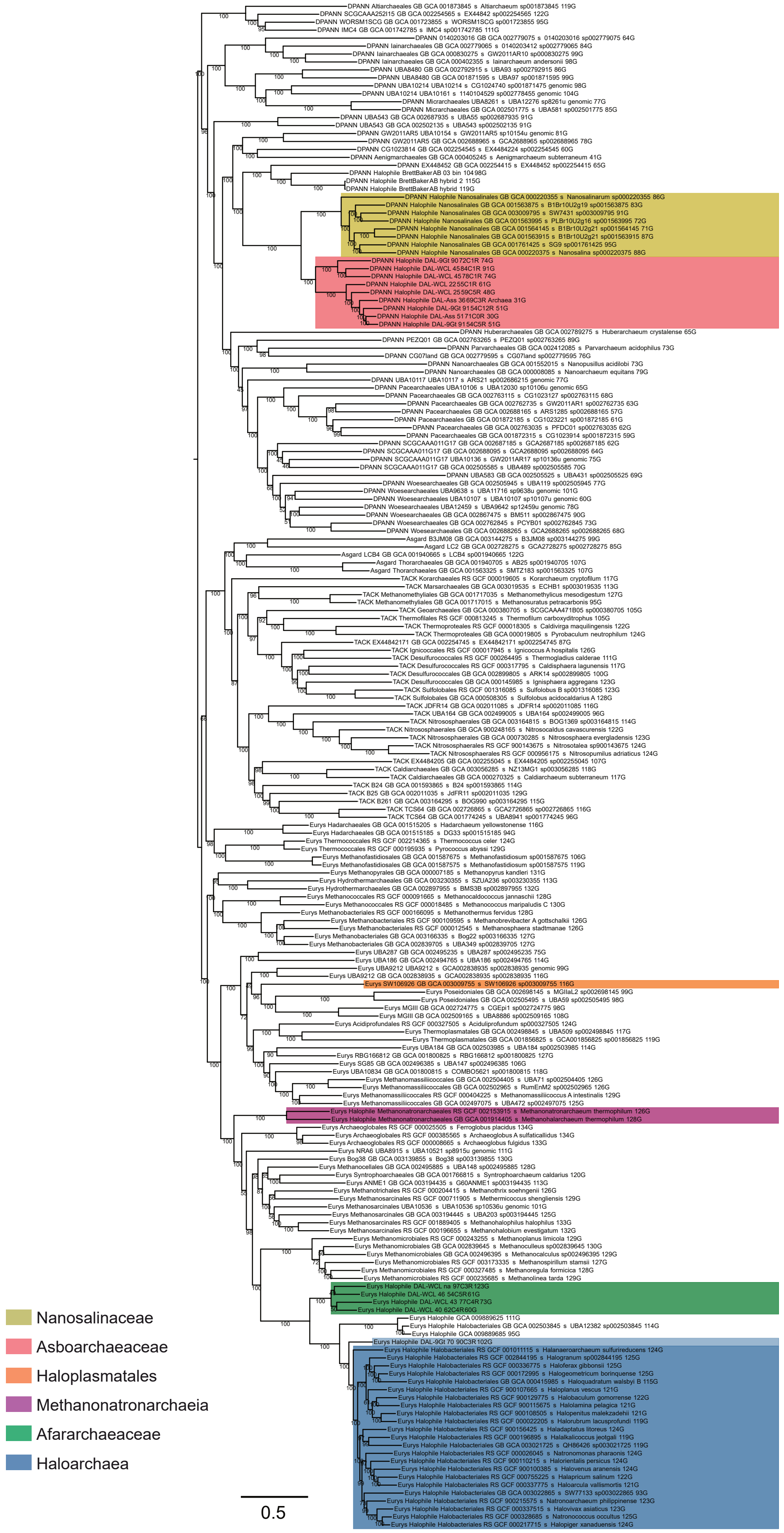


**Extended Data Fig. 2 | Maximum likelihood phylogeny of 192 archaea based on the NM dataset.** The ML tree was inferred with the LG+C60+F+Γ4 model of evolution with 1000 ultrafast bootstraps as implemented by IQ-TREE. Distinct halophilic clades are indicated in color, including the Nanosalinaceae (golden yellow), Asboarchaeaceae (salmon), Haloplasmatales (orange), Methanonatronarchaeia (purple), Afarachaeaceae (green), and Haloarchaea (blue). Scale bar indicates the expected average number of substitutions per site.

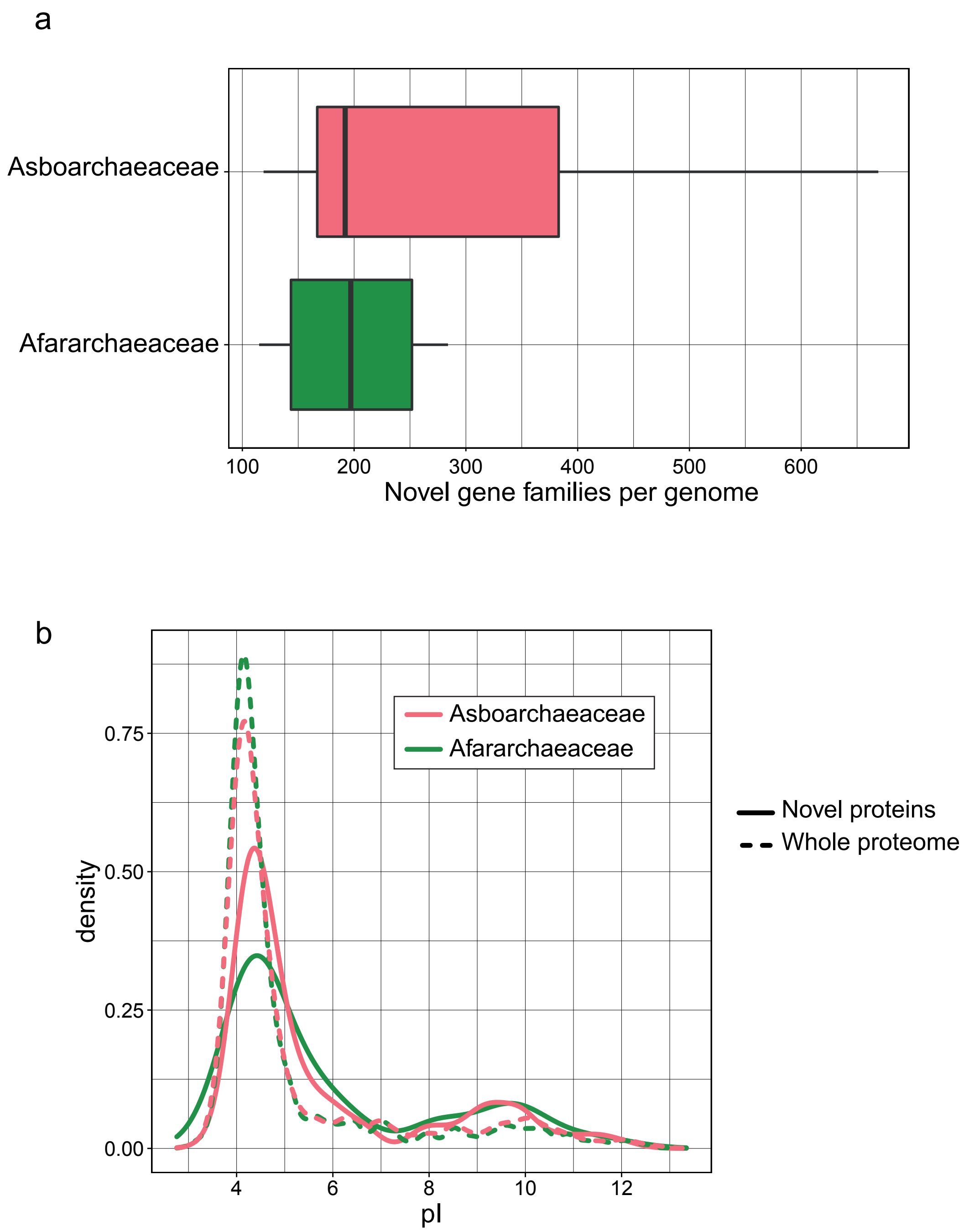


**Extended Data Fig. 3 | Number and isoelectric point of novel gene families identified in the Asboarchaeaceae and Afararchaeaceae MAGs. (a)** Average number of novel genes in the nine asboarchaeal and four afarachaeal MAGs described in this study (see Methods). **(b)** Isoelectric point of these novel proteins (solid lines) compared to the average isoelectric point of the whole proteomes (dashed lines).

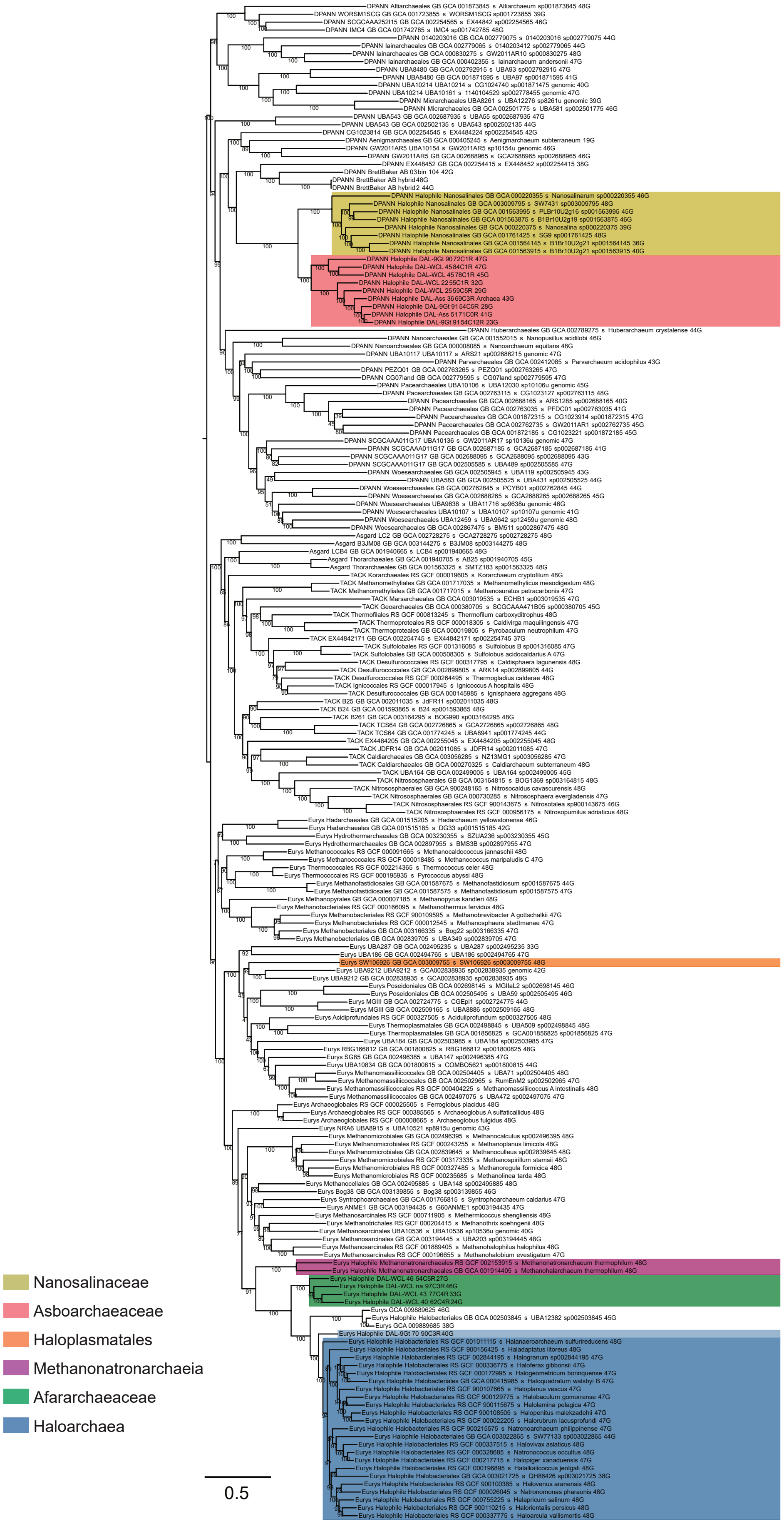


**Extended Data Fig. 4 | Maximum likelihood phylogeny of 192 archaea built with the RP dataset.** The tree was inferred under the LG+C60+F+Γ4 model of sequence evolution with 1000 ultrafast bootstraps as implemented by IQ-TREE. Distinct halophilic clades are indicated in color, including the Nanosalinaceae (golden yellow), Asboarchaeaceae (salmon), Haloplasmatales (orange), Methanonatronarchaeia (purple), Afarachaeaceae (green), and Haloarchaea (blue). Scale bar indicates the expected average number of substitutions per site.

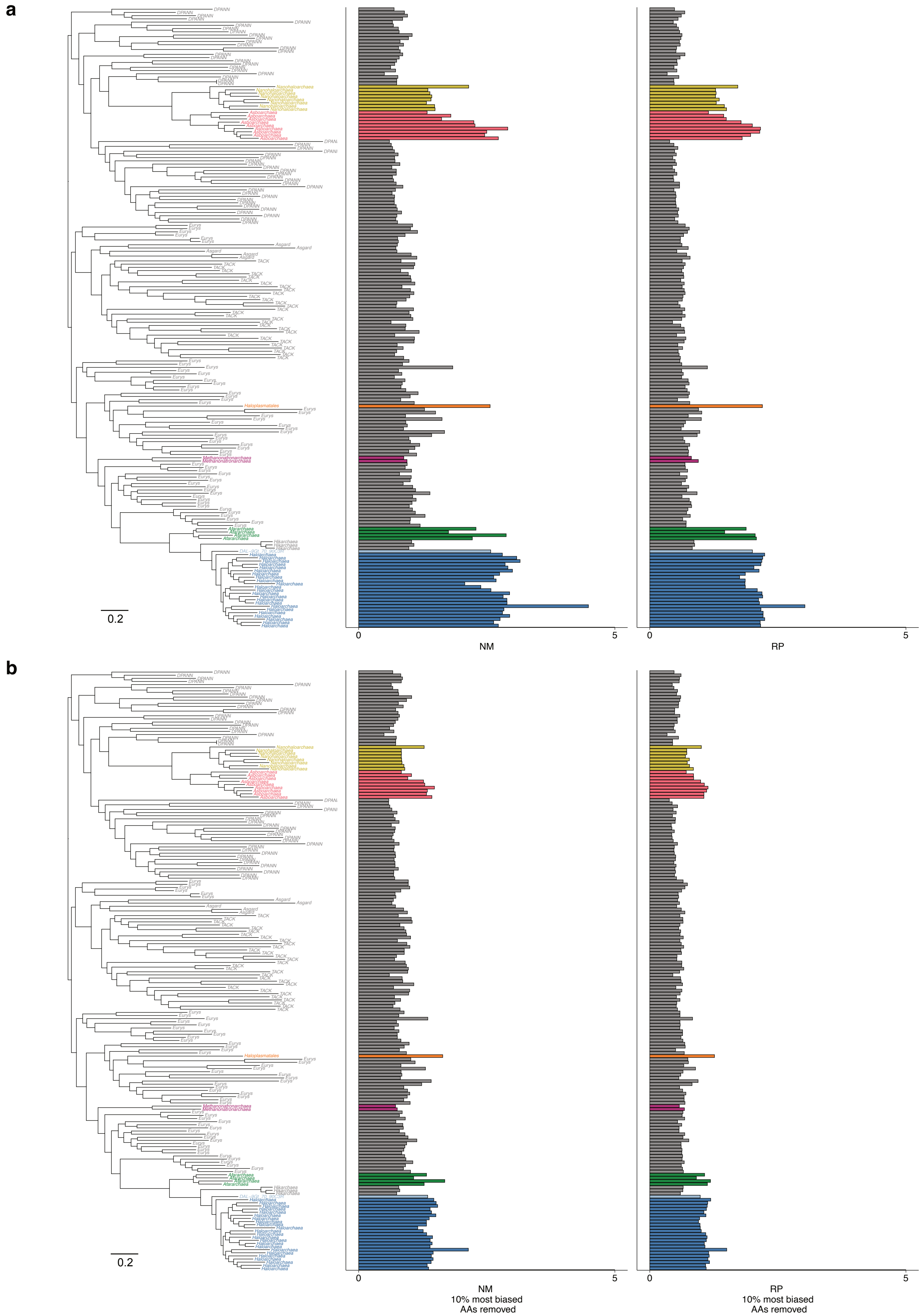


**Extended Data Fig. 5 | Halophilic-specific amino acid compositional biases along the phylogeny of 192 archaeal taxa. (a)** The ratio of [D+E/I+K] amino acids of 192 archaeal taxa calculated along the untreated NM and RP alignments (39,385 and 6,792 amino acid positions, respectively). **(b)** 10% of the most biased sites (i.e. those with the highest ratio) removed from the NM and RP alignments. Distinct halophilic clades are indicated in color, including the Nanosalinaceae (golden yellow), Asboarchaeaceae (salmon), Haloplasmatales (orange), Methanonatronarchaeia (purple), Afarachaeaceae (green), and Haloarchaea (blue). Scale bar indicates the expected average number of substitutions per site.

a

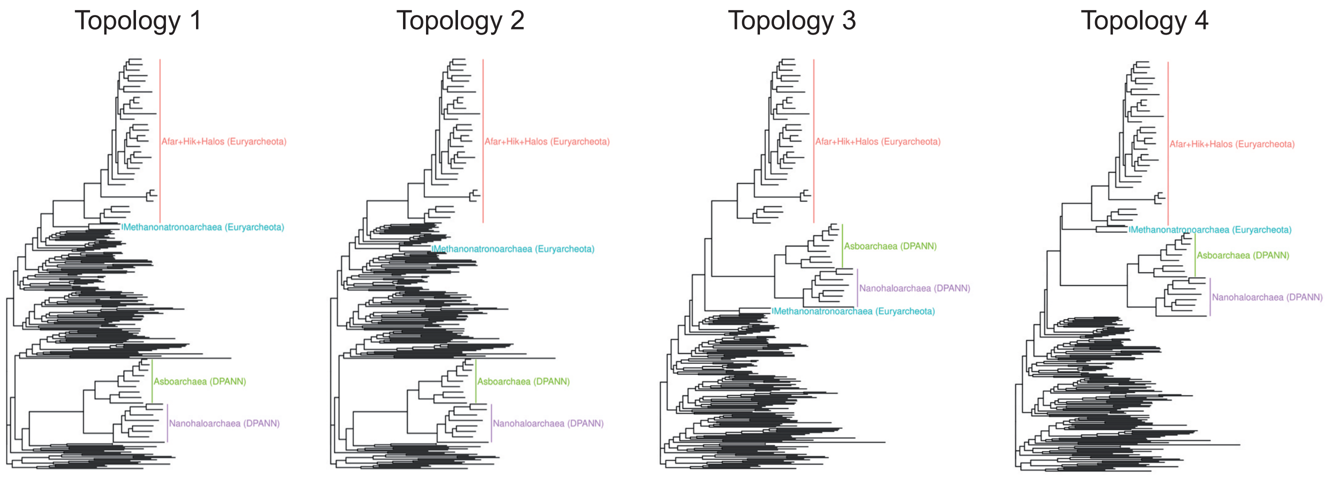


| 192-taxa RP dataset | | | | | | |
|---|---|---|---|---|---|---|
| **Model** | **Bin 1** | **Bin 2** | **Topology 1** | **Topology 2** | **Topology 3** | **Topology 4** |
| LG+C60+G+F | None | None | Likelihood=-1317976,09 (Total diff.=0) | Likelihood= -1318026,107 (Total diff.= -50,078) | Likelihood= -1318569,945 (Total diff.= -593,916) | Likelihood= -1318599,398 (Total diff.= -623,369) |
| LG+C60+G+F+GFmix | DE | IK | Likelihood=-1314937,111 (Total diff.=0) | Likelihood= -1314977,65 (Total diff.= -40,539) | Likelihood= -1315670,918 (Total diff.= -733,808) | Likelihood= -1315685,576 (Total diff.= -748,465) |
| LG+C60+G+F+GFmix | DEQTAVG | KILCMFYWS | Likelihood=-1315018,218 (Total diff.=0) | Likelihood= -1315065,115 (Total diff.= -46,898) | Likelihood= -1315704,916 (Total diff.= --686,698) | Likelihood= -1315727,147 (Total diff.= -708,93) |

b

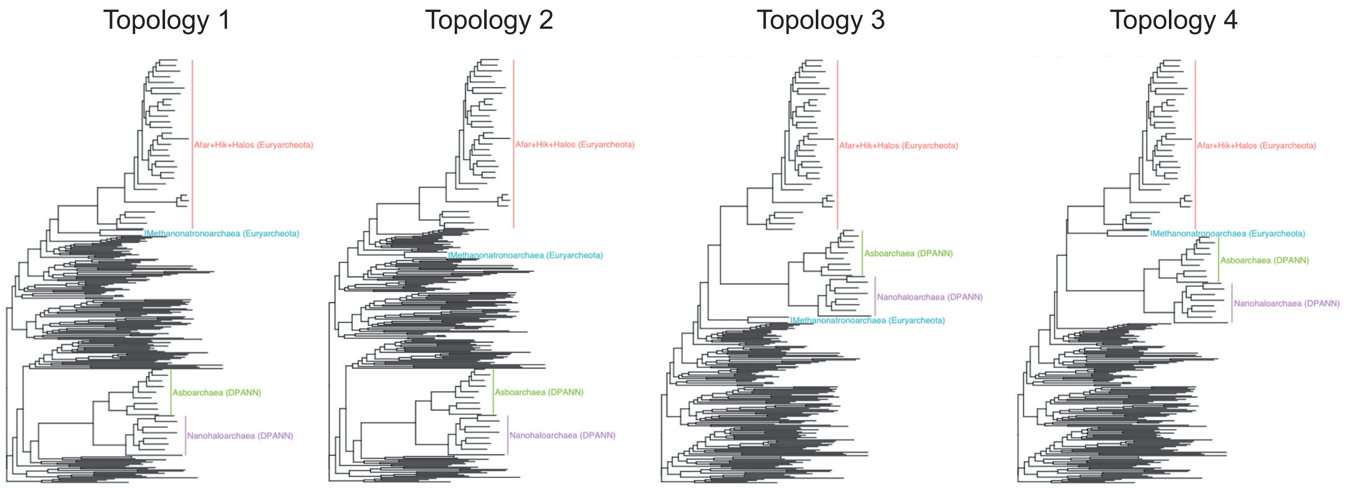


| 192-taxa PNM dataset | | | | | | |
|---|---|---|---|---|---|---|
| **Model** | **Bin 1** | **Bin 2** | **Topology 1** | **Topology 2** | **Topology 3** | **Topology 4** |
| LG+C60+G+F | None | None | Likelihood= -6557969,994 (Total diff.= -330,418) | Likelihood=-6557639,577 (Total diff.=0) | Likelihood= -6561223,122 (Total diff.= -3583,545) | Likelihood= -6561620,115 (Total diff.= -3980,538) |
| LG+C60+G+F+GFmix | DE | IK | Likelihood= -6547076,576 (Total diff.= -364,896) | Likelihood=-6546711,68 (Total diff.=0) | Likelihood= -6550656,783 (Total diff.= -3945,103) | Likelihood= -6550979,577 (Total diff.= -4267,898) |
| LG+C60+G+F+GFmix | DEAVTQGRHP | KINFSCLYM | Likelihood= -6541714,646 (Total diff.= -309,156) | Likelihood=-6541405,489 (Total diff.=0) | Likelihood= -6545093,421 (Total diff.= -3687,931) | Likelihood= -6545459,741 (Total diff.= -4054,252) |

**Extended Data Fig. 6 | Likelihood values for alternative positions of the extremely halophilic archaeal lineages.** Likelihoods are calculated using IQ-TREE with the LG+C60+G+F model alone or combined with the new GFmix model (taking into account all significantly enriched (Bin 1) or depleted (Bin 2) amino acids in halophiles or only the most extremely biased ones (D+E and I+K). The highest-scoring topology is indicated with a red rectangle for the **(a)** RP and **(b)** NM datasets. Likelihood differences between a given topology and the highest-scoring topology per model are given in parentheses.

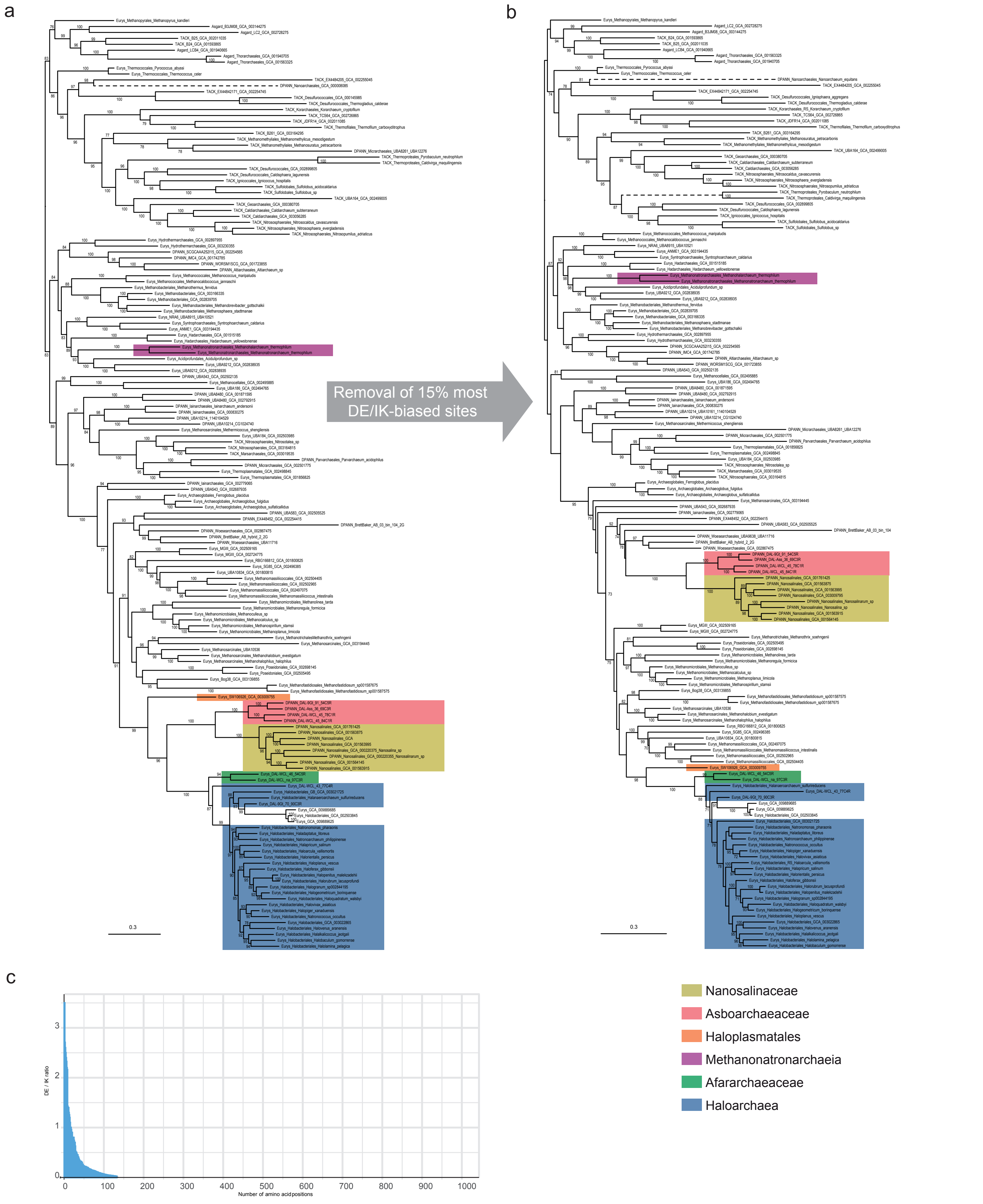


**Extended Data Fig. 7 | Impact of compositional bias on the phylogeny of archaeal ATP synthase.** Maximum likelihood phylogenetic trees based on the concatenation of ATP synthase subunits A and B **(a)** before and **(b)** after removal of 15% of sites with the highest D+E/I+K ratio. Distinct halophilic clades are indicated in color, including the Nanosalinaceae (golden yellow), Asboarchaeaceae (salmon), Haloplasmatales (orange), Methanonatronarchaeia (purple), Afarachaeaceae (green), and Haloarchaea (blue). Notice the shift in the position of the Nanosalinaceae+Asboarchaeaceae group. The trees were reconstructed using the LG+C60+G+F model of sequence evolution. Numbers at branches indicate ultrafast bootstrap support values. Only values >70% are indicated. Scale bar indicates the expected average number of substitution per site. **(c)** D+E/I+K ratio for all sites in the ATP synthase subunits A and B dataset ordered from highest to lowest values.

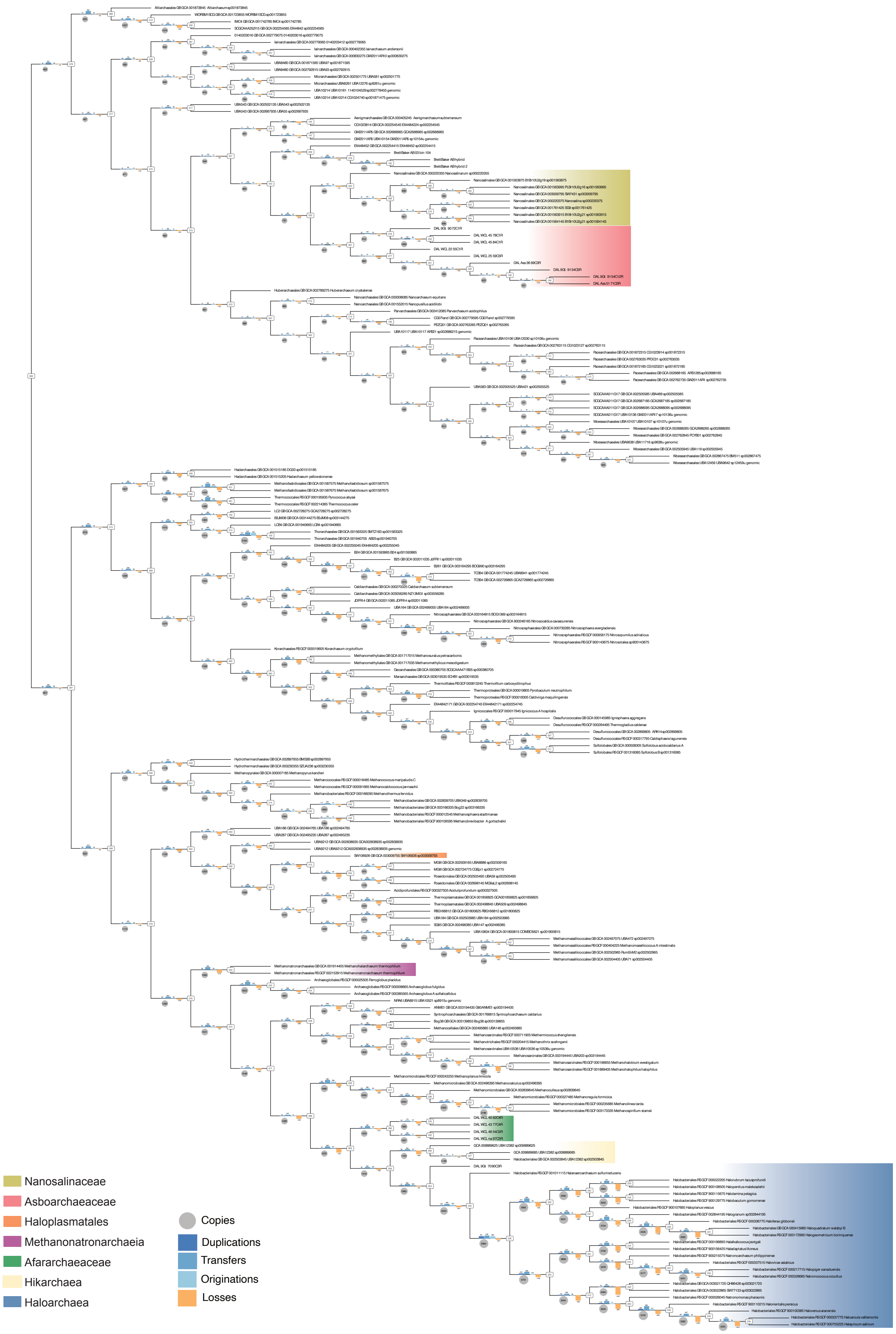


**Extended Data Fig. 8 | Ancestral proteome sizes and numbers of gene loss, duplication, and gain inferred from reconciliation analyses across 192 archaeal taxa.** Only gene copy and event numbers with reconciliation frequencies ≥ 0.3 were considered (see Methods). Barplots at each branch indicate duplication, transfer, and origination events (blue bars; see legend), and loss events (orange bars). Each grey circle indicates the inferred proteome size (i.e., number of protein-coding gene copies) for the ancestor to the right of the branch. Numbers in rounded rectangles correspond to the node identifiers. Halophilic clades are colored.

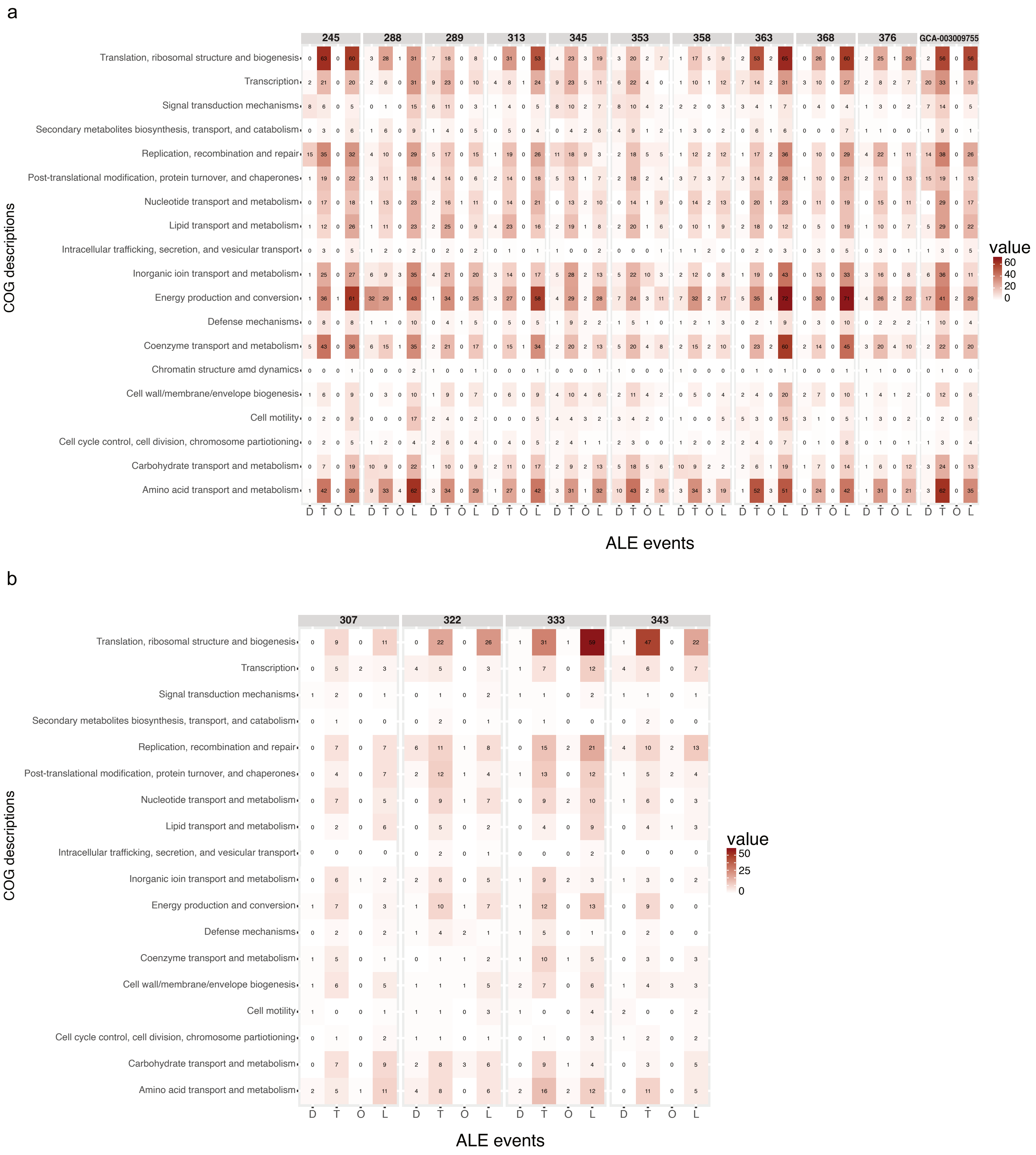


**Extended Data Fig. 9 | Heat map of the number of gene duplications, transfers, originations and losses in various archaeal halophilic lineages according to their COG classification.** The counts were obtained using the amalgamated likelihood estimation (ALE) tree reconciliation method on the set of 17,288 orthologous genes present in the 192-taxa genomic dataset for several nodes within the **(a)** Euryarchaeota and the **(b)** DPANN archaea (see Methods). Node numbers correspond to the nodes in the complete tree shown in Extended Data Fig. 8.

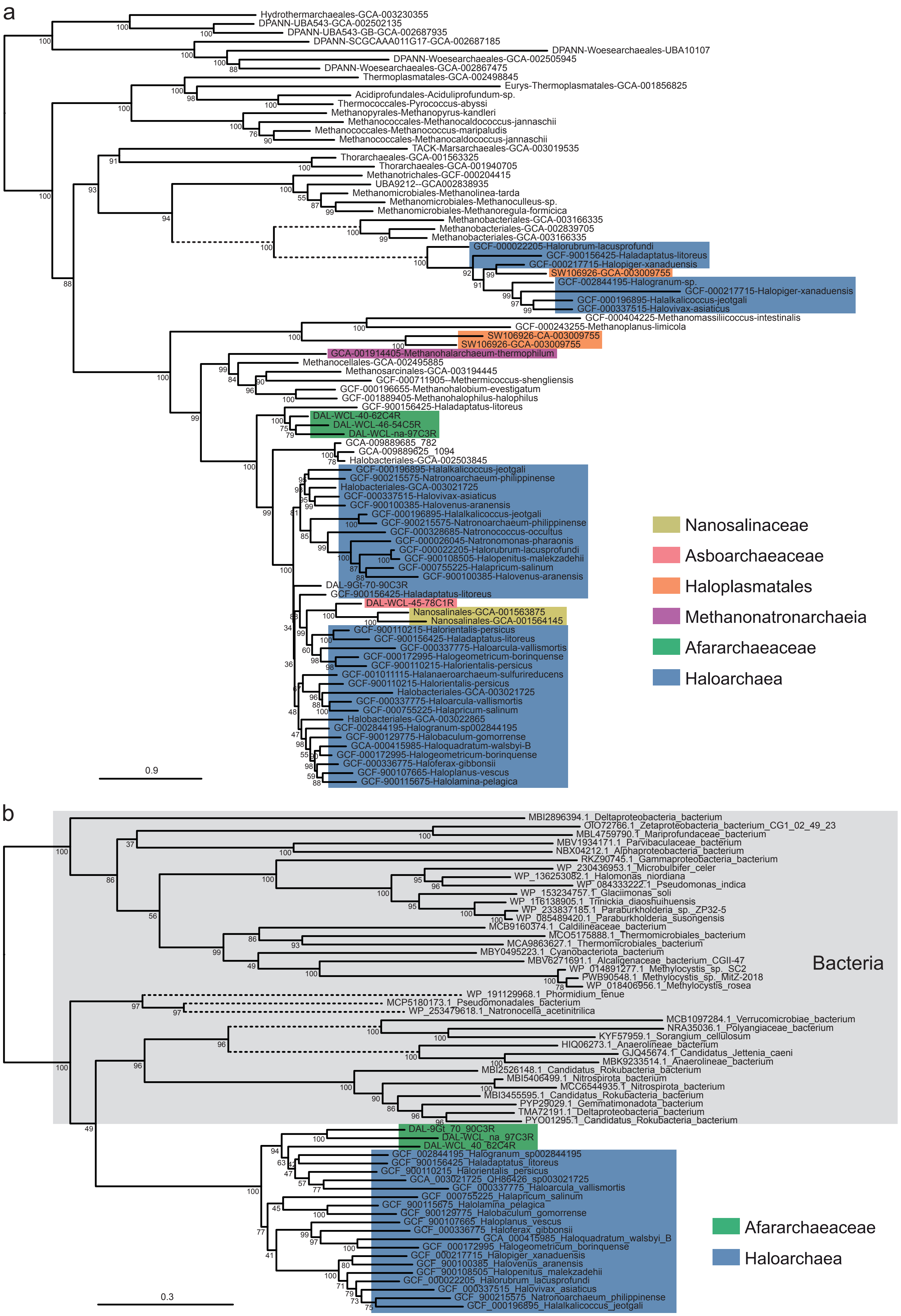


**Extended Data Fig. 10 | Maximum likelihood trees showing cases of horizontal gene transfer involving archaeal halophilic lineages. (a)** NhaP-type Na+/H+ and K+/H+ antiporters. **(b)** choline dehydrogenase BetA. The trees were constructed with the LG+C60+G+F model. Dashed branches have been shortened to half of their actual length. Scale bar indicates the expected average number of substitutions per site.

**Supplementary Table 1 |** Characteristics of the new MAGs included in this study.

| MAG ID | Classification | Origin | Completness (%) | Redundancy (%) | GC (%) | MAG size (bp) | No. of contigs | N50 |
|---|---|---|---|---|---|---|---|---|
| DAL-WCL_45_84C1R | Asboarchaeaceae | Western Canyon Lakes (WCL) | 83,95 | 1,23 | 61,15 | 1 201 103 | 81 | 40 248 |
| DAL-WCL_45_78C1R | Asboarchaeaceae | Western Canyon Lakes (WCL) | 78,4 | 0,62 | 57,47 | 1 454 511 | 344 | 5 895 |
| DAL-WCL_25_59C5R | Asboarchaeaceae | Western Canyon Lakes (WCL) | 59,26 | 4,94 | 56,82 | 599 589 | 232 | 2 980 |
| DAL-WCL_22_55C1R | Asboarchaeaceae | Western Canyon Lakes (WCL) | 54,94 | 1,23 | 58,67 | 470 854 | 26 | 26 678 |
| DAL-9Gt_90_72C1R | Asboarchaeaceae | La Grotte (9Gt) | 72,37 | 1,32 | 46,42 | 1 119 684 | 191 | 10104 |
| DAL-9Gt_91_54C5R | Asboarchaeaceae | La Grotte (9Gt) | 53,95 | 5,26 | 62,85 | 627 488 | 151 | 4 630 |
| DAL-9Gt_91_54C12R | Asboarchaeaceae | La Grotte (9Gt) | 53,94 | 11,84 | 64,33 | 643 450 | 102 | 8221 |
| DAL-Ass_51_71C0R | Asboarchaeaceae | Lake Assale (Ass) | 70,99 | 0 | 62 | 568 488 | 42 | 16 988 |
| DAL-Ass_36_69C3R | Asboarchaeaceae | Lake Assale (Ass) | 69,14 | 3,09 | 66,65 | 702 166 | 147 | 6 033 |
| DAL-WCL_na_97C3R | Afararchaeaceae | Western Canyon Lakes (WCL) | 96,91 | 3,09 | 54,9 | 1 895 979 | 117 | 39 046 |
| DAL-WCL_43_77C4R | Afararchaeaceae | Western Canyon Lakes (WCL) | 76,54 | 4,32 | 59,6 | 1 336 050 | 515 | 3 037 |
| DAL-WCL_40_62C4R | Afararchaeaceae | Western Canyon Lakes (WCL) | 61,73 | 3,7 | 54,82 | 1 704 136 | 724 | 2 656 |
| DAL-WCL_46_54C5R | Afararchaeaceae | Western Canyon Lakes (WCL) | 54,32 | 4,94 | 53,24 | 1 373 152 | 186 | 9 444 |
| DAL-9Gt_70_90C3R | Chewarchaeaceae | La Grotte (9Gt) | 89,47 | 2,63 | 61,52 | 2 933 771 | 132 | 38 607 |

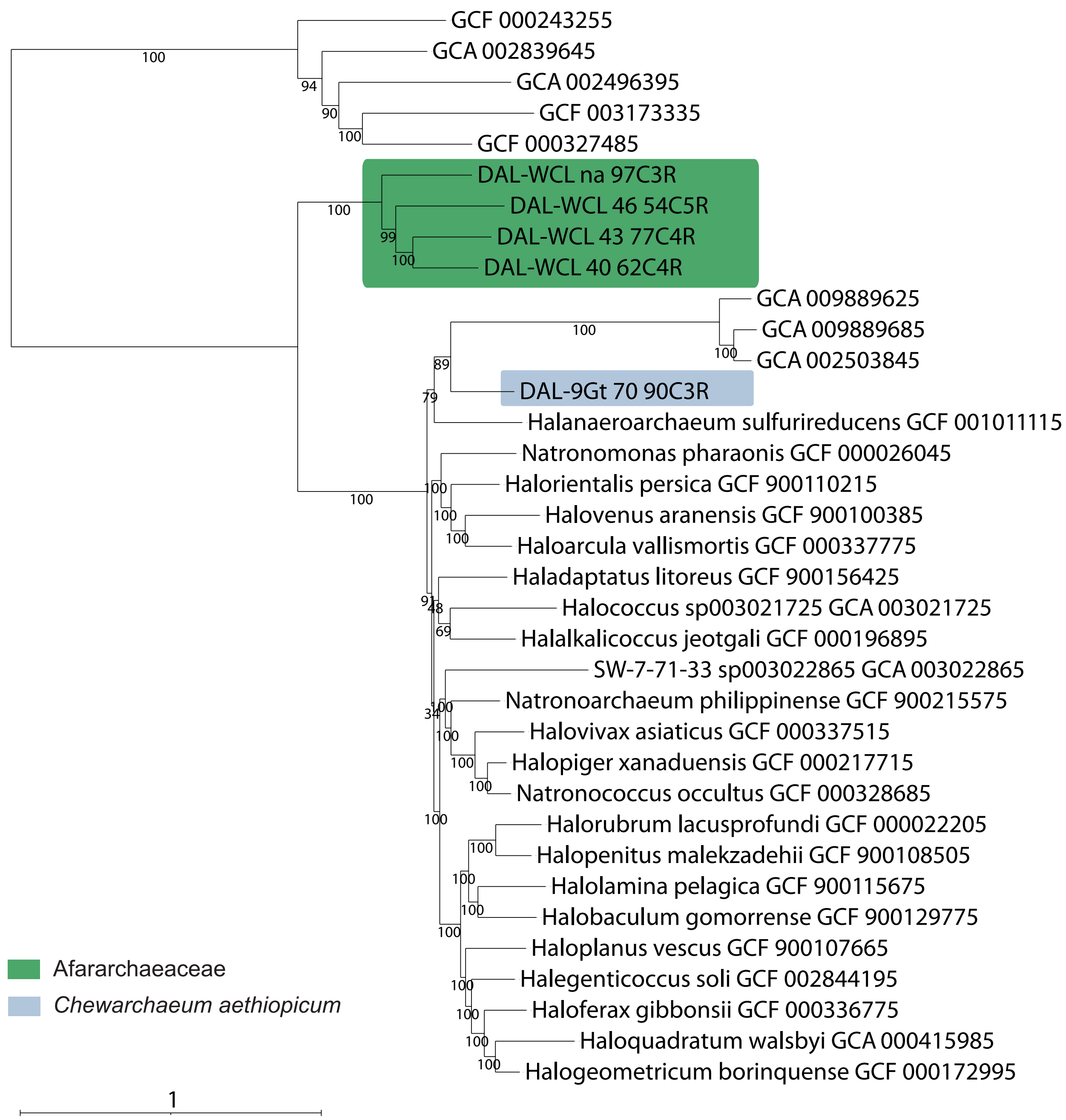


**Supplementary Figure 1 | Maximum likelihood phylogeny of 35 archaeal taxa based on the concatenated alignment of 122 single-copy proteins obtained from the Genome Taxonomy Database (GTDB).** The ML tree was constructed using the LG+C60+ F+Γ4 substitution model. Branch support was assessed using 1000 ultrafast bootstraps. The scale bar represents the estimated number of substitutions per site. Each tip contains a GTDB identification label.

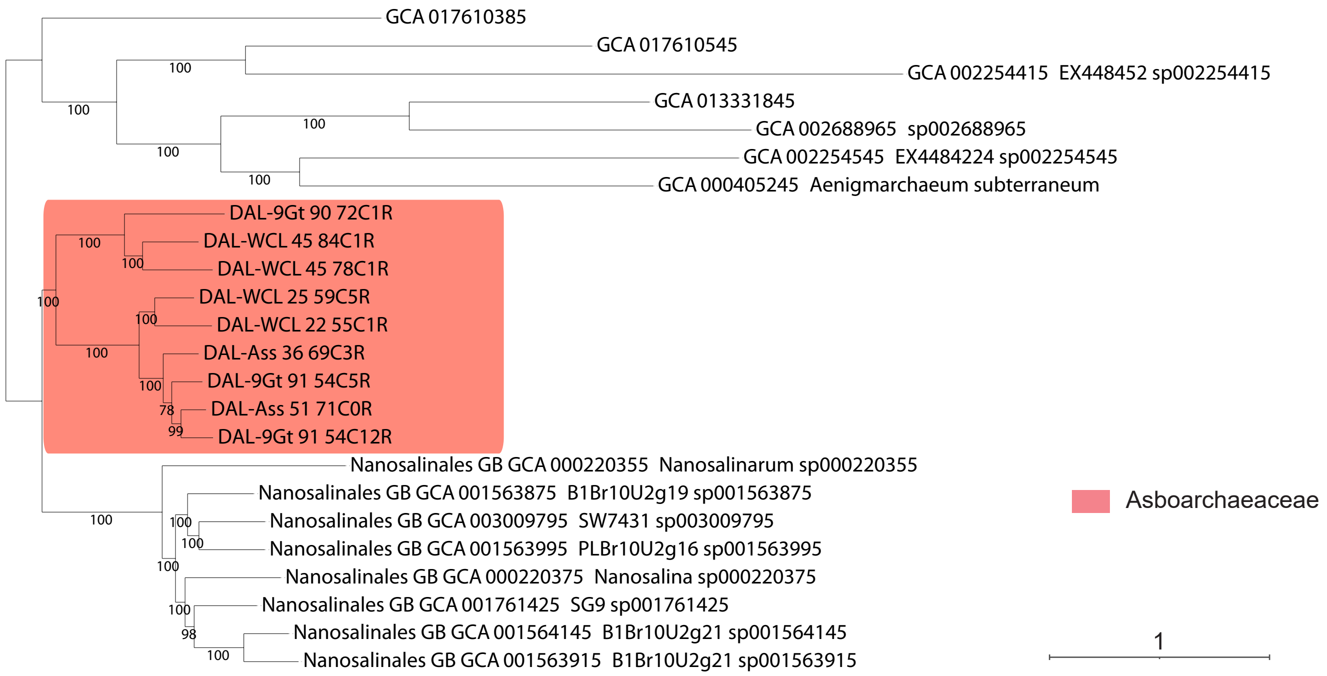


**Supplementary Figure 2 | Maximum likelihood phylogeny of 24 DPANN archaea based on the concatenated alignment of 99 single-copy proteins obtained from the Genome Taxonomy Database (GTDB).** The ML tree was constructed using the LG+C60+F+Γ4 substitution model of evolution. Branch support was assessed using 1000 ultrafast bootstraps. The scale bar represents the estimated number of substitutions per site. Each tip contains a GTDB identification label, in addition to the Latin name, when avalable.

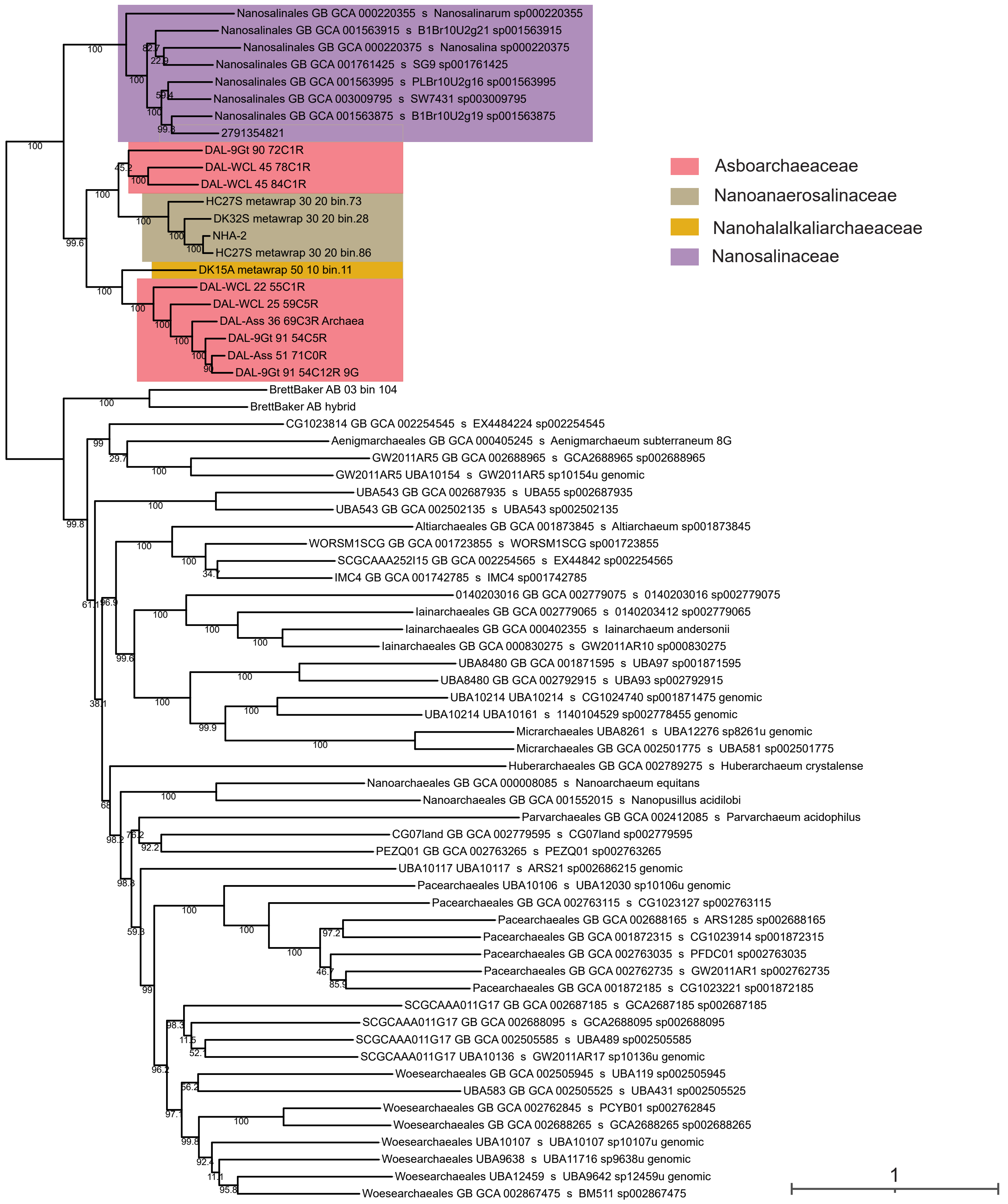


**Supplementary Figure 3 | Maximum likelihood phylogeny of 70 DPANN archaea based on the concatenated alignment of 24 large subunit ribosomal proteins.** The ML tree was constructed using the LG+C20+Γ4 substitution model. Branch support was assessed using 1000 ultrafast bootstraps. The scale bar represents the estimated number of substitutions per site. Each tip label shows the GTDB taxonomic order, accession number, and species identifier.

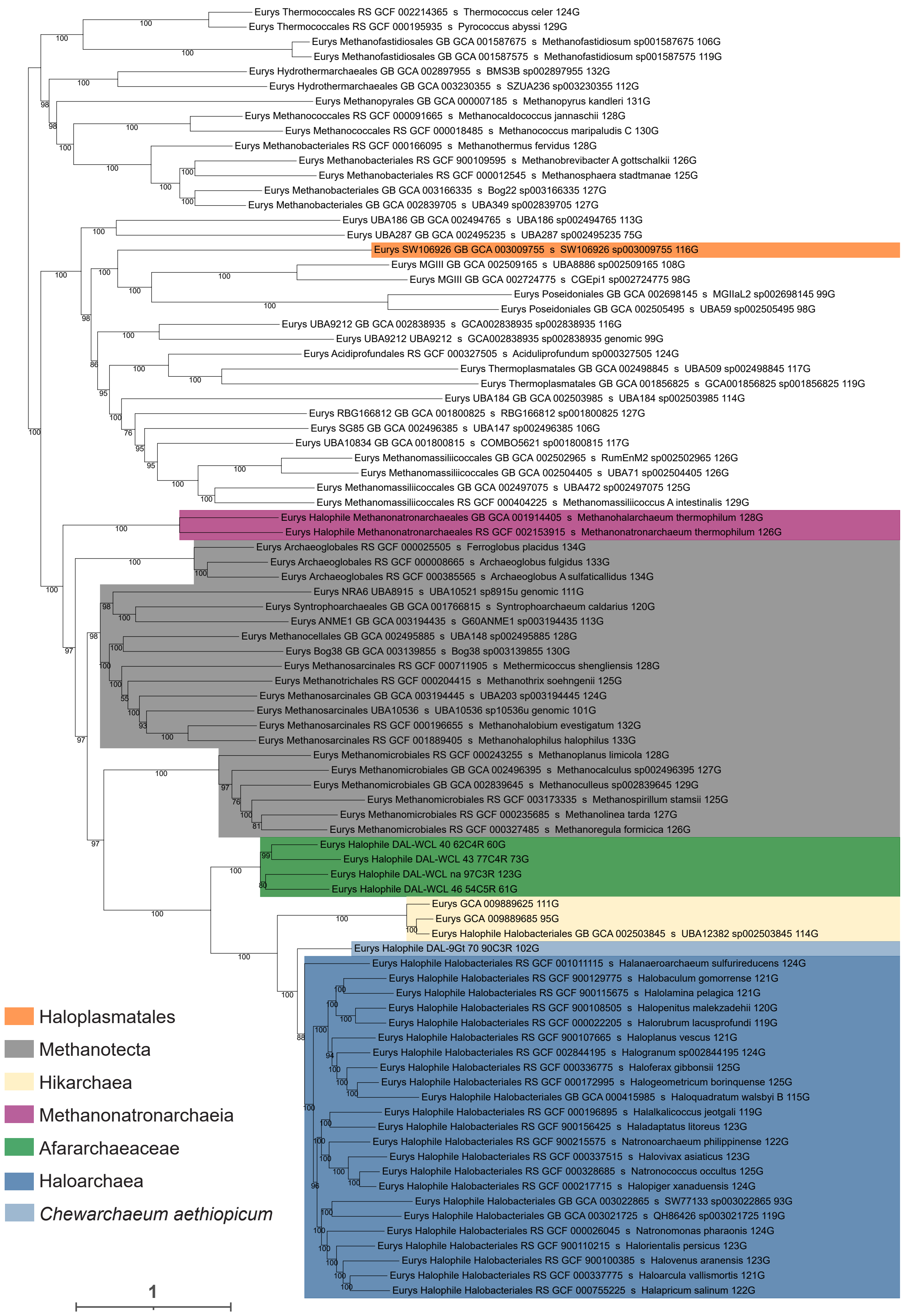


**Supplementary Figure 4 | Maximum likelihood phylogeny of 87 archaeal taxa based on the concatenated alignment of 136 new marker (NM) protein dataset.** The ML tree was constructed using the LG+C60+F+Γ4 substitution model. Branch support was assessed using 1000 ultrafast bootstraps. The scale bar represents the estimated number of substitutions per site. Each tip label provides information about the taxonomic order based on GTDB, GTDB accession number, species identification, and the number of markers identified for each taxon out of the 136 NM proteins.

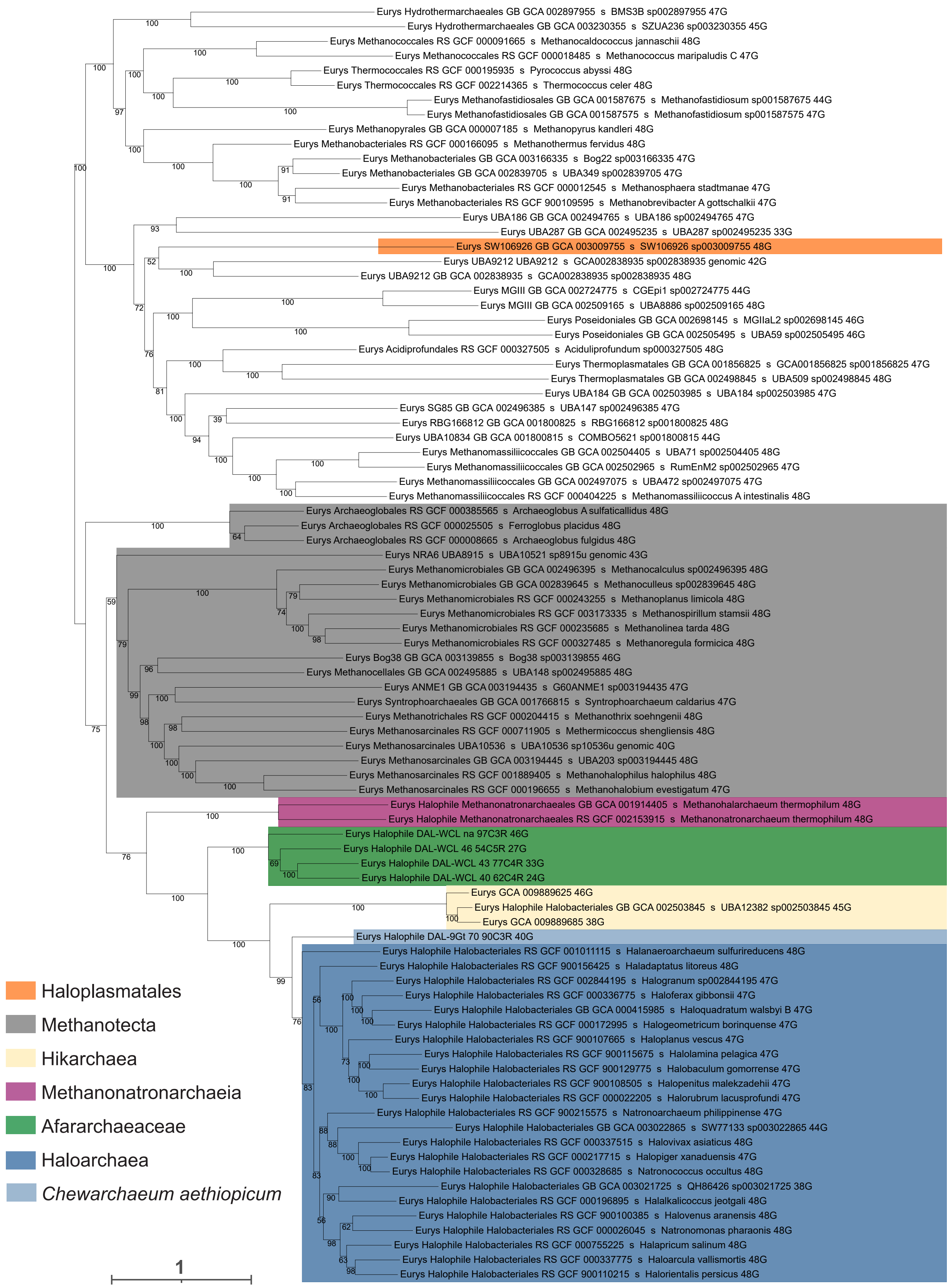


**Supplementary Figure 5 | Maximum likelihood phylogeny of 87 archaeal taxa based on the concatenated alignment of 48 ribosomal protein (RP) dataset.** The ML tree was constructed using the LG+C60+F+Γ4 substitution model. Branch support was assessed using 1000 ultrafast bootstraps. The scale bar represents the estimated number of substitutions per site. Each tip label provides information about the taxonomic order based on GTDB, GTDB accession number, species identification, and the number of markers identified for each taxon out of the 48 RP proteins.

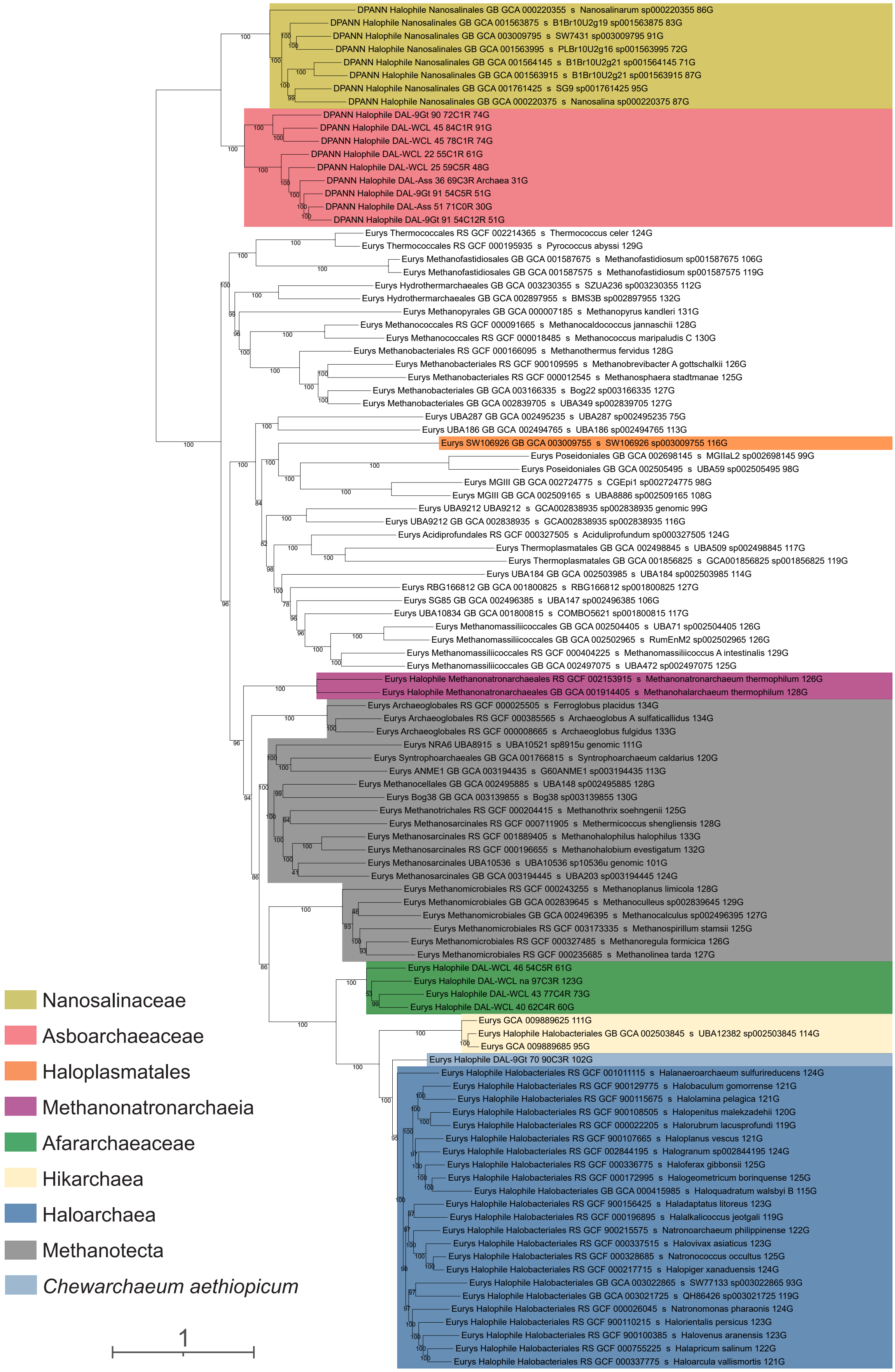


**Supplementary Figure 6 | Maximum likelihood phylogeny of 104 archaeal taxa based on the concatenated alignment of 136 new marker (NM) protein dataset.** The ML tree was constructed using the LG + C60 + F + Γ4 substitution model. Branch support was assessed using 1000 ultrafast bootstraps. The scale bar represents the estimated number of substitutions per site. Each tip label provides information about the taxonomic order based on GTDB, GTDB accession number, species identification, and the number of markers identified for each taxon out of the 136 NM proteins.

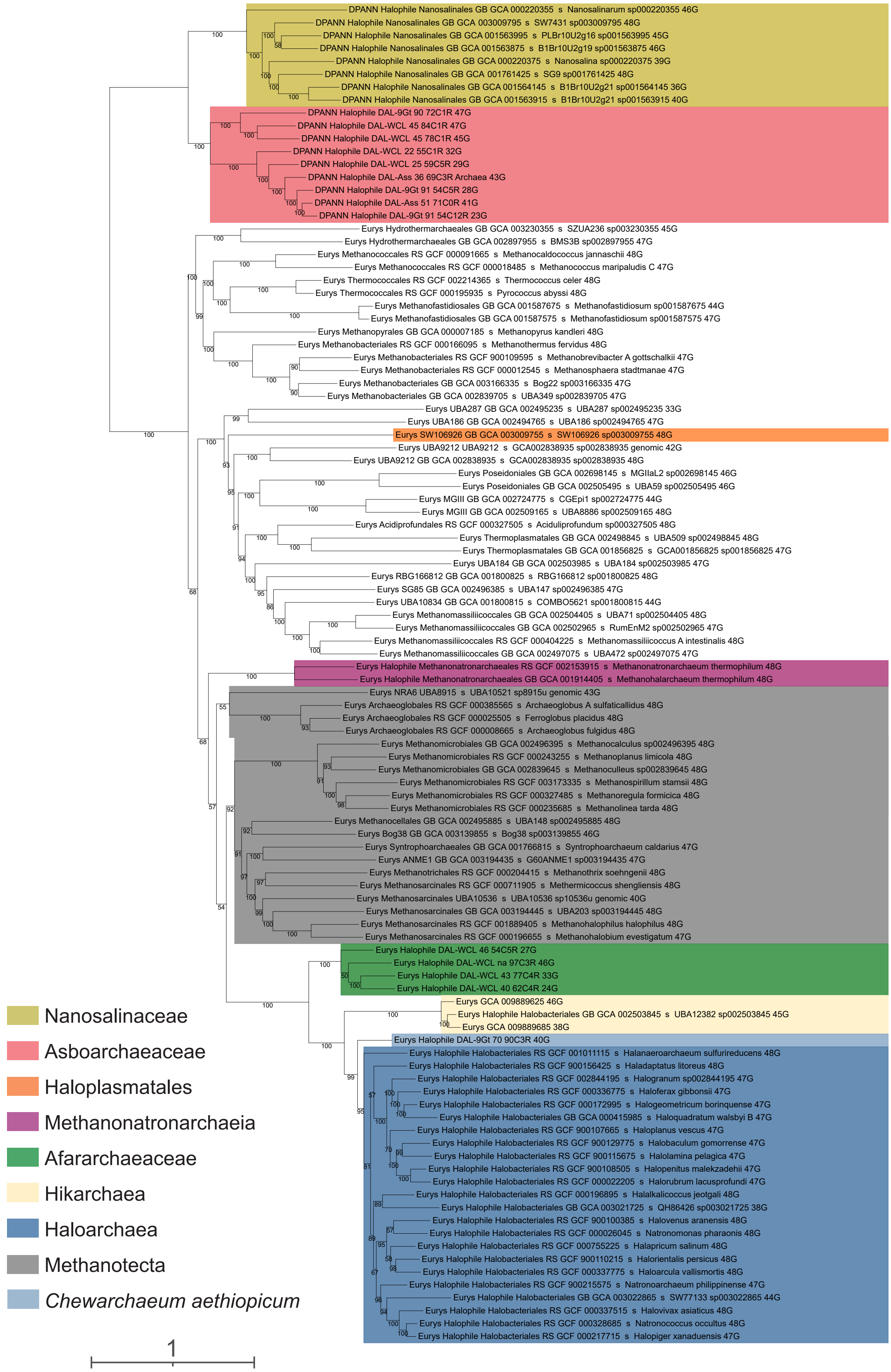


**Supplementary Figure 7 | Maximum likelihood phylogeny of 104 archaeal taxa based on the concatenated alignment of 48 ribosomal protein (RP) dataset.** The ML tree was constructed using the LG + C60 + F + Γ4 substitution model. Branch support was assessed using 1000 ultrafast bootstraps. The scale bar represents the estimated number of substitutions per site. Each tip label provides information about the taxonomic order based on GTDB, GTDB accession number, species identification, and the number of markers identified for each taxon out of the 48 RP proteins.

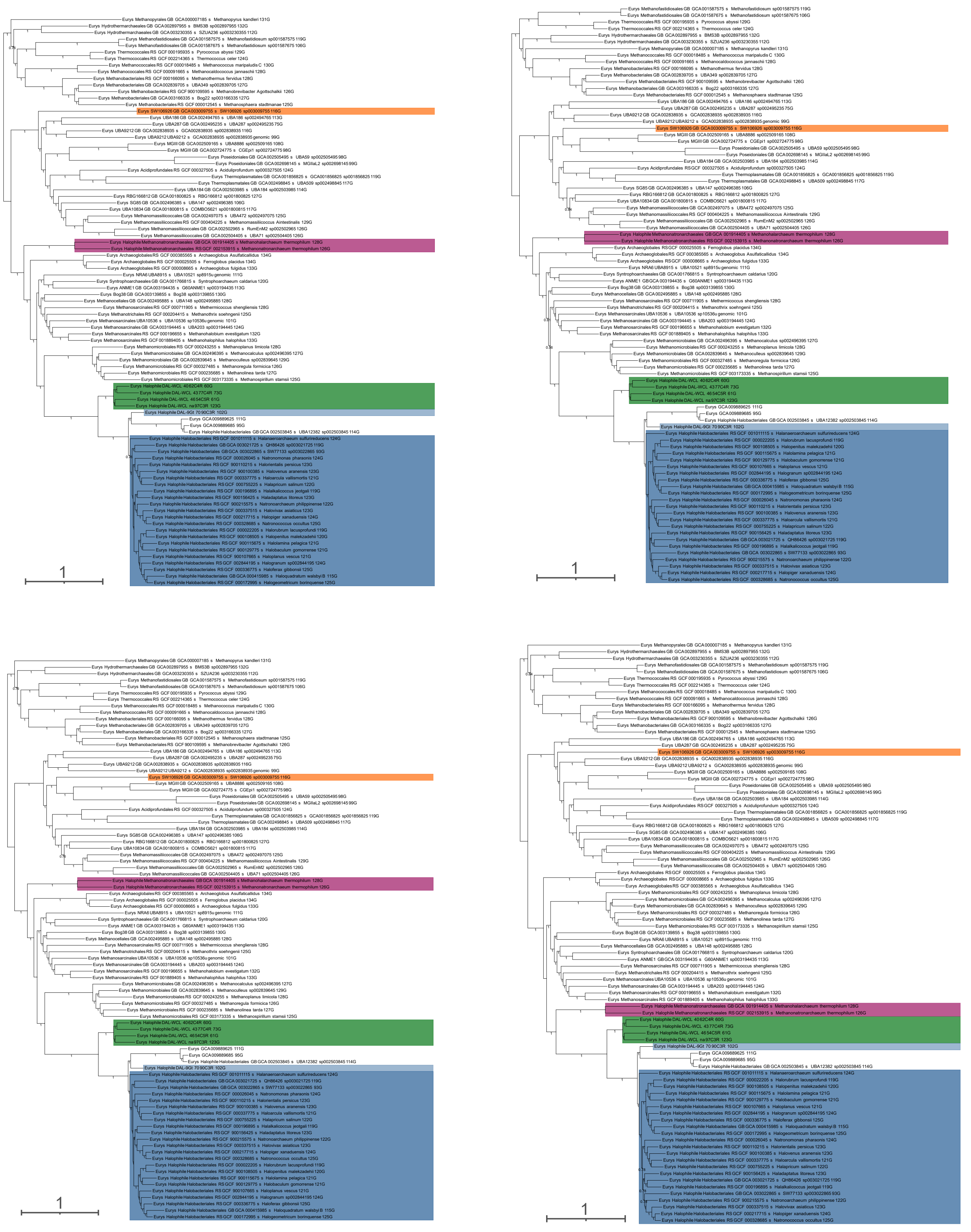


**Supplementary Figure 8 | Consensus tree for each of the four MCMC chains for the 87-NM dataset.** Four independent Markov chain Monte Carlo (MCMC) chains were inferred using PhyloBayes (CAT+GTR, 15,000 generations with a burn-in of 3,000 generations). Support at branches corresponds to posterior probabilities estimated post-burnin. The scale bar represents the estimated number of substitutions per site. Different halophilic clades are visually represented with distinct colors: haloplasmatales (orange), methanonatronarchaea (purple), afararchaea (green), and haloarchaea (blue). Each tip label provides information about the archaeal supergroup, taxonomic order based on GTDB, GTDB accession number, species identification, and the number of markers identified for each taxon out of the 136 NM proteins.

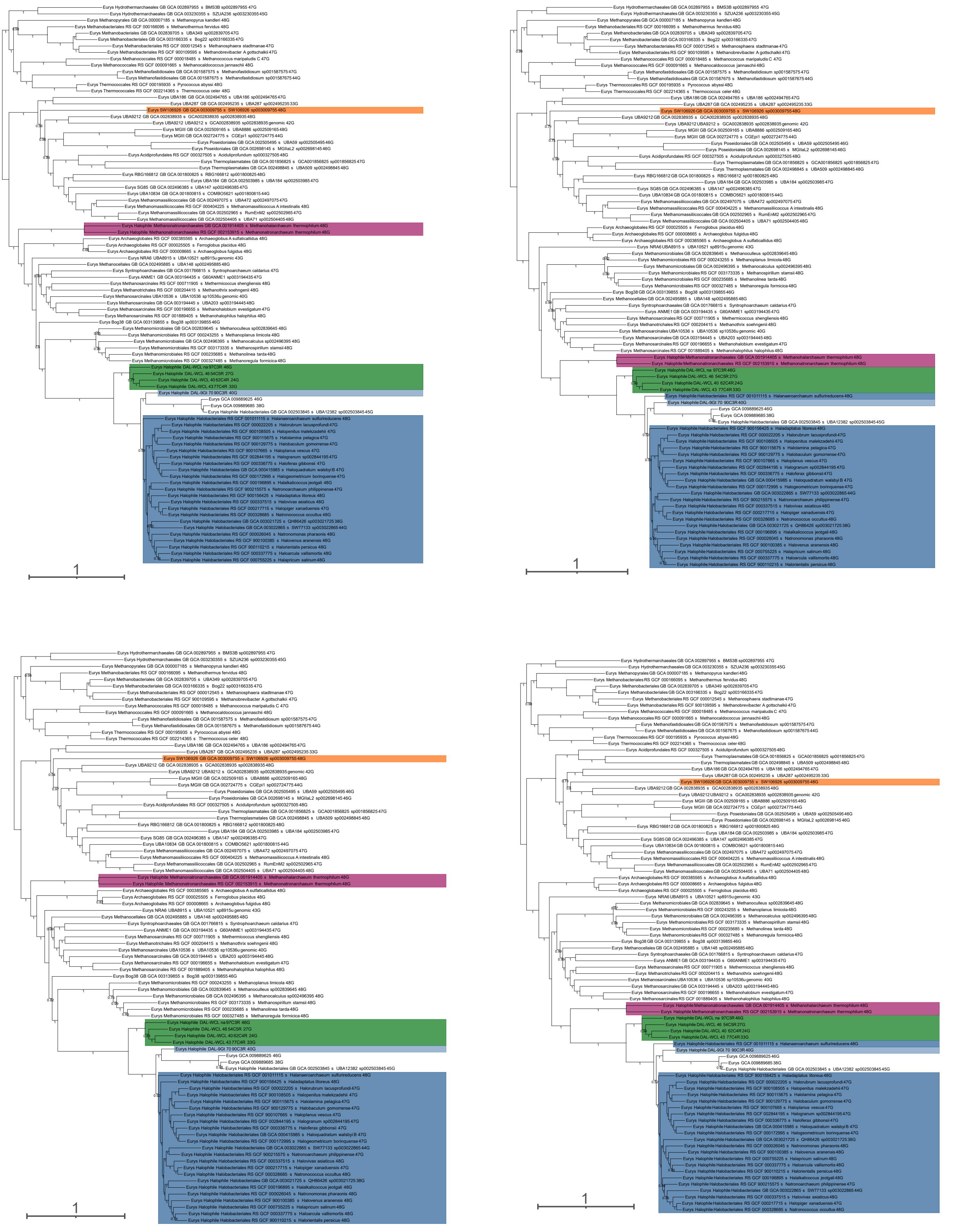

**Supplementary Figure 9 | Consensus tree for each of the four MCMC chains for the 87-RP dataset.** Four independent Markov chain Monte Carlo (MCMC) chains were inferred using PhyloBayes (CAT+GTR, 15,000 generations with a burn-in of 3,000 generations). Support at branches corresponds to posterior probabilities estimated post-burnin. The scale bar represents the estimated number of substitutions per site. Different halophilic clades are visually represented with distinct colors: haloplasmatales (orange), methanonatronarchaea (purple), afararchaea (green), and haloarchaea (blue). Each tip label provides information about the archaeal super group, taxonomic order based on GTDB, GTDB accession number, species identification, and the number of markers identified for each taxon out of the 48 RP proteins.

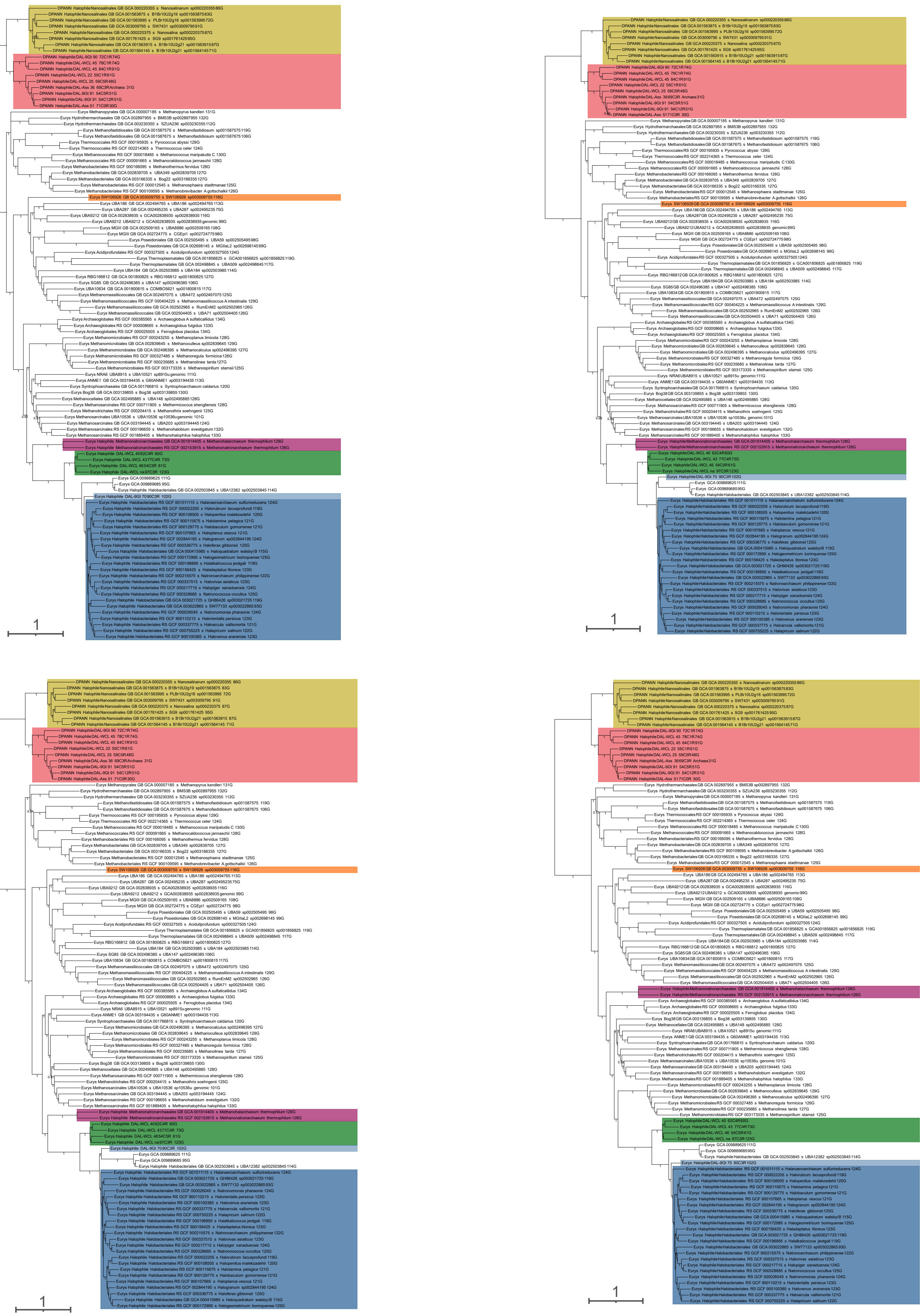

**Supplementary Figure 10 | Consensus tree for each of the four MCMC chains for the 104-NM dataset.** Four independent Markov chain Monte Carlo (MCMC) chains were inferred using PhyloBayes (CAT+GTR, 15,000 generations with a burn-in of 3,000 generations). Support at branches corresponds to posterior probabilities estimated post-burnin. The scale bar represents the estimated number of substitutions per site. Different halophilic clades are visually represented with distinct colors: haloplasmatales (orange), methanonatronarchaea (purple), afararchaea (green), and haloarchaea (blue). Each tip label provides information about the archaeal supergroup, taxonomic order based on GTDB, GTDB accession number, species identification, and the number of markers identified for each taxon out of the 136 NM proteins.

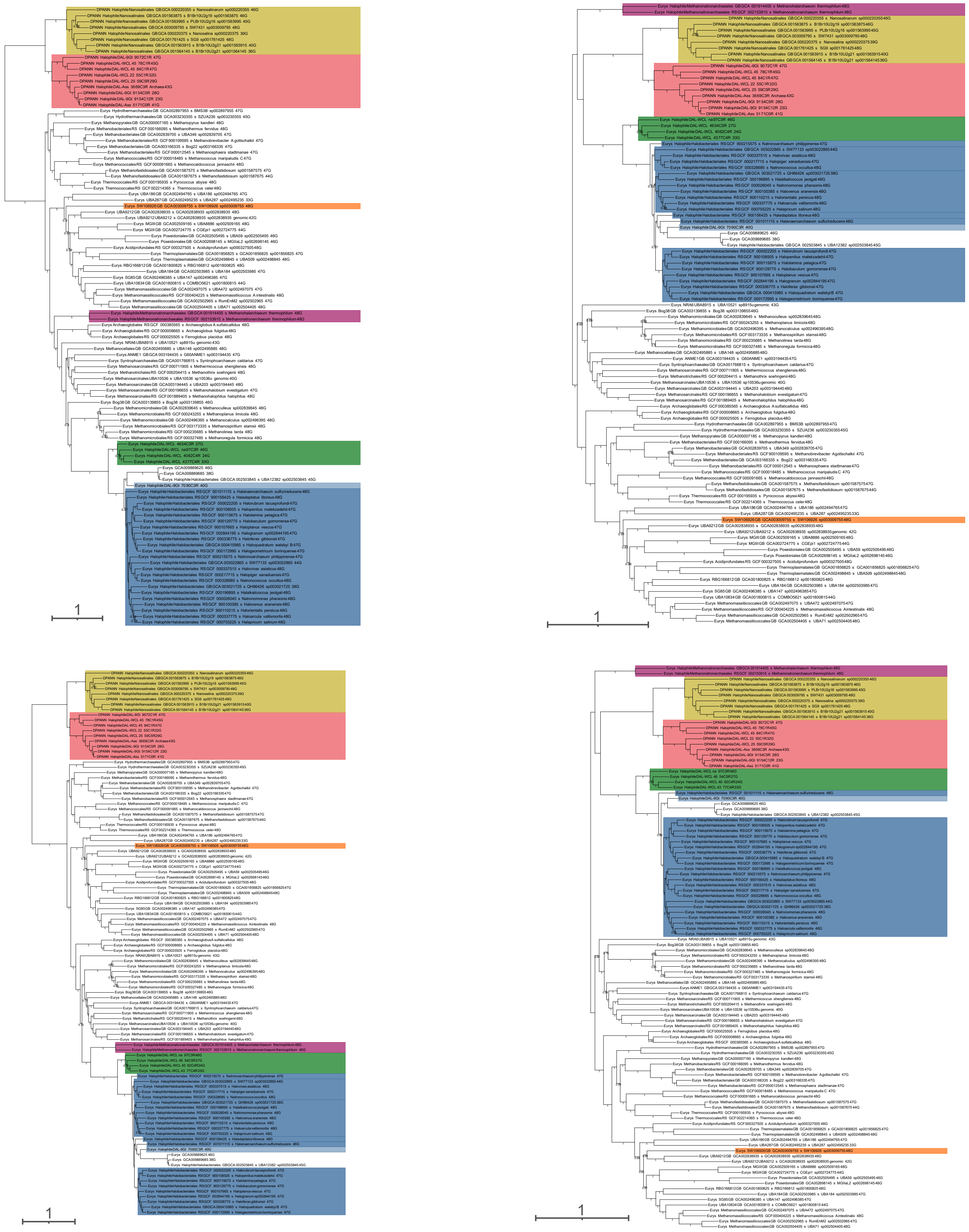

**Supplementary Figure 11 | Consensus tree for each of the four MCMC chains for the 104-RP dataset.** Four independent Markov chain Monte Carlo (MCMC) chains were inferred using PhyloBayes (CAT+GTR, 15,000 generations with a burn-in of 3,000 generations). Support at branches corresponds to posterior probabilities estimated post-burnin. The scale bar represents the estimated number of substitutions per site. Different halophilic clades are visually represented with distinct colors: haloplasmatales (orange), methanonatronarchaea (purple), afararchaea (green), and haloarchaea (blue). Each tip label provides information about the archaeal super-group, taxonomic order based on GTDB, GTDB accession number, species identification, and the number of markers identified for each taxon out of the 48 RP proteins.

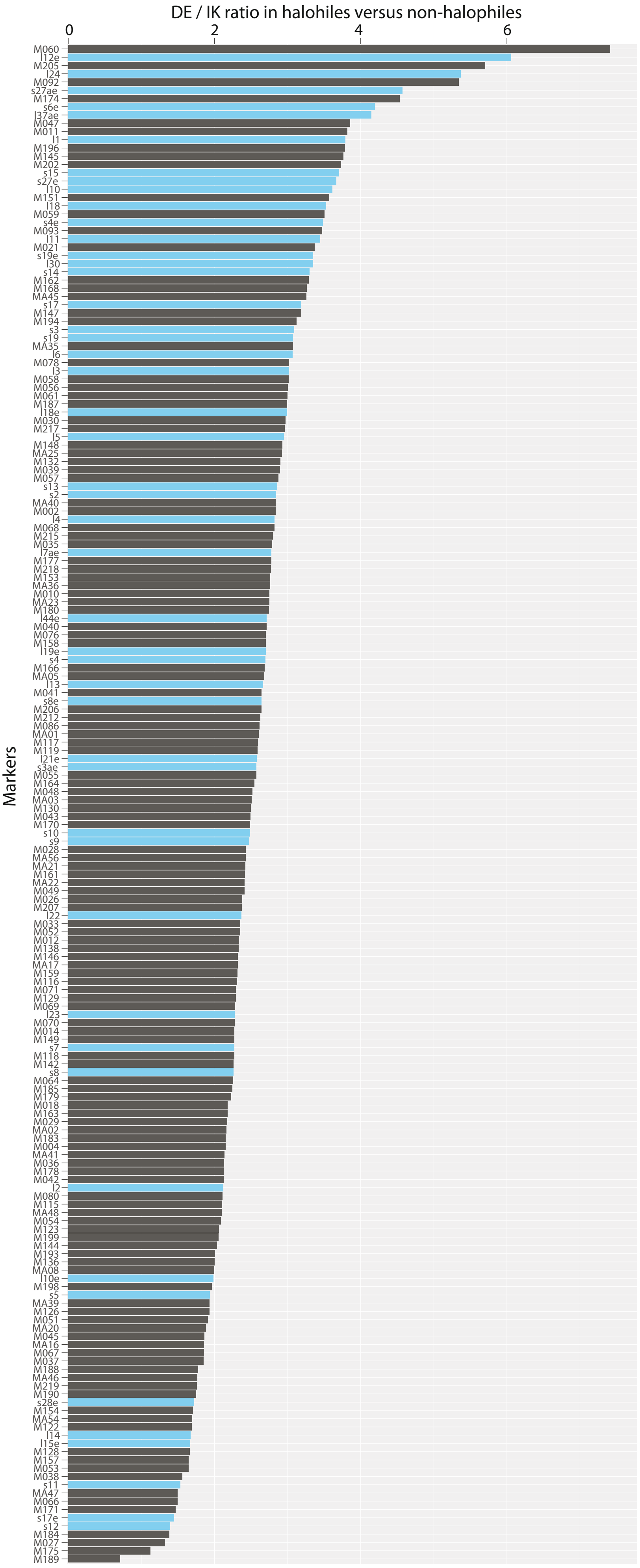


**Supplementary Figure 12 | RP (blue) and NM (black) markers ranked from the most to least biased based on the calculated DE/IK ratio of halophiles versus the DE/IK ratio of non-halophiles.**

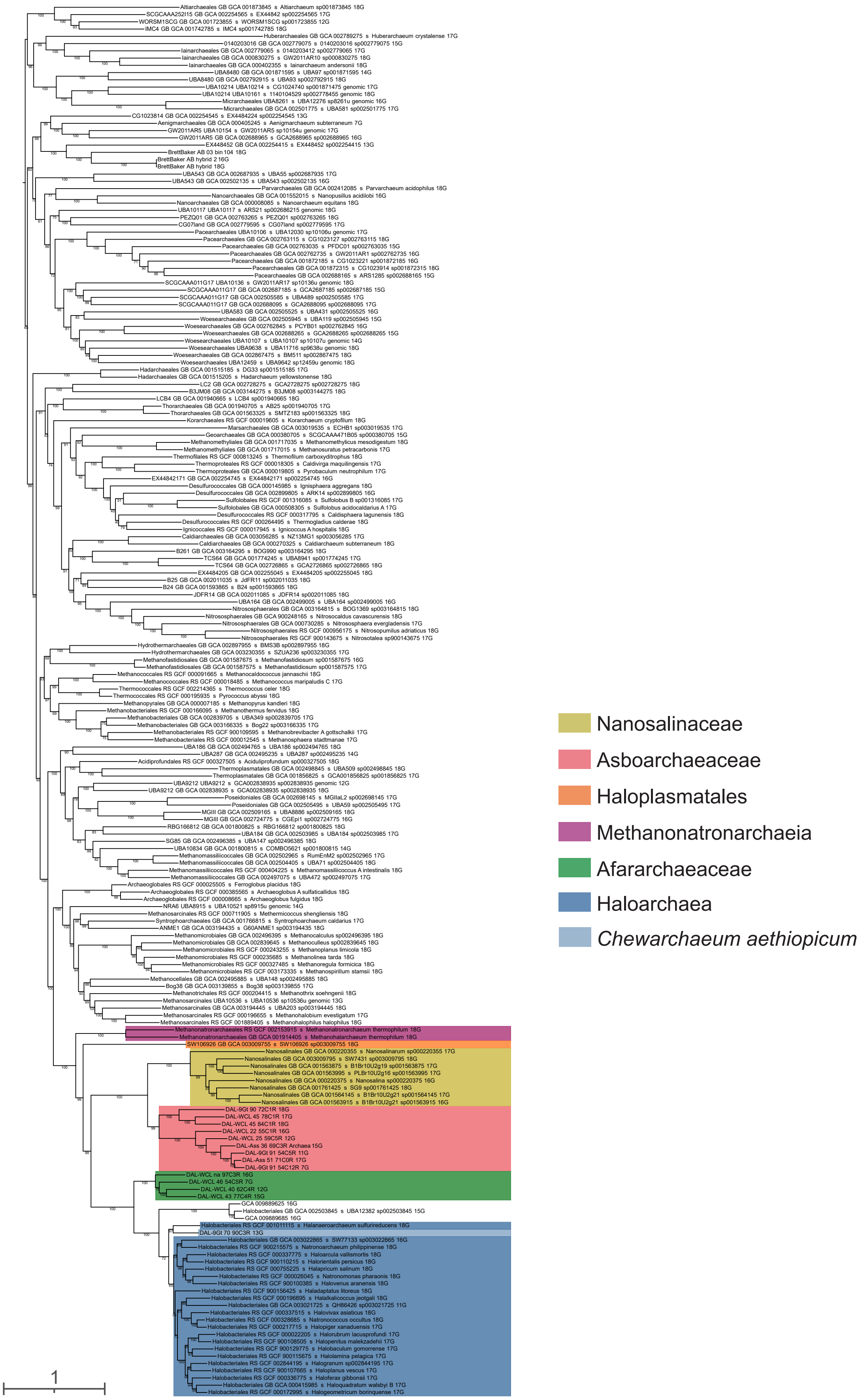


**Supplementary Figure 13 | Maximum likelihood phylogeny based on concatenating the 18 most compositionally biased ribosomal proteins.** The ML tree was constructed using the LG+C60+F+Γ4 substitution model of evolution. Branch support was assessed using 1000 ultrafast bootstraps. The scale bar represents the estimated number of substitutions per site. Each tip label provides information about the archaeal super-group, taxonomic order based on GTDB, GTDB accession number, species identification, and the number of markers identified for each taxon out of the 18 RP proteins.

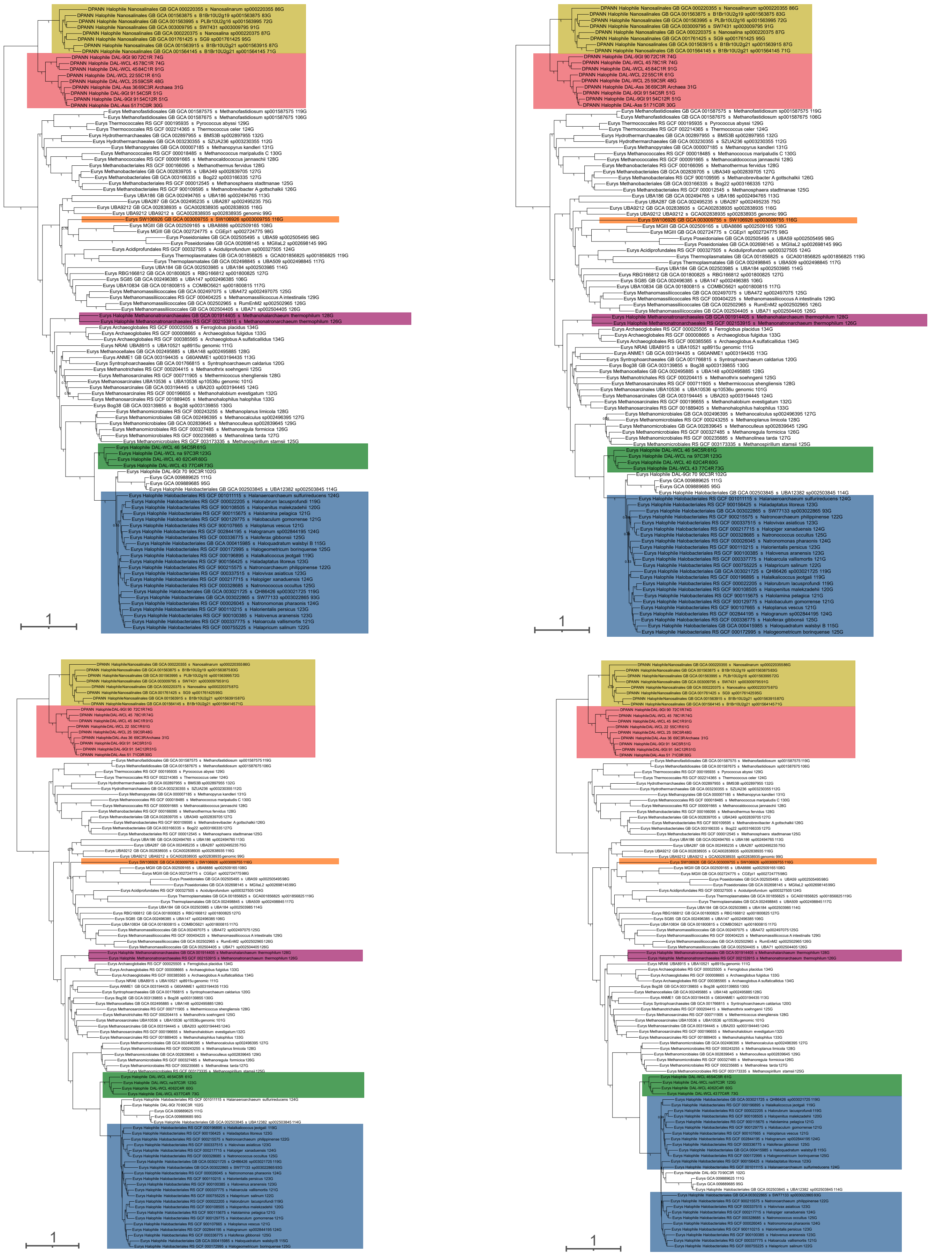

**Supplementary Figure 14 | Consensus tree for each of the four MCMC chains for the 104-RP dataset with 20% of most biased sites removed.** Four independent Markov chain Monte Carlo (MCMC) chains were inferred using PhyloBayes (CAT+GTR, 15,000 generations with a burn-in of 3,000 generations). Support at branches corresponds to posterior probabilities estimated post-burnin. The scale bar represents the estimated number of substitutions per site. Different halophilic clades are visually represented with distinct colors: haloplasmatales (orange), methanonatronarchaea (purple), afararchaea (green), and haloarchaea (blue). Each tip label provides information about the archaeal super group, taxonomic order based on GTDB, GTDB accession number, species identification, and the number of markers identified for each taxon out of the 48 RP proteins.

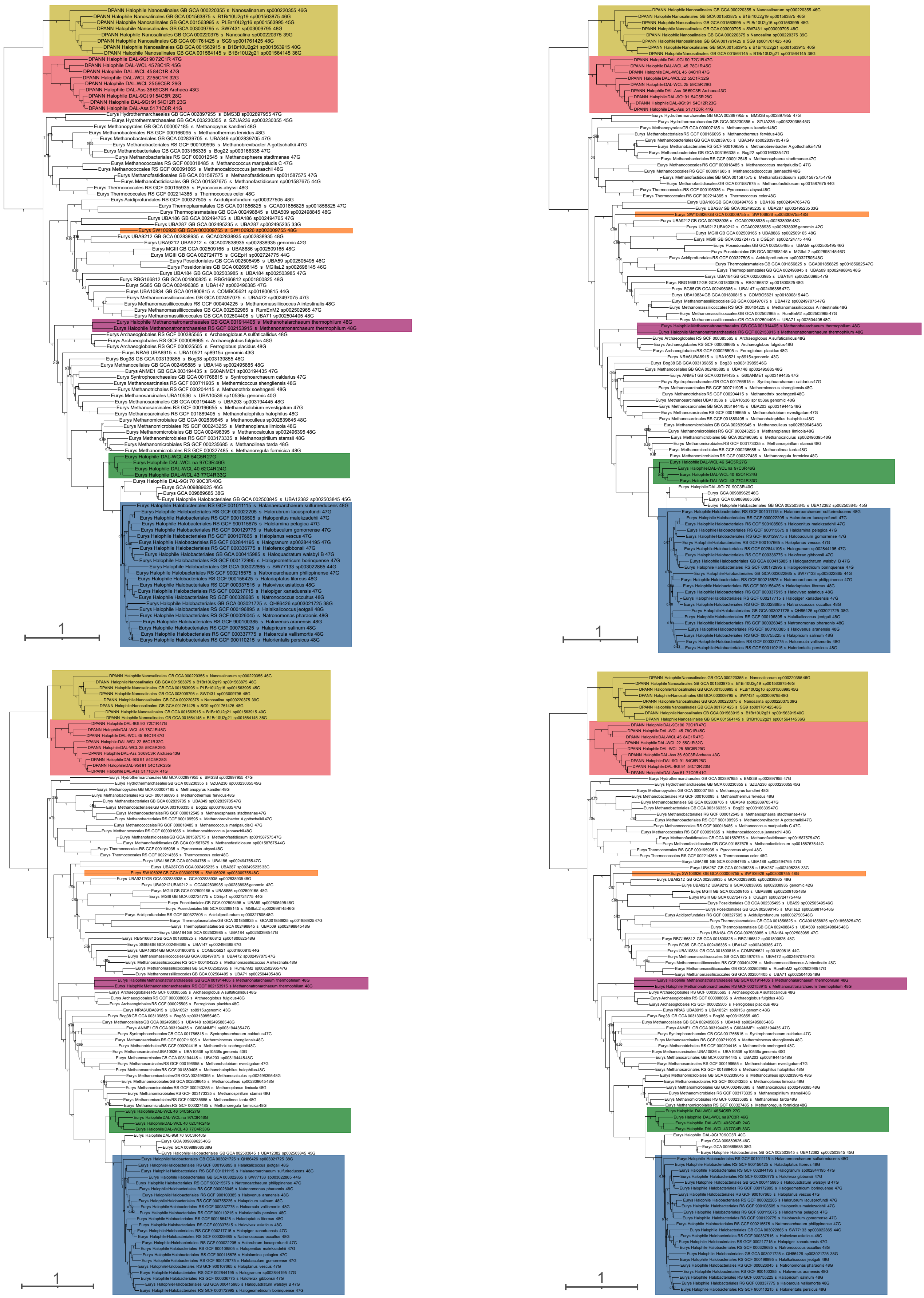

**Supplementary Figure 15 | Consensus tree for each of the four MCMC chains for the 104-NM dataset with 20% of most biased sites removed.** Four independent Markov chain Monte Carlo (MCMC) chains were inferred using PhyloBayes (CAT+GTR, 15,000 generations with a burn-in of 3,000 generations). Support at branches corresponds to posterior probabilities estimated post-burnin. The scale bar represents the estimated number of substitutions per site. Different halophilic clades are visually represented with distinct colors: haloplasmatales (orange), methanonatronarchaea (purple), afararchaea (green), and haloarchaea (blue). Each tip label provides information about the archaeal supergroup, taxonomic order based on GTDB, GTDB accession number, species identification, and the number of markers identified for each taxon out of the 136 NM proteins.

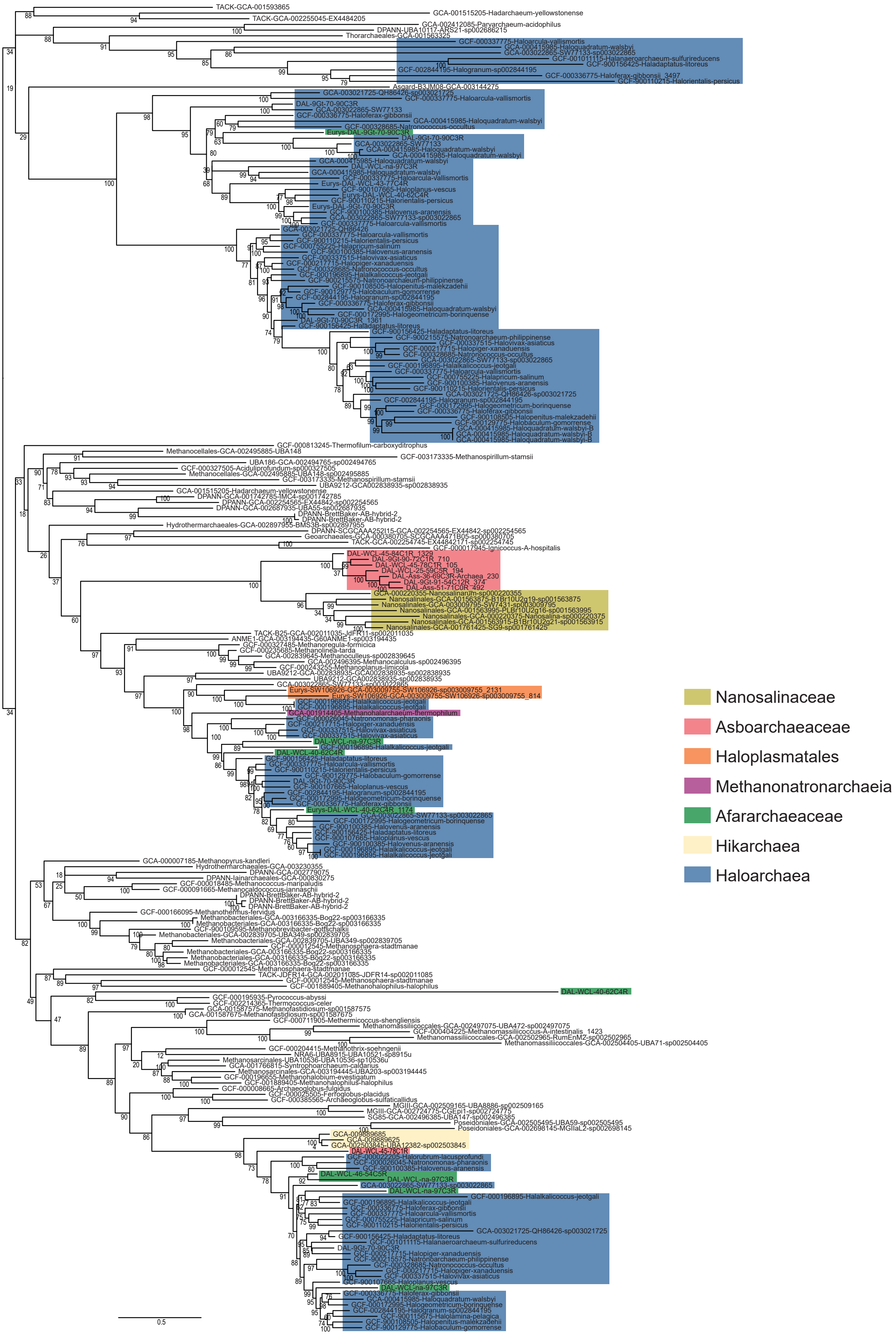


**Supplementary Fig. 16 | Maximum likelihood phylogenetic tree of TrkA K+ transporter**. The tree was constructed with the LG+C60+Γ4+F substitution model. Branch support was assessed using 1000 ultrafast bootstraps. The scale bar represents the estimated number of substitutions per site.

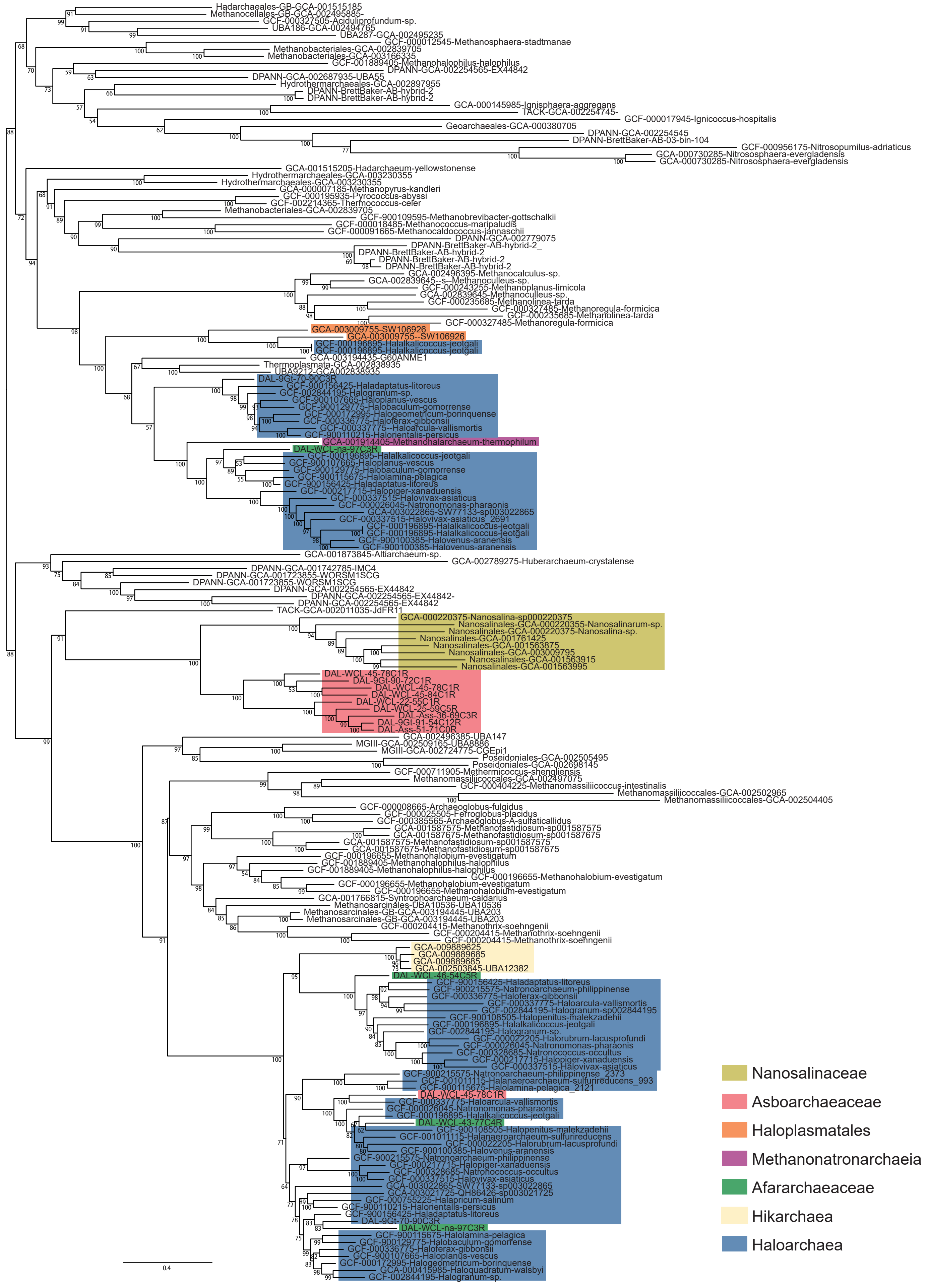


**Supplementary Fig. 17 | Maximum likelihood phylogenetic tree of TrkH K+ transporter**. The tree was constructed with the LG+C60+Γ4+F substitution model. Branch support was assessed using 1000 ultrafast bootstraps. The scale bar represents the estimated number of substitutions per site.

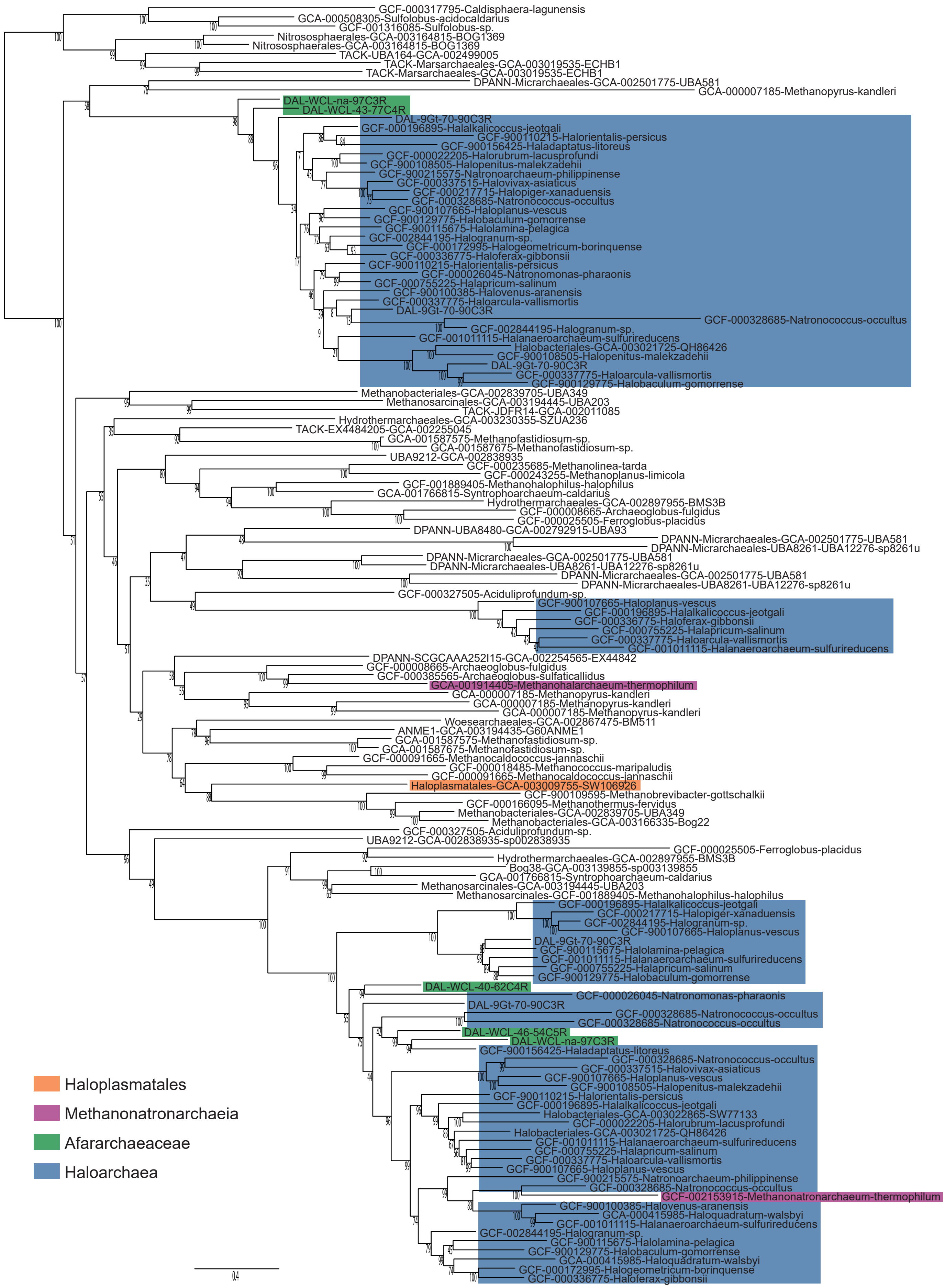


**Supplementary Fig. 18 | Maximum likelihood phylogenetic tree of TrkA-N K+ transporter**. The tree was constructed with the LG+C60+Γ4+F substitution model. Branch support was assessed using 1000 ultrafast bootstraps. The scale bar represents the estimated number of substitutions per site.

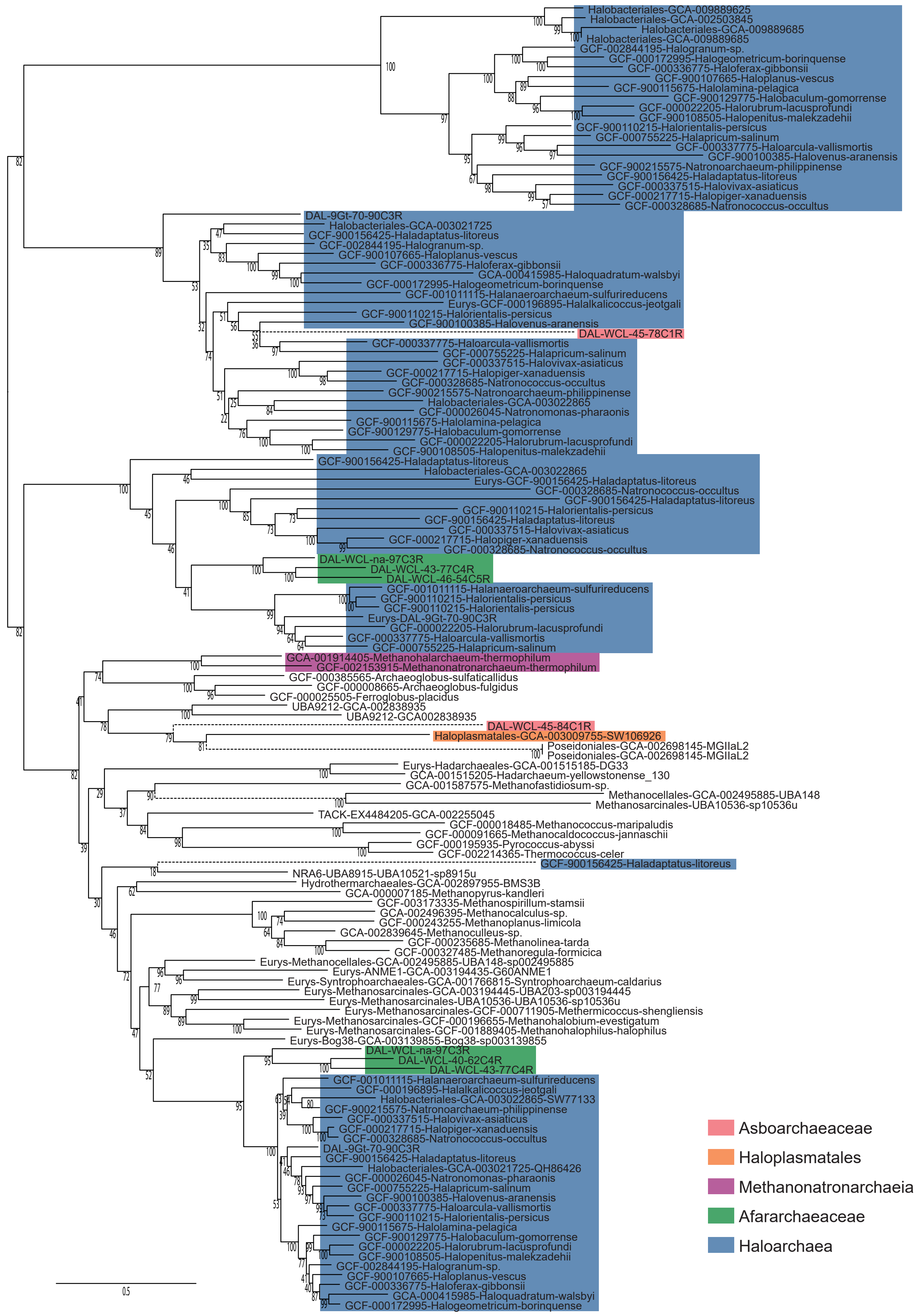


**Supplementary Fig. 19 | Maximum likelihood phylogenetic tree of Kef K+ transporters**. The tree was constructed with the LG+C60+Γ4+F substitution model. Branch support was assessed using 1000 ultrafast bootstraps. The scale bar represents the estimated number of substitutions per site.

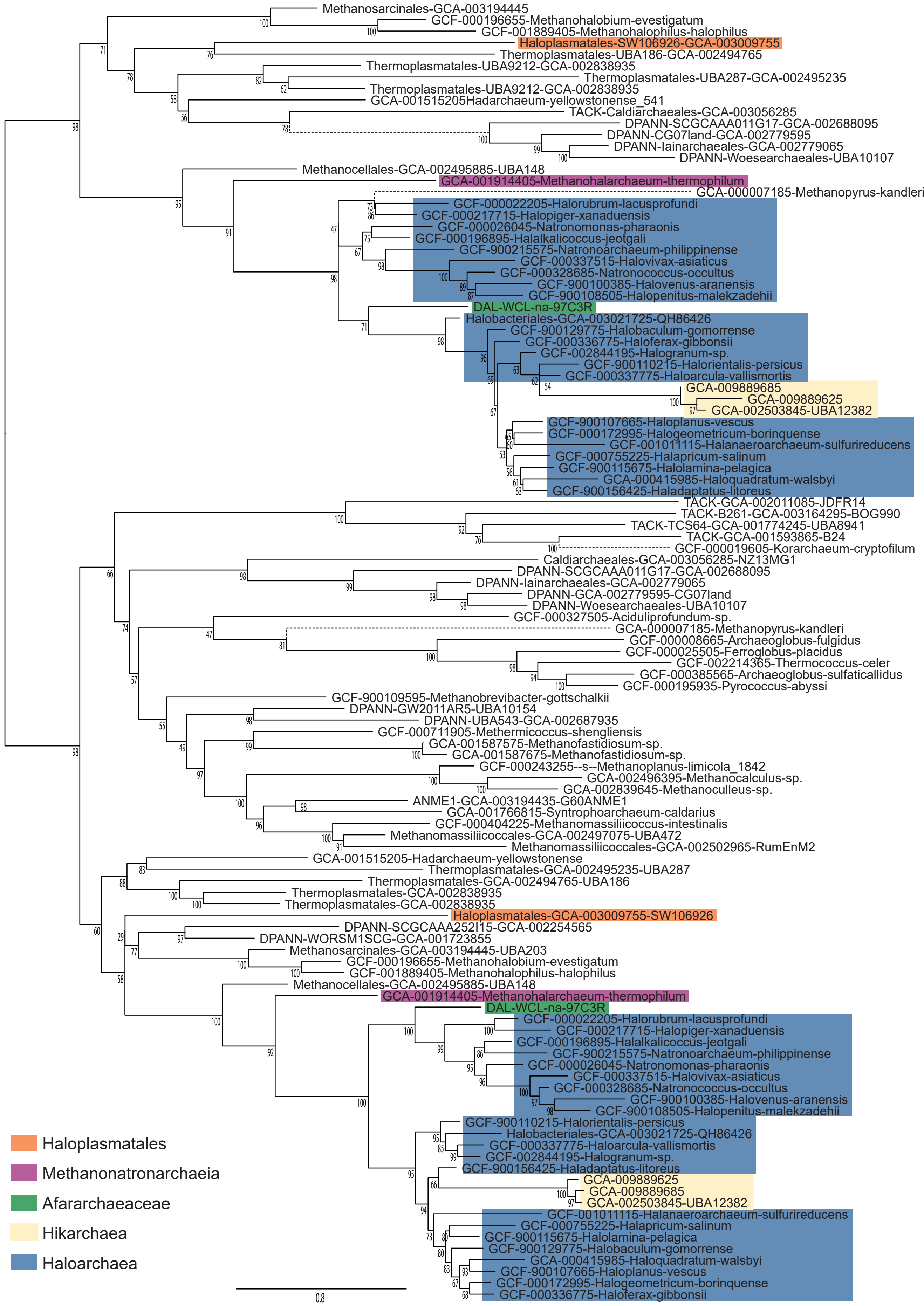


**Supplementary Fig. 20 | Maximum likelihood phylogenetic tree of Mg2+ transporters**. The tree was constructed with the LG+C60+Γ4+F substitution model. Branch support was assessed using 1000 ultrafast bootstraps. The scale bar represents the estimated number of substitutions per site.

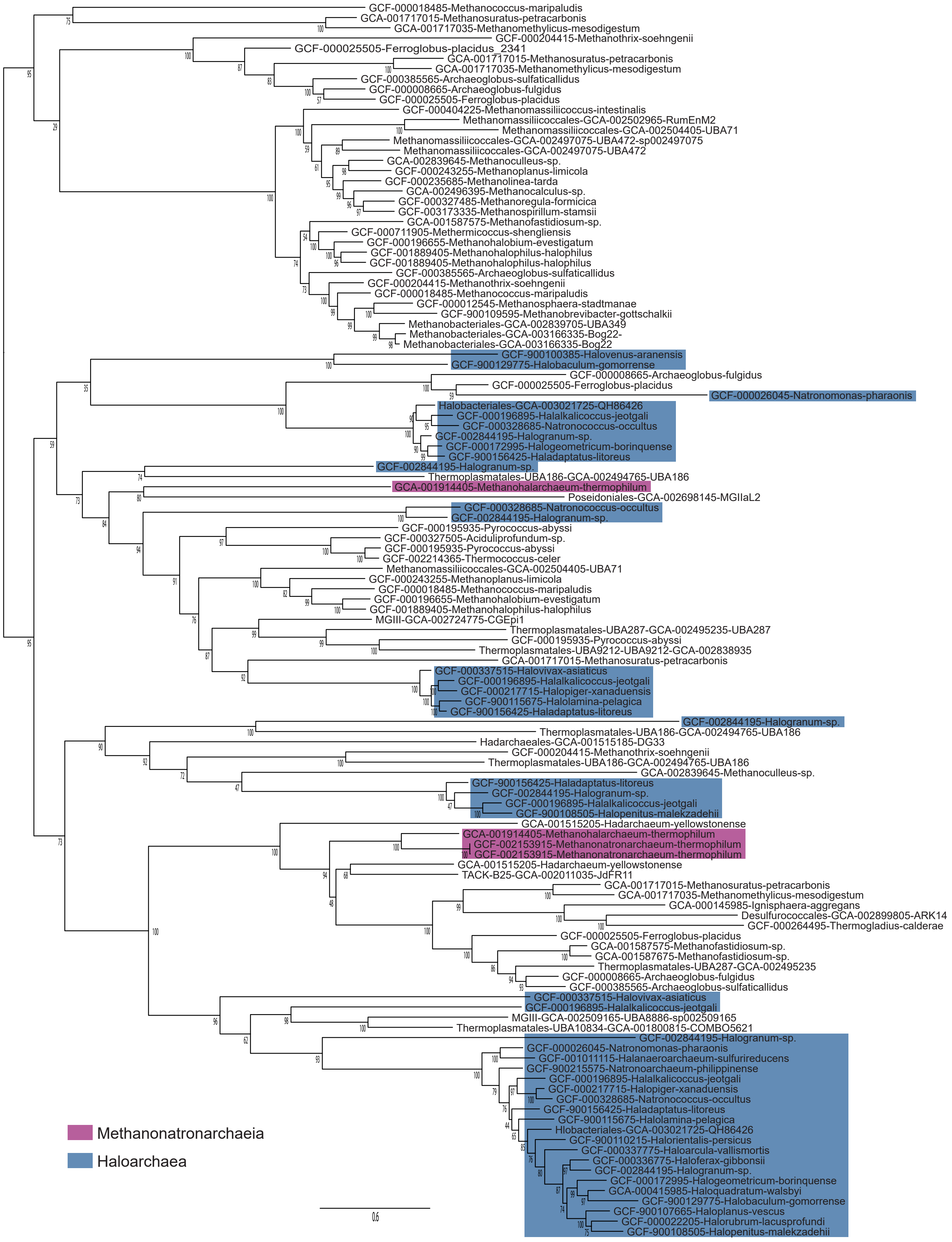


**Supplementary Fig. 21 | Maximum likelihood phylogenetic tree of SSF Na+/solute symporters**. The tree was constructed with the LG+C60+Γ4+F substitution model. Branch support was assessed using 1000 ultrafast bootstraps. The scale bar represents the estimated number of substitutions per site.

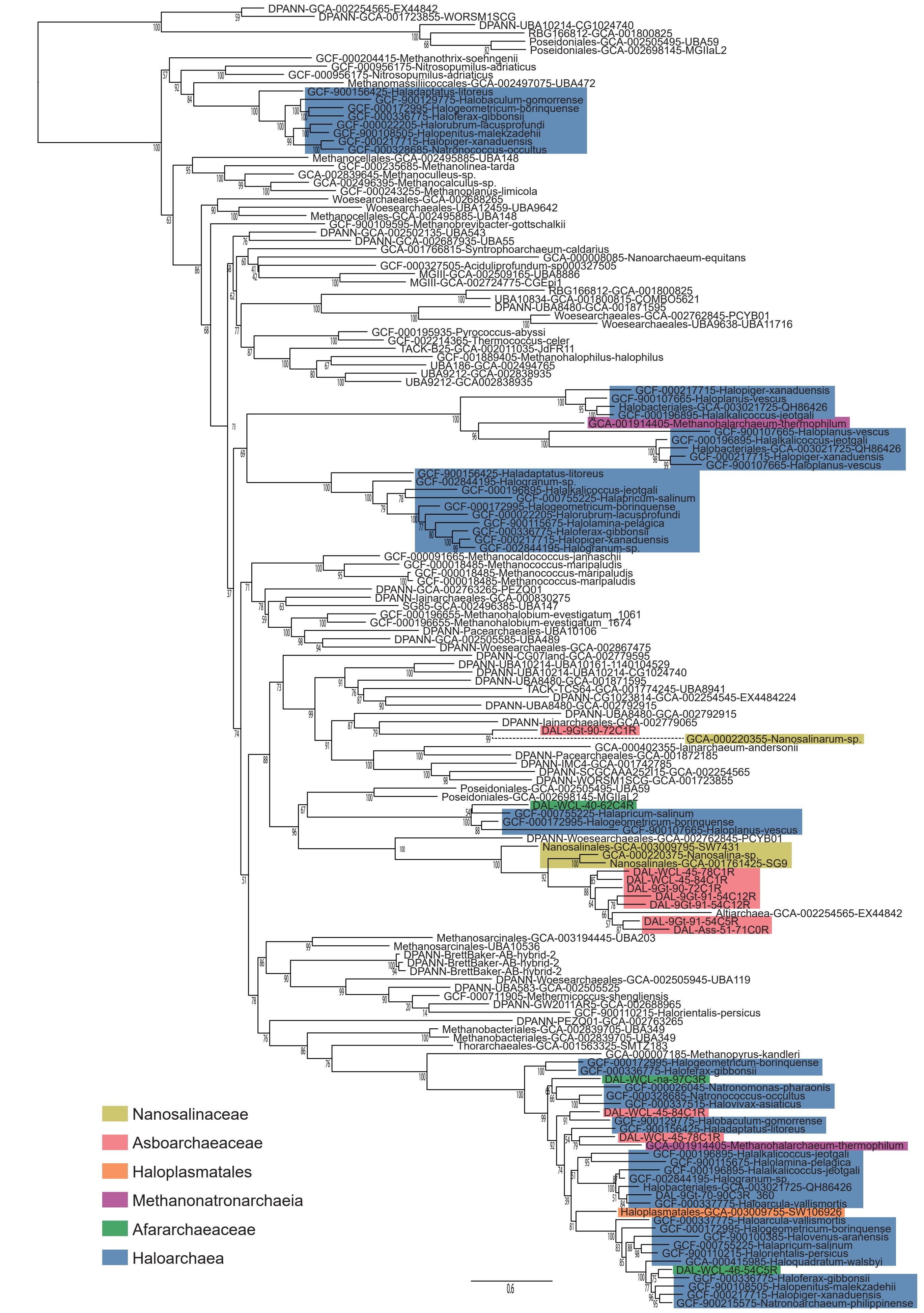


**Supplementary Fig. 22 | Maximum likelihood phylogenetic tree of Ca+/Na+ antiporters**. The tree was constructed with the LG+C60+Γ4+F substitution model. Branch support was assessed using 1000 ultrafast bootstraps. The scale bar represents the estimated number of substitutions per site.

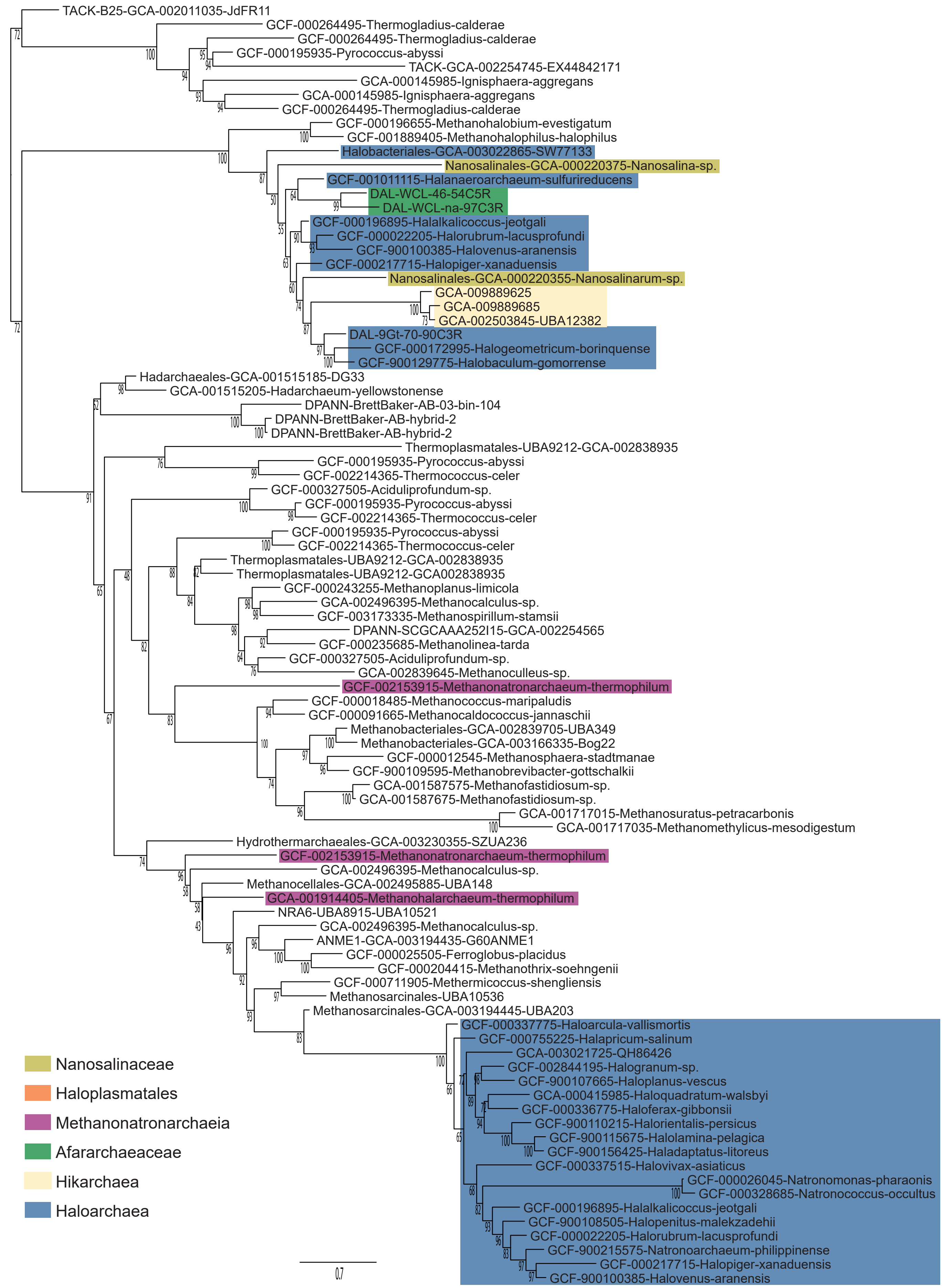


**Supplementary Fig. 23 | Maximum likelihood phylogenetic tree of Na+/H+ antiporters**. The tree was constructed with the LG+C60+Γ4+F substitution model. Branch support was assessed using 1000 ultrafast bootstraps. The scale bar represents the estimated number of substitutions per site.

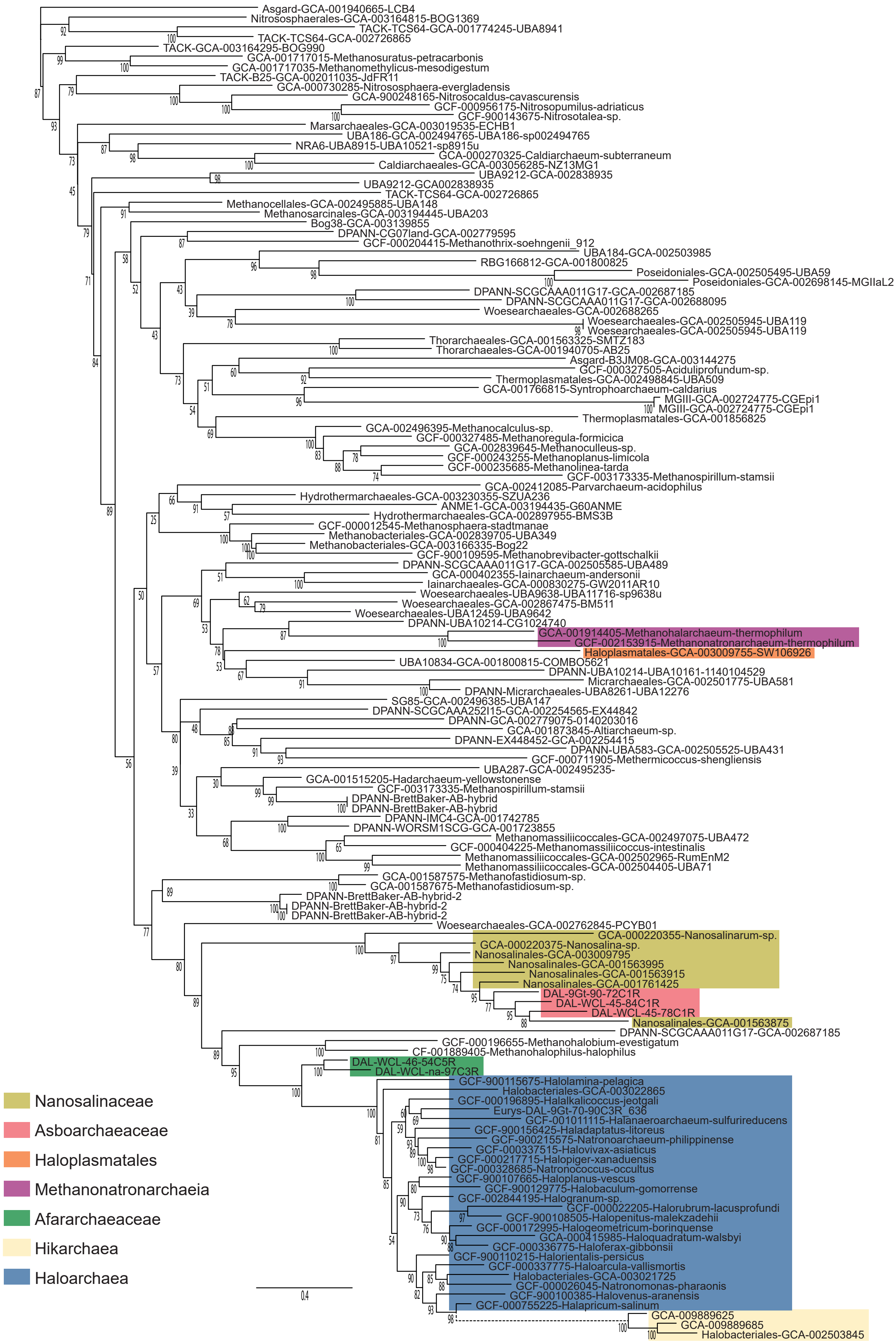


**Supplementary Fig. 24 | Maximum likelihood phylogenetic tree of the chaperone GrpE**. The tree was constructed with the LG+C60+Γ4+F substitution model. Branch support was assessed using 1000 ultrafast bootstraps. The scale bar represents the estimated number of substitutions per site.

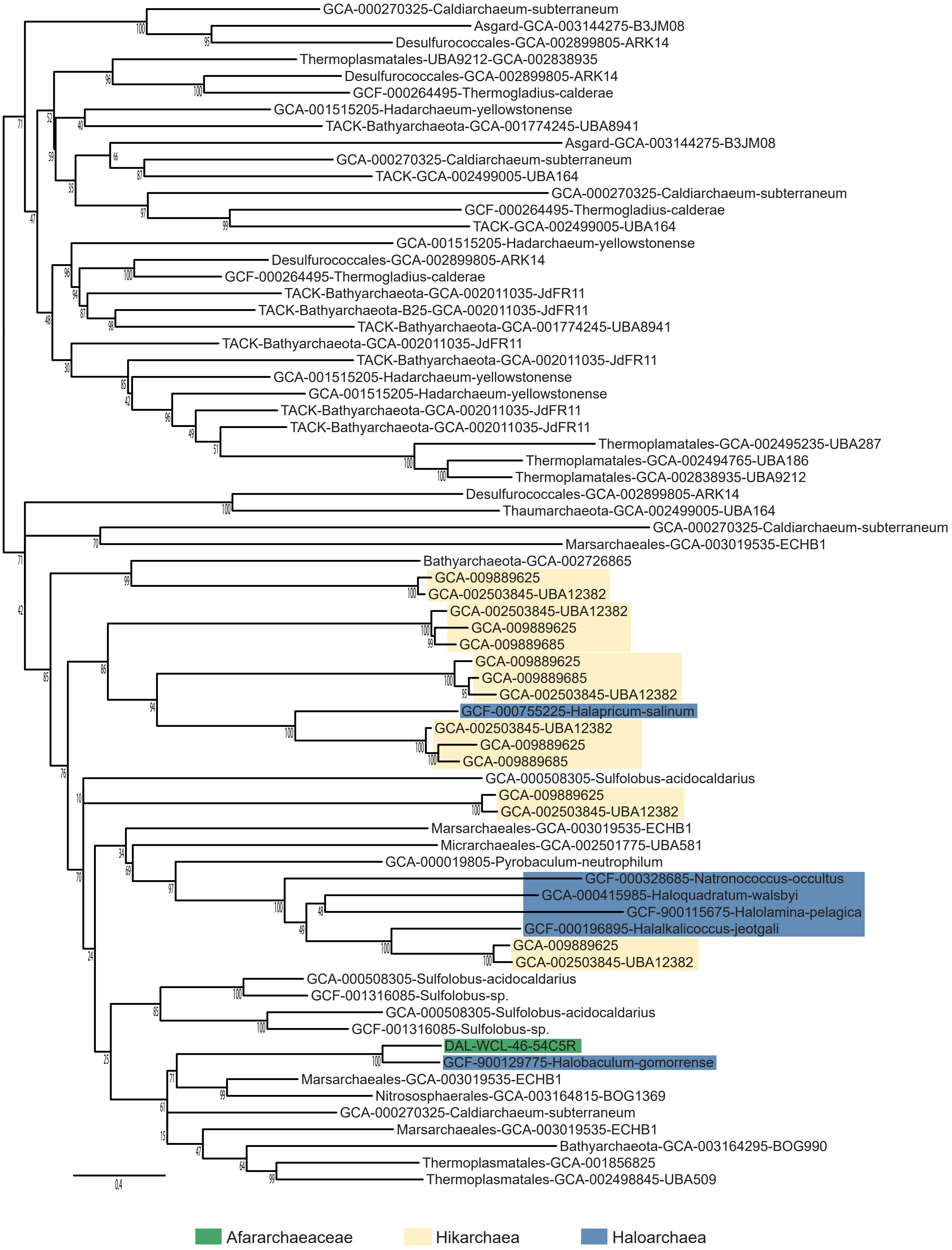


**Supplementary Fig. 25 | Maximum likelihood phylogenetic tree of the aerobic-type carbon monoxide dehydrogenase**. The tree was constructed with the LG+C60+Γ4+F substitution model. Branch support was assessed using 1000 ultrafast bootstraps. The scale bar represents the estimated number of substitutions per site.

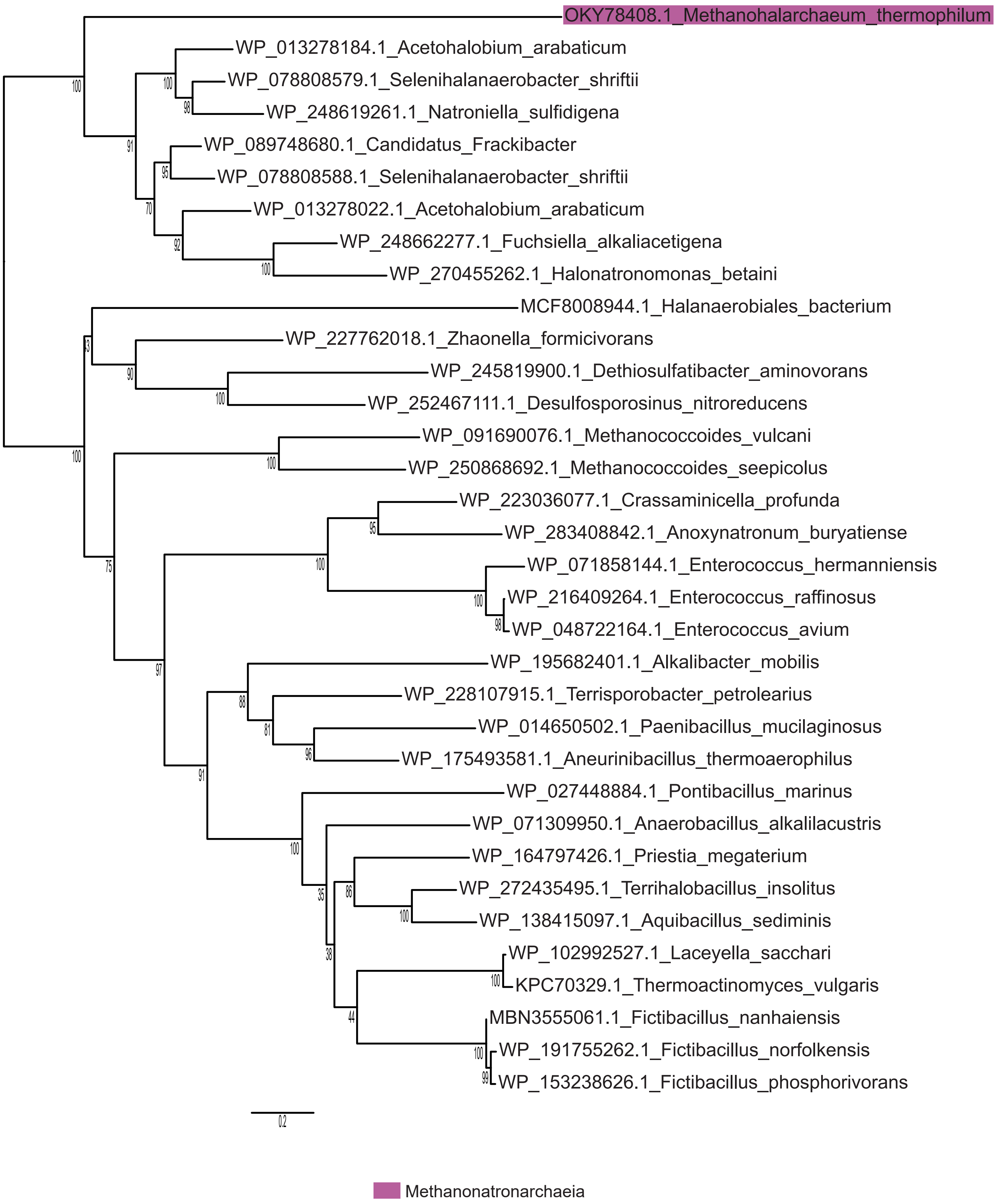


**Supplementary Fig. 26 | Maximum likelihood phylogenetic tree of BCCT transporter**. The tree was constructed with the LG+C60+Γ4+F substitution model. Branch support was assessed using 1000 ultrafast bootstraps. The scale bar represents the estimated number of substitutions per site.

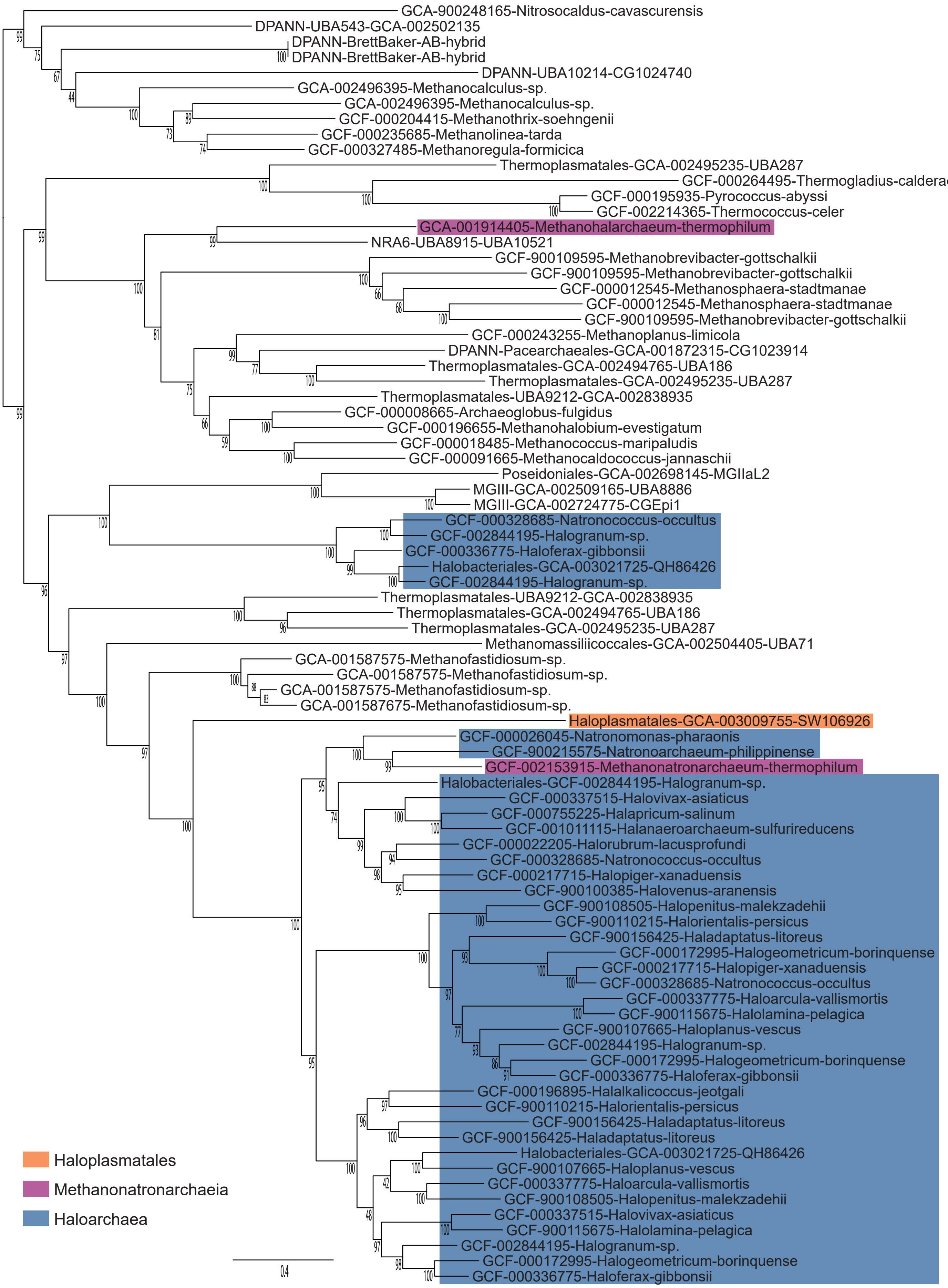


**Supplementary Fig. 27 | Maximum likelihood phylogenetic tree of SNF-family Na+-dependent transporters**. The tree was constructed with the LG+C60+Γ4+F substitution model. Branch support was assessed using 1000 ultrafast bootstraps. The scale bar represents the estimated number of substitutions per site.

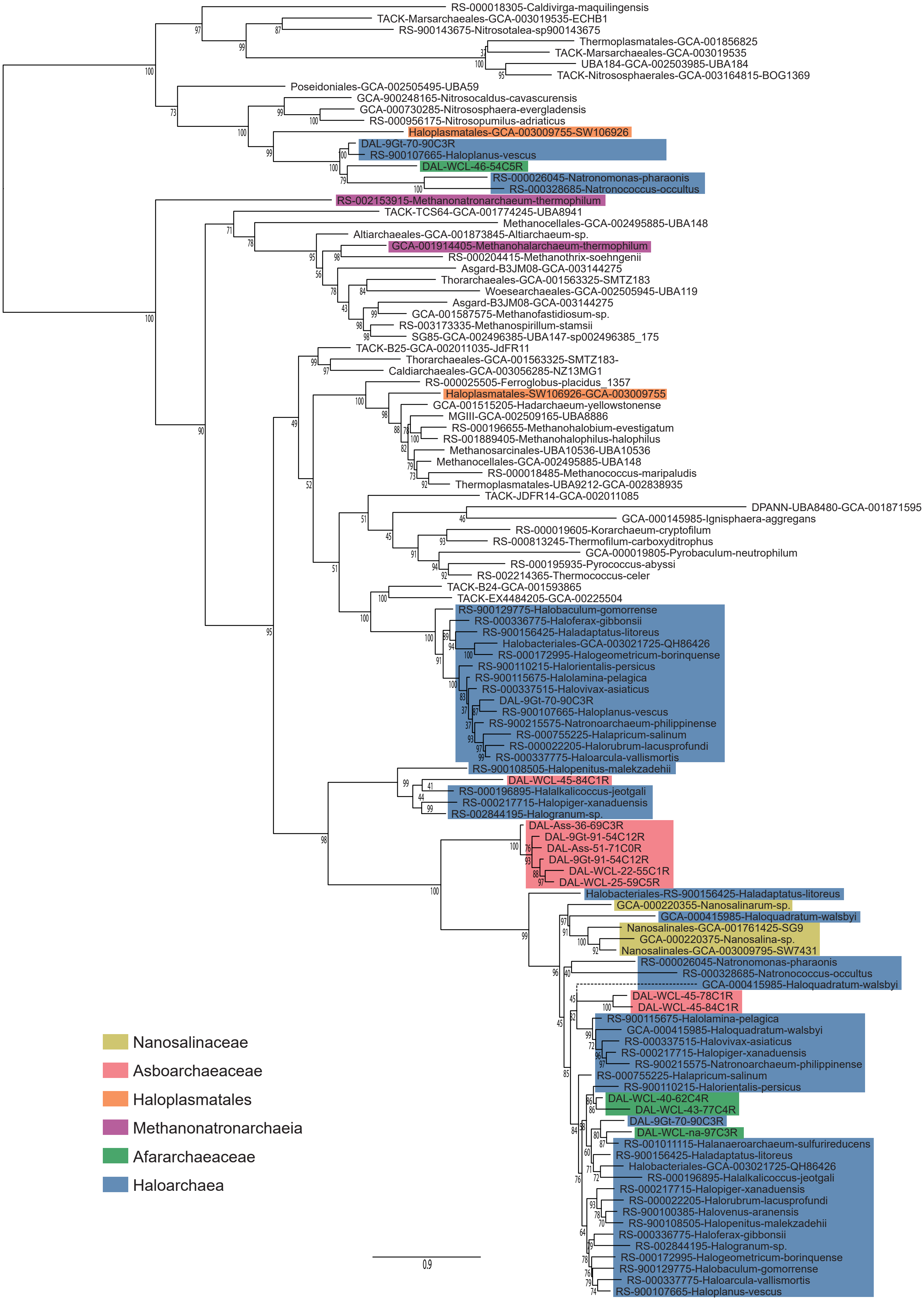


**Supplementary Fig. 28 | Maximum likelihood phylogenetic tree of ZupT metal transporters**. The tree was constructed with the LG+C60+Γ4+F substitution model. Branch support was assessed using 1000 ultrafast bootstraps. The scale bar represents the estimated number of substitutions per site.

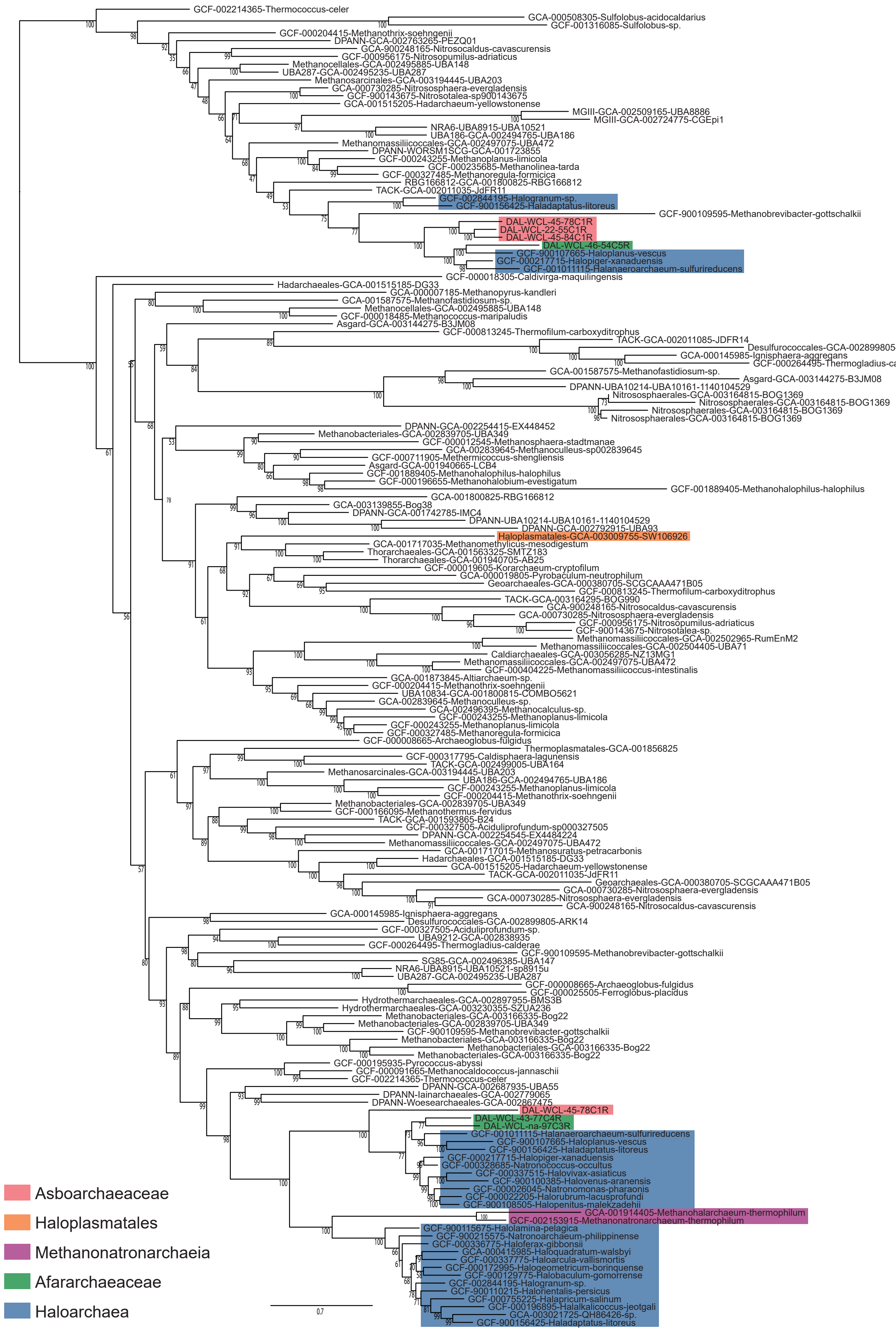


**Supplementary Fig. 29 | Maximum likelihood phylogenetic tree of FieF metal transporters**. The tree was constructed with the LG+C60+Γ4+F substitution model. Branch support was assessed using 1000 ultrafast bootstraps. The scale bar represents the estimated number of substitutions per site.

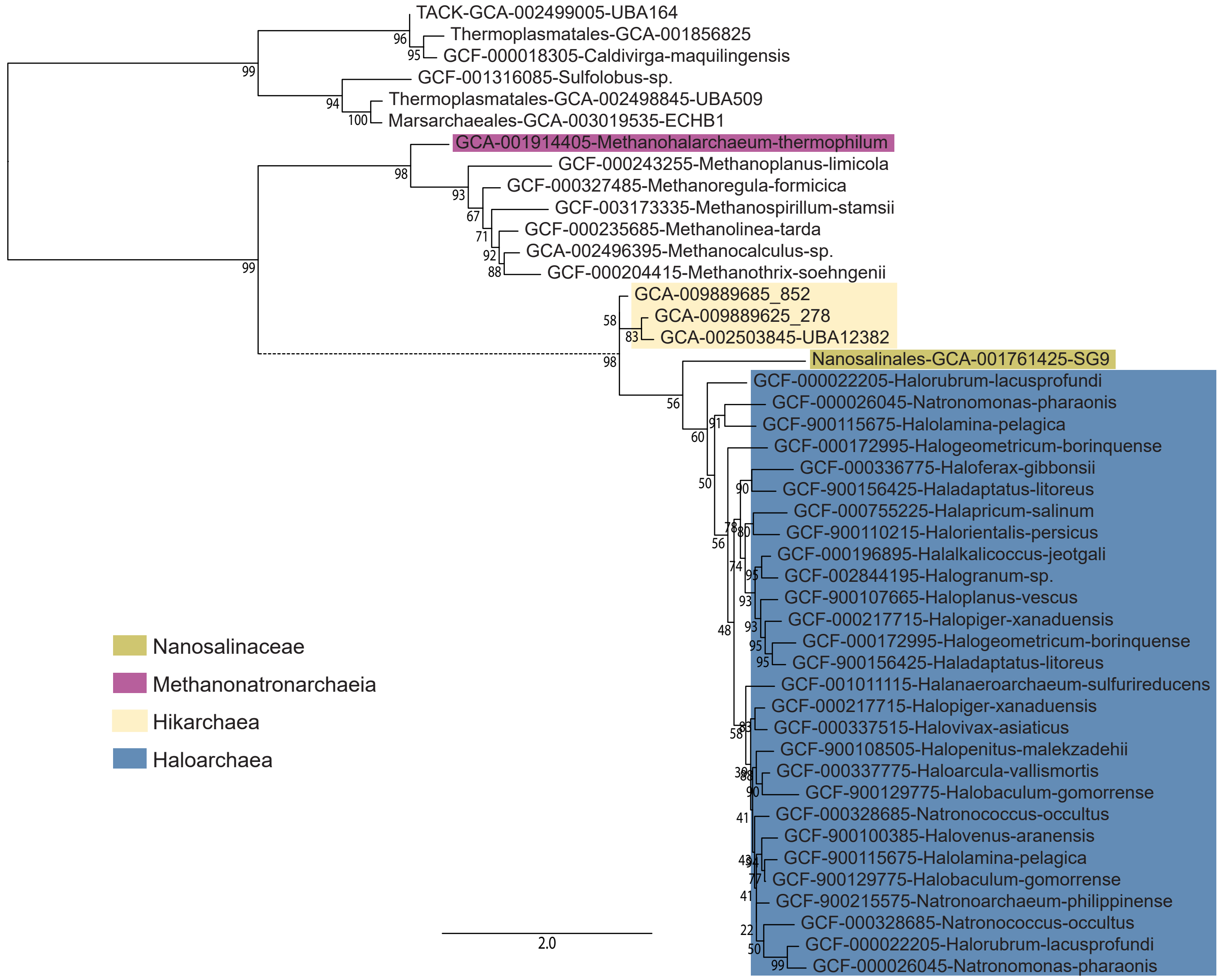


**Supplementary Fig. 30 | Maximum likelihood phylogenetic tree of sulfur transporters**. The tree was constructed with the LG+C60+Γ4+F substitution model. Branch support was assessed using 1000 ultrafast bootstraps. The scale bar represents the estimated number of substitutions per site.

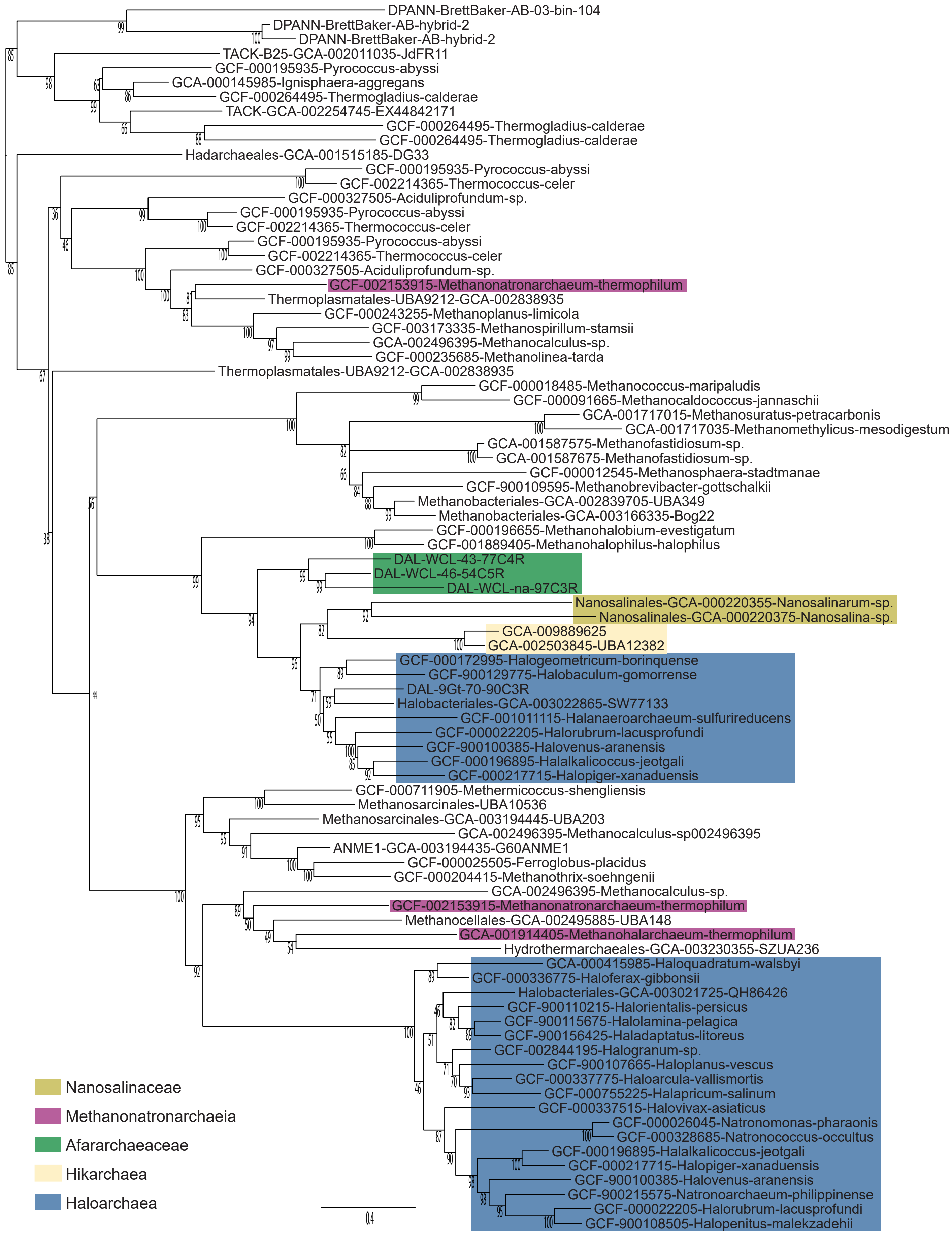


**Supplementary Fig. 31 | Maximum likelihood phylogenetic tree of Na+/H+ antiporters**. The tree was constructed with the LG+C60+Γ4+F substitution model. Branch support was assessed using 1000 ultrafast bootstraps. The scale bar represents the estimated number of substitutions per site.

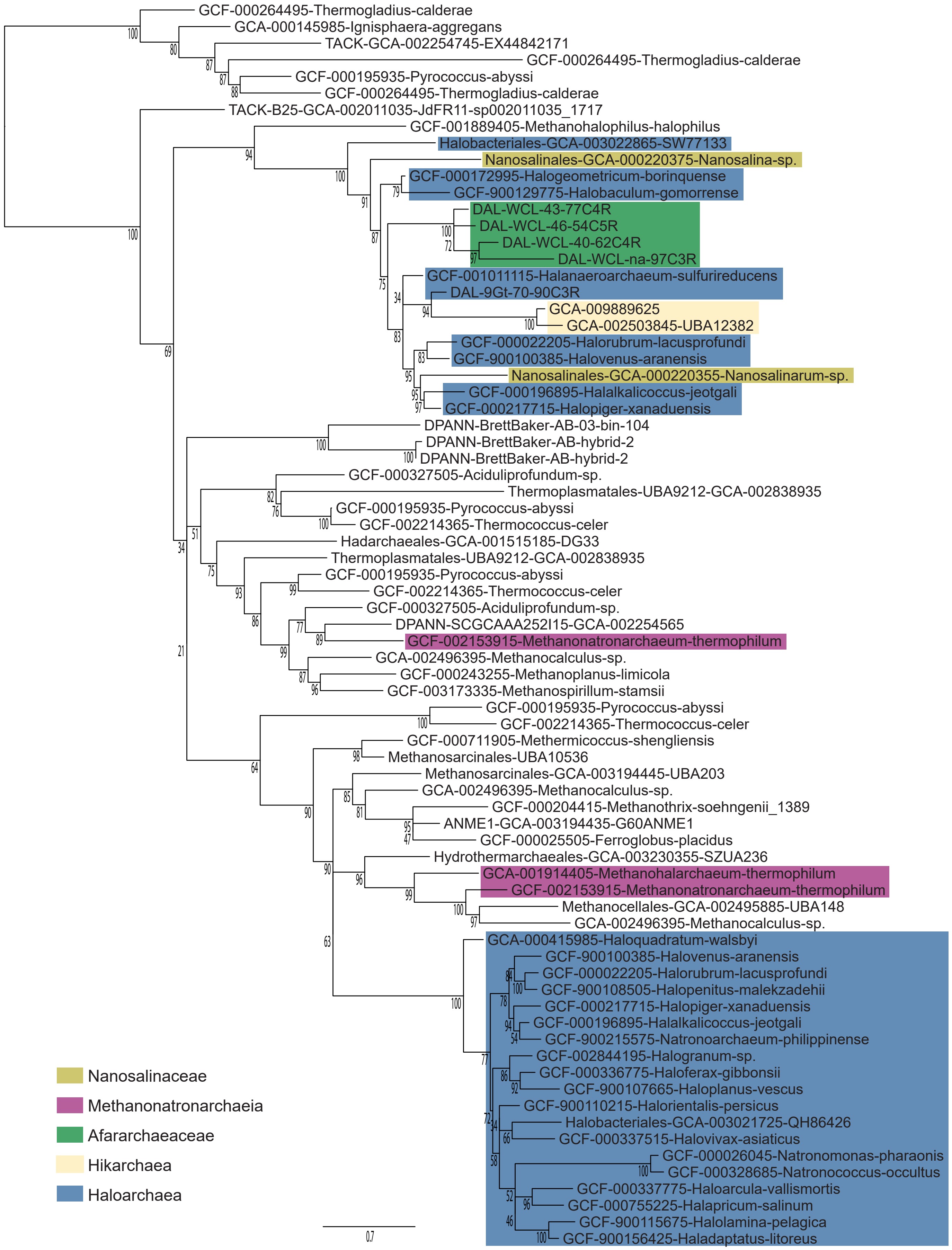


**Supplementary Fig. 32 | Maximum likelihood phylogenetic tree of Na+/H+ antiporters**. The tree was constructed with the LG+C60+Γ4+F substitution model. Branch support was assessed using 1000 ultrafast bootstraps. The scale bar represents the estimated number of substitutions per site.

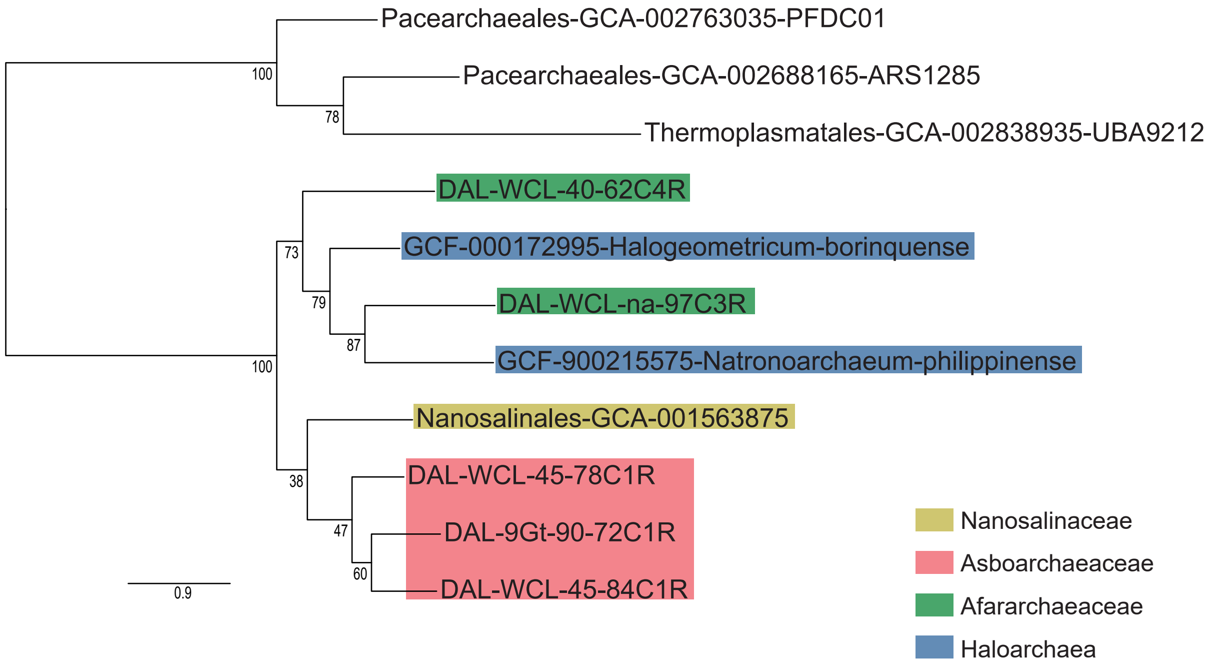


**Supplementary Fig. 33 | Maximum likelihood phylogenetic tree of Na+/phosphate symporters**. The tree was constructed with the LG+C60+Γ4+F substitution model. Branch support was assessed using 1000 ultrafast bootstraps. The scale bar represents the estimated number of substitutions per site.

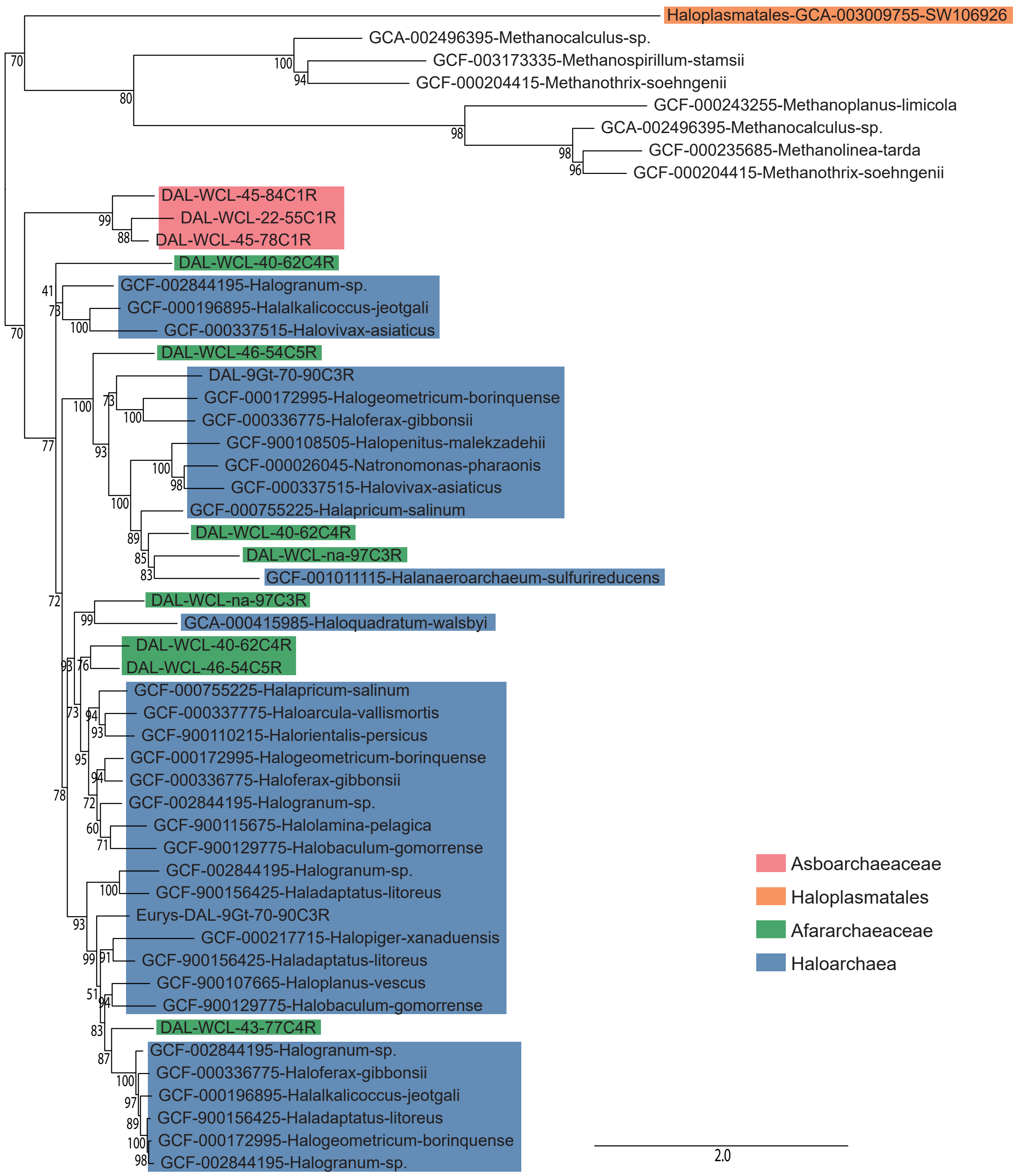


**Supplementary Fig. 34 | Maximum likelihood phylogenetic tree of transporters of di- and tricarboxylate Krebs cycle intermediates**. The tree was constructed with the LG+C60+Γ4+F model of sequence substitution model. Branch support was assessed using 1000 ultrafast bootstraps. The scale bar represents the estimated number of substitutions per site.

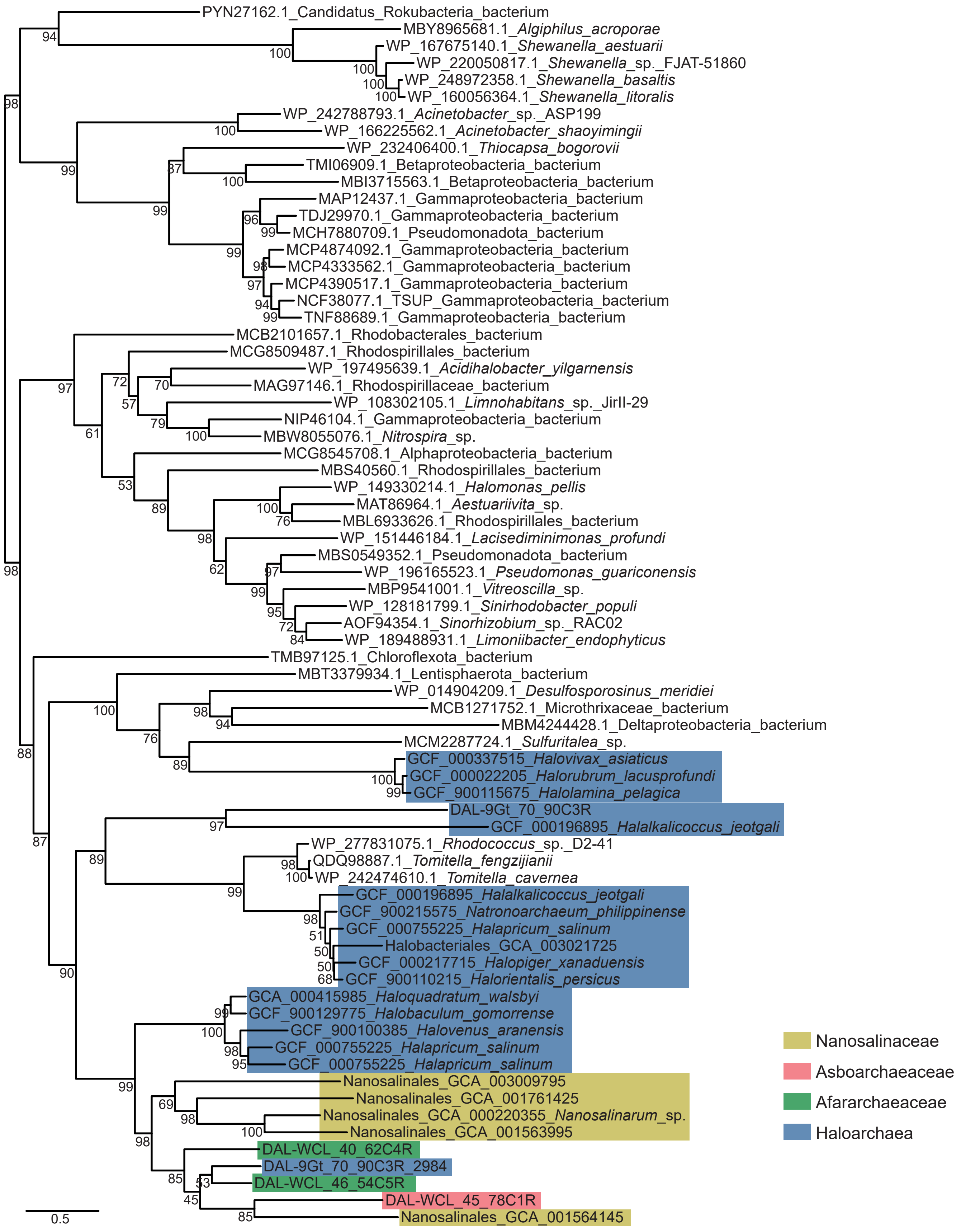


**Supplementary Fig. 35 | Maximum likelihood phylogenetic tree of the AmiS/UreI urea transporter**. The tree was constructed with the LG+C60+Γ4+F substitution model. Branch support was assessed using 1000 ultrafast bootstraps. The scale bar represents the estimated number of substitutions per site.

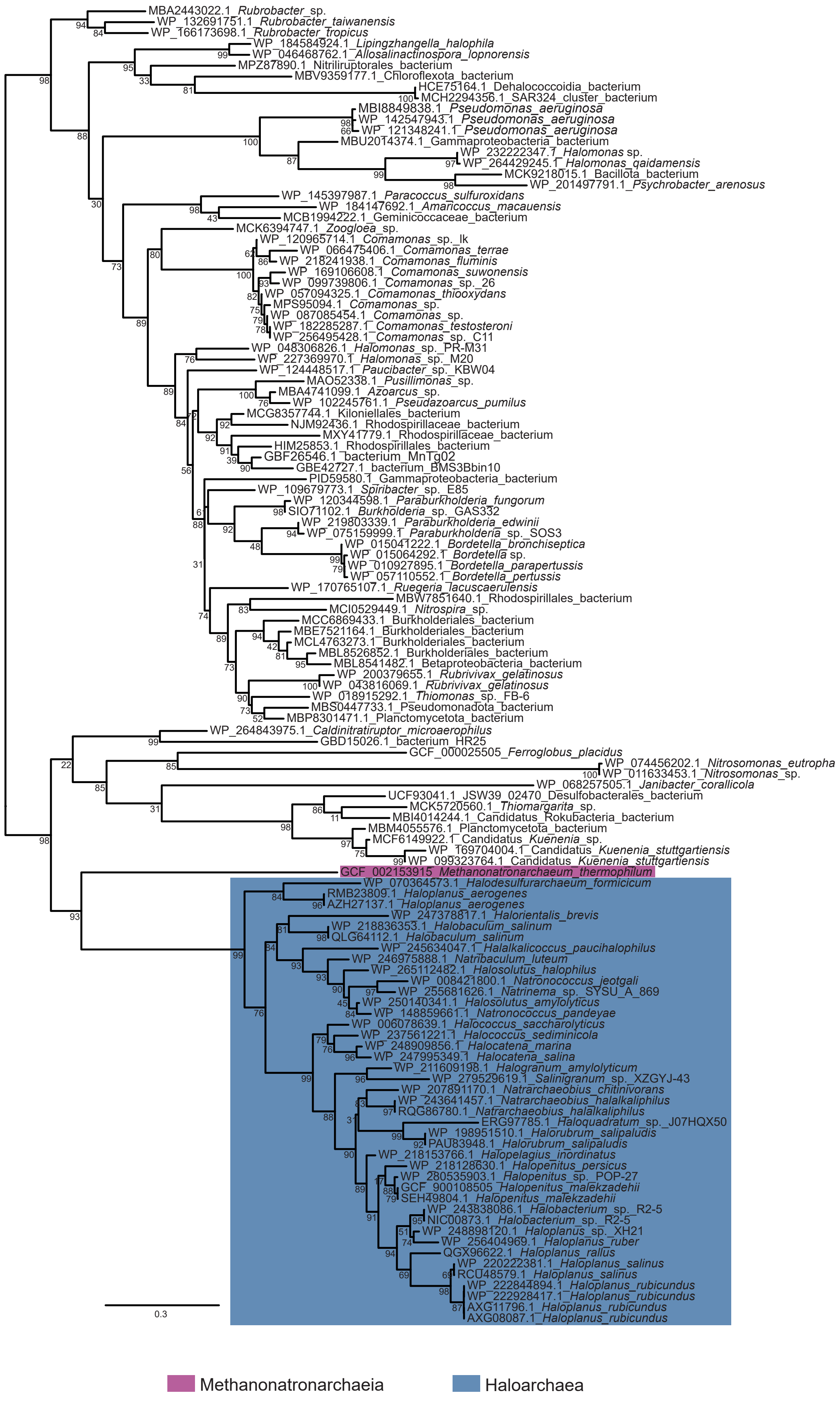


**Supplementary Fig. 36 | Maximum likelihood phylogenetic tree of TauE/SafE sulfite exporter**. The tree was constructed with the LG+C60+Γ4+F substitution model. Branch support was assessed using 1000 ultrafast bootstraps. The scale bar represents the estimated number of substitutions per site.
